# Mathematical Modeling Unveils Optimization Strategies for Targeted Radionuclide Therapy of Blood Cancers

**DOI:** 10.1101/2024.05.22.595377

**Authors:** Maxim Kuznetsov, Vikram Adhikarla, Enrico Caserta, Xiuli Wang, John E. Shively, Flavia Pichiorri, Russell C. Rockne

## Abstract

Targeted radionuclide therapy is based on injections of cancer-specific molecules conjugated with radioactive nuclides. Despite the specificity of this treatment, it is not devoid of side-effects limiting its use and is especially harmful for rapidly proliferating organs well perfused by blood, like bone marrow. Optimization of radioconjugates administration accounting for toxicity constraints can increase treatment efficacy. Based on our experiments on disseminated multiple myeloma mouse model treated by ^225^Ac-DOTA-daratumumab, we developed a mathematical model which investigation highlighted the following principles for optimization of targeted radionuclide therapy. 1) Nuclide to antibody ratio importance. The density of radioconjugates on cancer cells determines the density of radiation energy deposited in them. Low labeling ratio as well as accumulation of unlabeled antibodies and antibodies attached to decay products in the bloodstream can mitigate cancer radiation damage due to excessive occupation of specific receptors by antibodies devoid of radioactive nuclides. 2) Cancer binding capacity-based dosing. The rate of binding of drug to cancer cells depends on the total number of their specific receptors, which therefore can be estimated from the pharmacokinetic curve of diagnostic radioconjugates. Injection of doses significantly exceeding cancer binding capacity should be avoided since radioconjugates remaining in the bloodstream have negligible efficacy to toxicity ratio. 3) Particle range-guided multi-dosing. The use of short-range particle emitters and high-affinity antibodies allows for robust treatment optimization via initial saturation of cancer binding capacity, enabling redistribution of further injected radioconjugates and deposited dose towards still viable cells that continue expressing specific receptors.

**Significance:** Mathematical modeling yields general principles for optimization of targeted radionuclide therapy in mouse models of multiple myeloma that can be extrapolated on another cancer models and on clinical setting.

## Introduction

### Biological background

Targeted therapy is based on suppressing survival and proliferation of cancer cells through interactions with molecules specific to the cancer type under consideration. Large class of targeted agents are monoclonal antibodies that specifically bind to receptors on the surfaces of cancer cells [1]. One such example is daratumumab, which targets CD38. These receptors are overexpressed in multiple myeloma, which is a white blood cell cancer resulting in about a hundred thousand deaths worldwide annually [2]. The main mode of action of daratumumab is induction of cancer cell killing by the immune system [3]. Clinical trials have shown favorable safety profile of daratumumab [4]. However, its action results in highly heterogeneous outcomes, including frequent cancer relapse after initial response. Relevant studies suggest that the mechanisms of multiple myeloma resistance to daratumumab are associated with immune escape [5], as relapsing patients show stable expression of unmutated CD38 on cancer cells, which preserves their ability to be targeted by daratumumab [6].

A way to enhance the efficacy of targeted therapy is to attach additional therapeutic payload to antibodies, ensuring its selective delivery to malignant cells. Conjugation of antibodies with cytotoxic agents has already led to more than a dozen clinically approved drugs [1], and conjugation of antibodies with radioactive nuclides is gaining growing clinical interest [7]. Two main types of radionuclides are being investigated in trials: emitters of *β*-particles (electrons), which deposit comparably low energy over long distance, and emitters of *α*-particles (two protons and two neutrons bound together), which deposit higher energy over short range, corresponding to only a few cell diameters.

Toxicity of targeted radionuclide therapy (TRT) is a more complicated matter compared to that of external beam radiotherapy [8]. Radioconjugates attach to specific receptors, expressed in some types of healthy cells, and thus affect them. Clearance of a part of radionuclides through liver and kidneys is toxic for these organs. Radionuclides circulating in the bloodstream pose a significant threat to well-perfused rapidly proliferating organs such as bone marrow, and thus suppress haematopoiesis. Therefore, bone marrow is often regarded as the dose-limiting organ for TRT [9].

In light of stable expression of CD38 in multiple myeloma cells, it is encouraging to use daratumumab to guide delivery of radionuclides to them. Previously we have compared the efficacy and toxicity of *α*-emitter ^225^Ac-DOTA-daratumumab and *β*-emitter ^177^Lu-DOTA-daratumumab in a mouse model of disseminated multiple myeloma. We concluded that the actinium-based *α*-emitter shows more promise for clinical translation [10], and now we are conducting the phase I clinical trial for assessing its safety for patients (NCT#05363111).

Although radiolabeling of daratumumab increases its efficacy [11, 12], achieving long-term response represents an extremely challenging task. Formally, cancer cure implies elimination of every single clonogenic malignant cell, while even a technically undetectable residual number of them can promote cancer recurrence [13]. Treatment efficacy can be compromised not only by dose-limiting side effects, but also by inherent restrictions in cancer cells exposure to drug. The choice of proper therapy protocol is further complicated by inter-patient variability [7]. Mathematical modeling provides a tool to help navigate this complexity and facilitate treatment optimization.

### Mathematical background

In this work we focus on mechanistic mathematical modeling, which implies representation of cancer, its environment and treatment as a system of equations, which terms correspond to specific natural laws [14]. This approach allows formulating the tasks of treatment optimization as optimal control problems, which can be solved analytically or numerically. Quantitative agreement of modeling predictions with experimental data cannot be accurate on individual level due to significant heterogeneity of biological objects and unavoidable uncertainly in individual quantitative characteristics. This issue can be addressed with *in silico* trials simulating treatment outcomes for heterogeneous virtual populations [15]. Robust conclusions gained by mathematical modeling can be verified in clinical trials and eventually be implemented into clinical decision-making pipeline [16].

Mathematical modeling of TRT relies on modeling of drug pharmacokinetics, which is an extensive research area [17], and on modeling of continuous exposure of cells to irradiation, which has a well-established mathematical foundation [18]. Yet, to date there exist only a few studies on mathematical modeling of TRT. The works by Kletting et al. present detailed pharmacokinetic models, designed with the goal of predicting the biologically effective doses received by tumors and healthy organs [19, 20] as well as the tumor response [21] for varying amounts of injected *β*-emitters. These works consider data of patients with prostate cancer, neuroendocrine cancer and meningioma. Other works of this and other research groups focus on constructing mathematical models using data from mouse tumor experiments, which include xenografted neuroendocrine tumor model treated by *α*-emitting ^212^Pb-DOTAMTATE [22], thyroid cancer xenograft model treated by *α*-emitting ^212^At which naturally accumulates in thyroid [23], and transgenic murine model of metastatic breast cancer treated by *α*-emitters ^225^Ac and ^213^Bi, conjugated with anti-HER2/neu antibodies [24]. To the best of our knowledge, the latter work is the only study providing simulations of multi-dose TRT. Its results suggest that dose fractionation can affect mice survival, however, an explicit optimization problem is not considered.

Previously, based on our experiments on disseminated multiple myeloma mouse model and its treatment by ^225^Ac-DOTA-daratumumab, we have developed the first mathematical model of TRT of blood cancer [10]. Further we incorporated the experimental data on CAR T cell treatment within it in order to provide ground for optimization of scheduling of such combination therapy [25, 26]. The current study is a continuation of this research, aimed at suggesting ways for optimization of TRT in single-dose and multi-dose setting.

For the sake of conciseness of this paper, a significant part of mathematical reasoning and simulation results are provided in supplementary material [27–30]. All the computational codes were implemented in Wolfram Mathematica software.

## Materials and Methods

The presented mathematical model is parametrized using the data from our preclinical studies investigating the therapeutic efficacy and toxicity of ^225^Ac-DOTA-daratumumab in disseminated human MM.1S xenograft mouse model, reported previously [10]. The details of animal studies are summarized in supplementary Section S.1.1. The mathematical model is supplemented by relevant literature data for quantifying aspects not directly measured in our experiments.

For brevity, radioconjugates with yet undecayed radionuclides are referred to as *active antibodies*. Antibodies conjugated with final decay products and unlabeled antibodies are together termed *inert antibodies*. Likewise, receptors bound to active/inert antibodies are denoted as *active/inert receptors*. Radionuclides attached/unattached to cancer cells are referred to as *anchored/unanchored nuclides*. The schematic view of the model is shown in **Fig. 1**. The model is built on the following assumptions.

**Figure 1:**
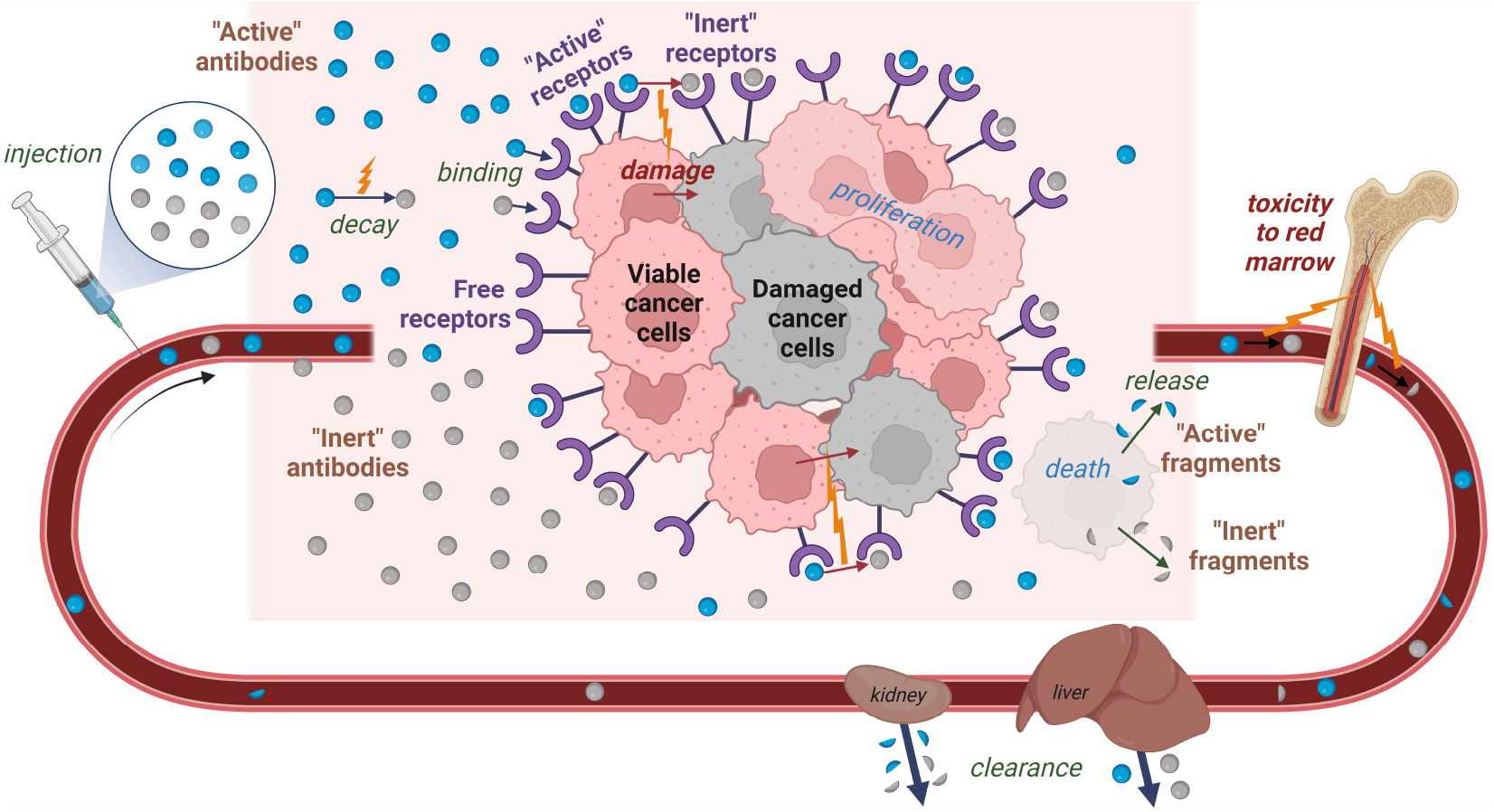
Schematic view of the main processes considered in the mathematical model. Created with BioRender.

### One-step nuclear reaction is considered

Active antibodies transform directly into inert antibodies upon radionuclide decay. Such approximation is justified for ^225^Ac, which half-life significantly exceeds the half-lives of its daughter nuclides [31].

### Antibody-receptor binding is irreversible

We have shown previously that daratumumab is rapidly internalized by multiple myeloma cells [32]. For simplicity, we do not consider internalized antibodies as a separate variable and assume sufficiently high affinity of antibodies that allows neglecting their unbinding.

### Blood plasma and cancer cells microenvironment represent a well-mixed medium

Multiple myeloma cells are predominately confined to bone marrow. The rapid exchange of substances in it takes place through the discontinuous endothelium of venous sinuses. Experimental data suggest that equilibration of intravenously administered high molecular weight substances in blood plasma and bone marrow is achieved within only minutes [33].

### Cancer cell population is homogeneous

Viable cells proliferate with constant rate. This process can be ceased by their irradiation damage, with cancer cell having equal radiosensitivity. Damaged cells immediately stop proliferating and die at a constant rate. All cancer cells are reachable by antibodies and have an equal number of specific receptors.

### Damage of cancer cells is irrepairable, with its rate being proportional to the delivered dose

This approach is justified for modeling of the effect of *α*-particles manifested in DNA double-strand breaks [34].

### Antibodies are degraded upon cancer cell death and are released back in bloodstream in fragmented form

Therefore, some of the radionuclides return back to the bloodstream being attached to the fragments of antibodies. The clearance rate of these fragments is higher than that of intact antibodies.

### Only decays of nuclides unattached to cancer cells are toxic to normal cells

Decays of unanchored nuclides are attributed to treatment toxicity, since in general a part of them leads to the damage of red marrow, implied as the crucial organ at risk. We assume that the particles emitted from nuclides anchored to cancer cells effectively deposit all their energy in them, resulting in negligible toxicity to normal cells.

### Antibodies do not elicit therapeutic action, other than that due to the radionuclides

Our experiments involved immunodeficient mice, with cancer dynamics not affected by unlabeled daratumumab.

### Conjugation of nuclides to antibodies and their decays do not alter antibody biodistribution

Therefore, active and inert antibodies have equal rates of binding and equal rates of clearance.

### Antibodies do not bind to normal cells

Our experiments used human cancer xenografts in mice and human antibodies, unable to attach to receptors of mouse cells. Immunoglobulin was preinjected to suppress antibodies recycling.

The following equations govern the model dynamics. All the used variables and parameters values are non-negative. The injection terms represent the external control of the otherwise autonomous system of equations. Initially, the number of viable cancer cells is *N* (0) = *N*_0_; damaged cancer cells are absent: *D*(*t*) = 0; all receptors are free: *f*_*F N*_ (0) = *f*_*F D*_(0) = 1, *f*_*AN*_ (0) = *f*_*AD*_(0) = 0; and antibodies are absent: *a*(0) = *b*(0) = *p*(0) = 0.

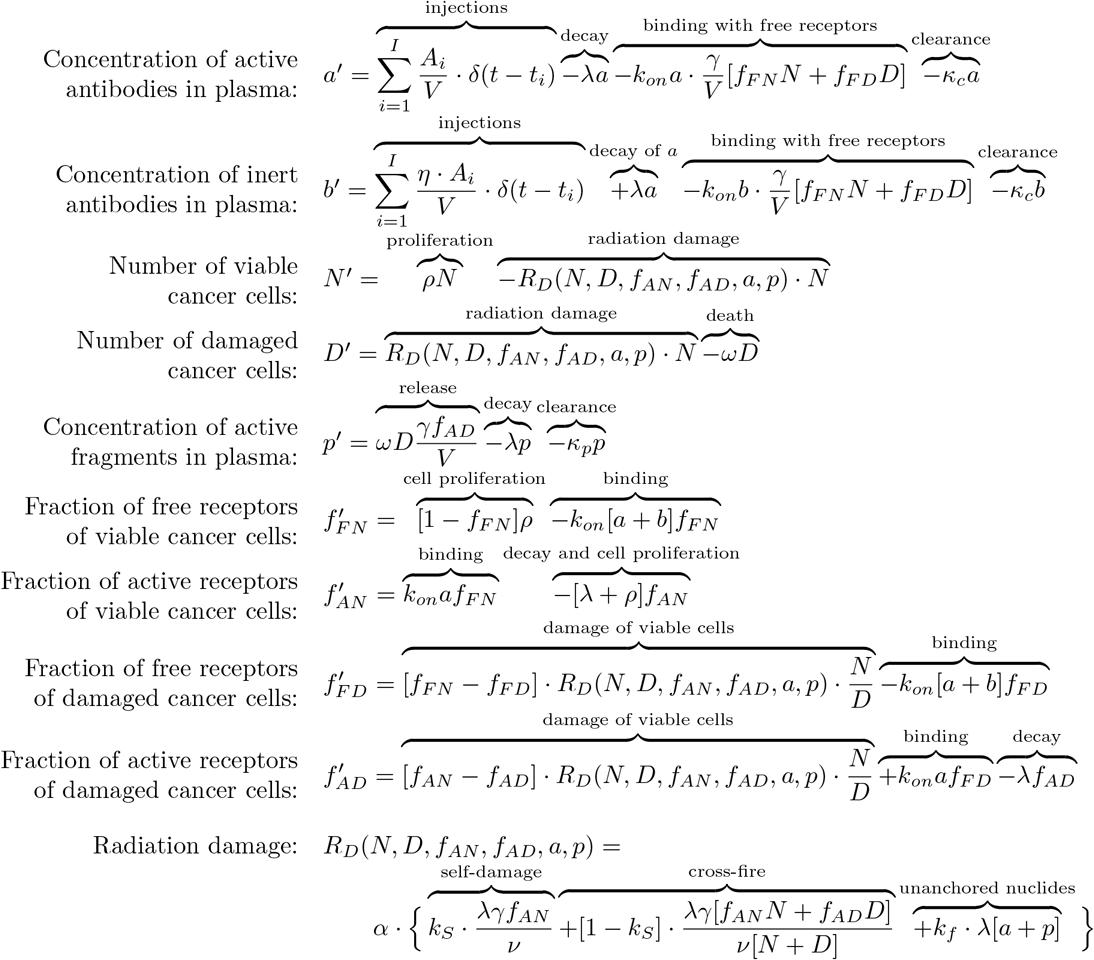

Injections of active antibodies result in instantaneous increase of their plasma concentration. Simultaneous injections of inert antibodies can be accounted for, with *η* taking positive values for such cases. This reflects the presence of impurities that accompany the production of radioconjugates, which are unlabeled antibodies and antibodies conjugated with non-radioactive ions [35]. This can also reflect deliberate dilution of drug within unlabeled antibodies.

The function of radiation damage accounts for three sources of radiation that can affect viable cancer cells. The terms of self-damage and cross-fire stand for the damage of a cancer cell due to the decays taking place on its own receptors and on the receptors of neighboring cells, respectively. The numerator in each of these terms denotes the rate of decays within a certain volume, provided in the denominator. The relative significance of self-damage *k*_*s*_ for *α*-particles in dense cancer tissue can be small, since their range is generally greater than a cell diameter and their ionization density increases towards its end [36]. The value of *k*_*s*_ should, however, increase with the decrease of cancer cells density. The third term of radiation damage corresponds to decays of unanchored nuclides. For brevity, we will refer to them as *decays in blood*.

Derivation of equations for receptors is performed in supplementary Section S.1.2. Section S.1.3 describes the process of estimation of model parameters and fitting of experimental data. The fitting process suggested significant heterogeneity of cancer cell response to treatment, in particular manifested in the decrease of cancer cell radiosensitivity under the increase of injected dose. Nevertheless, the presented model holds value for theoretical investigation, which is performed below. An augmentation of the model with account of cancer cell heterogeneity is also introduced during this study for verification and generalization of obtained qualitative conclusions.

Table 1 lists the model parameters, formalizes their meaning and provides their basic values as well as the ranges of their variation during parameter sweep. The following normalization parameters are used: *day* for time, *ml* for volume, *pmol* for amounts of antibodies and receptors, *nM* = *pmol/ml* for their concentrations.

**Table 1:**
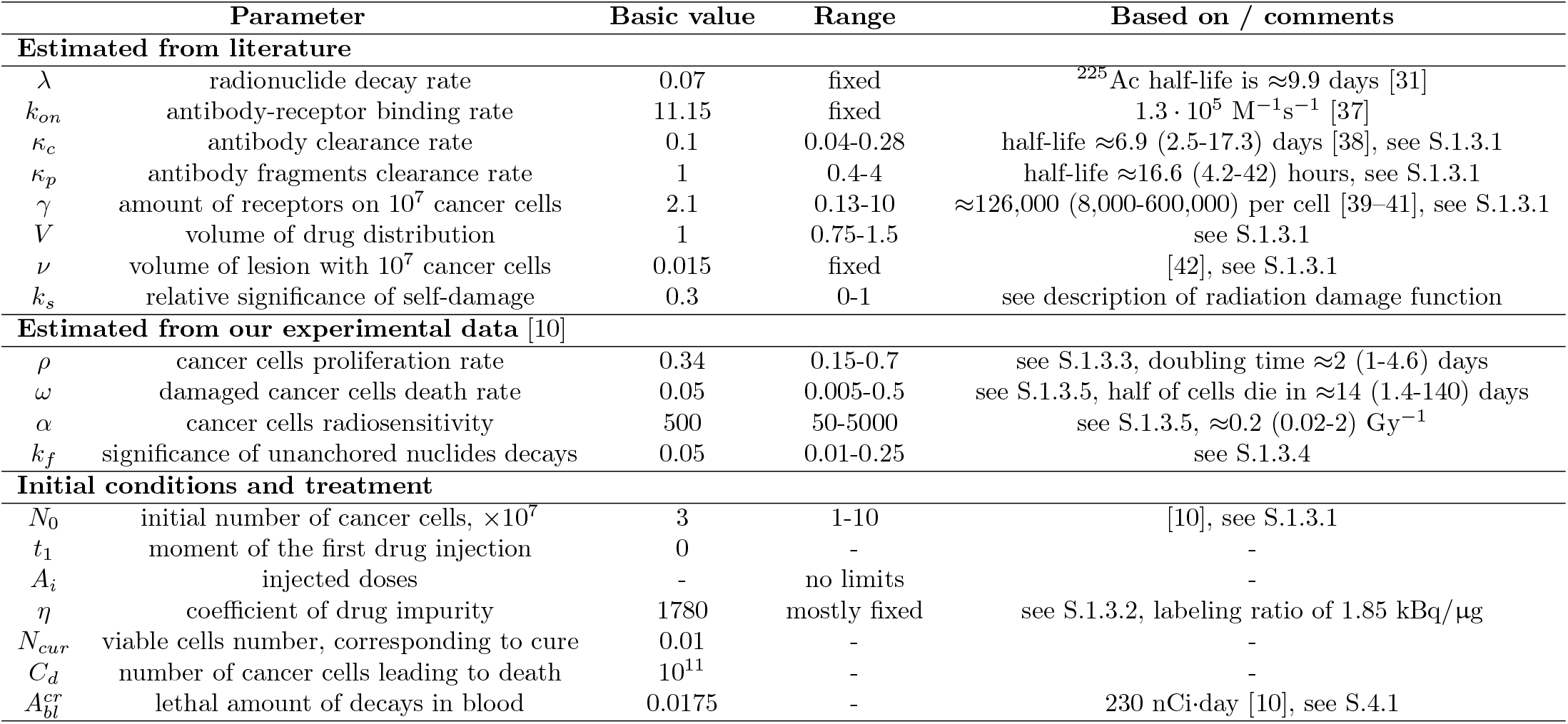
Model parameters.

## Results

### Single-dose treatment by pure radioconjugates

In this section we keep the coefficient of drug impurity *η* = 0. Thus, no inert antibodies are injected simultaneously with active antibodies. Understanding the model behavior in this setting serves as an important steppingstone towards further investigation of the model with account of drug impurities in single-dose and multi-dose settings.

If the amount of injected radioconjugates is notably lower than the amount of free receptors on cancer cells, then the binding of drug to receptors is effectively performed within several hours, as illustrated in **Fig 2A**, and irradiation from unanchored nuclides plays a minor role in the overall damage of cancer cells. These features facilitate analytical investigation of the model, which is performed in supplementary Section S.2.1. The global parameter sweep, performed in Section S.2.2, confirms its validity.

**Figure 2:**
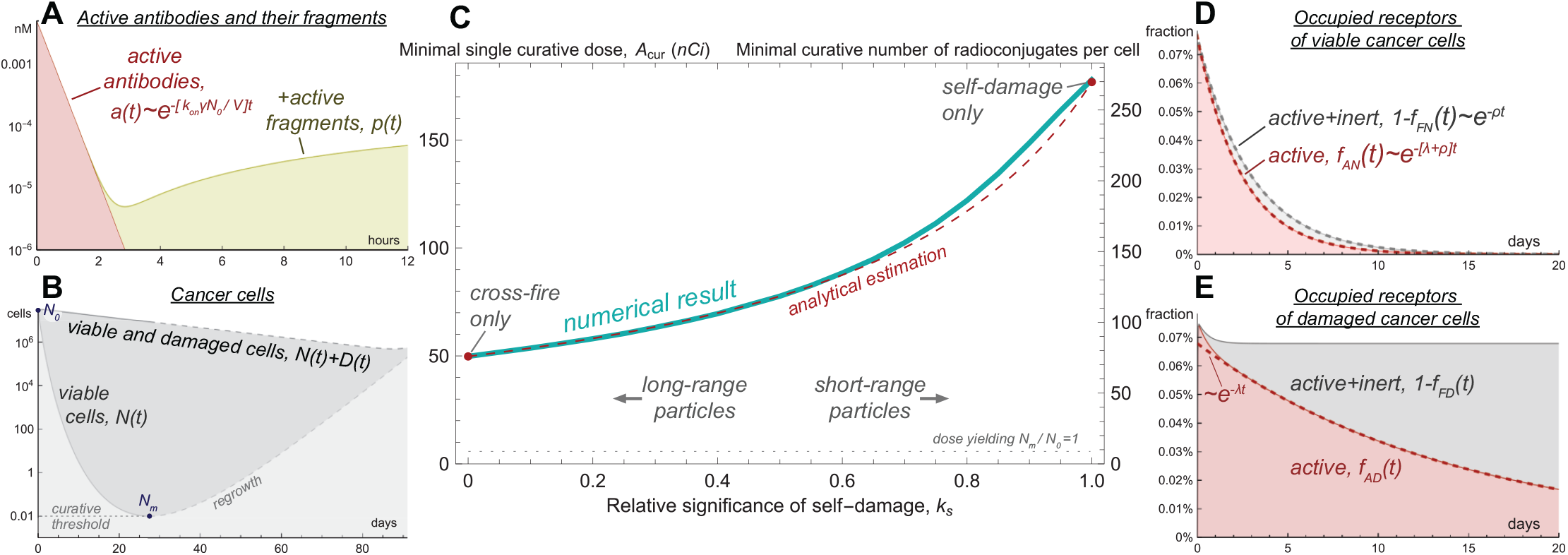
Model simulations of treatment by pure radioconjugates (*η* = 0) for the basic set of parameters. **A**, dynamics of active antibodies and their fragments in plasma. **B**, dynamics of cancer cells. **C**, dependence of minimal single curative dose on relative significance of self-damage, *k*_*s*_. **D**, dynamics of occupied receptors on viable cells. **E**, dynamics of occupied receptors on damaged cells.

**Figure 2B** shows an example of cancer cell dynamics under prolonged irradiation. The minimal number of viable cancer cells *N*_*m*_ is achieved when the rate of nuclear decays falls to the level at which they are not able to compensate for ongoing cell proliferation. The ratio *N*_*m*_*/N*_0_ can be regarded as *minimal surviving fraction* of cancer cells. For injected dose *A* it can be estimated as follows for self-damage-only and cross-fire-only settings:

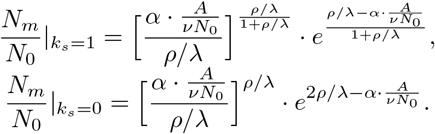

In the limit of *λ → ∞* (immediate nuclide decay) or *ρ →* 0 (absence of cancer growth) analytical estimation of minimal surviving fraction in both settings tends to 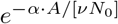. This expression corresponds to the classical formula for surviving fraction of cells after instantaneous irradiation, widely used in external beam setting, since its exponent corresponds to the energy deposited per unit of cancer mass. With the increase of cancer cell proliferation rate the minimal surviving fraction grows, highlighting the negative influence of cancer cell repopulation on the treatment outcome. These formulas are accurate unless the injected dose is sufficiently small or the decay rate of nuclides is so fast that a large fraction of them decays in the bloodstream before anchoring to cancer cells (see supplementary Fig. S.6).

Given the continuous nature of our modeling approach, the numbers of cells are expressed as real numbers. We regard the cases, in which viable cancer cells number falls down to *N*_*m*_ *< N*_*cur*_ = 0.01 cell, as curative. Such approach allows introducing *minimal single curative dose A*_*cur*_ as the dose, resulting in *N*_*m*_ = *N*_*cur*_. **Figure 2C** illustrates the influence of particle range on *A*_*cur*_, emphasizing the qualitative difference of extreme opposite cases, in which irradiation of cells is provided mainly by either self-damage or cross-fire. Self-damage-only case demands greater dose for cure, since the density of radionuclides on viable cells falls not only due to decay of nuclides, but also due to their redistribution among the newborn cells. Therefore, as **Fig. 2D** shows, after binding of antibodies the fraction of active receptors of viable cells decreases as 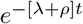. **Figure 2E** shows that the fraction of occupied receptors of damaged cells initially quickly decreases due to the transition of viable cells into damaged state, then it falls only due to decay of nuclides, decreasing as *e*^*−λt*^.

The ratio *A*_*cur*_*/N*_0_ can be regarded as the minimal activity of radioconjugates, initially residing on each cancer cell, that eventually will lead to cure. It can be converted into the *minimal curative number of radioconjugates per cell*. For the depicted variation of particle range it spans from 76 to 272, which is significantly smaller than the average number of CD38 receptors on a MM.1S cell, estimated as *≈*126,000 [39]. However, in practice, the binding of radioconjugates to cancer cells is accompanied by binding of unlabeled antibodies, which are inevitable impurities of the drug and previously injected antibodies remaining in the bloodstream. This highlights the importance of *nuclide to antibody ratio* for achieving the curative density of radioconjugates on cancer cells.

The fraction of activity spent via decays in blood in considered curative setting can be estimated as

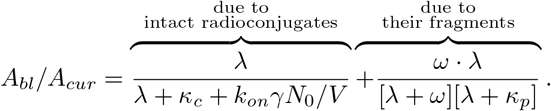

In the used parameter range, *≈*97% of decays in blood are due to the fragments of radioconjugates, released back to the bloodstream from dead cancer cells.

The majority of radiation released in cancer is deposited in already damaged cells and thus formally does not contribute to the overall therapeutic effect. During the parameter sweep the fraction of injected activity spent on viable cancer cells varies in the range 0.4%-3.3%. Analytical estimations show that in the single-dose curative setting this measure cannot exceed 1*/ln*(*N*_0_*/N*_*cur*_), which is *<*5% if the initial number of cancer cells is greater than 10^7^.

Ongoing cancer cell proliferation hampers treatment efficacy. On the other hand, newborn cells express specific receptors, increasing the cancer capacity for drug binding. During the parameter sweep, the number of cancer cells born during treatment by minimal single curative doses constitutes 3-25% of their initial number for the vast majority of cases, and is always less than 50%.

### Variation by radioconjugates impurity

The influence of dilution of radioconjugates by unlabeled antibodies on the minimal single curative dose, *A*_*cur*_, is illustrated in **Fig. 3A**. Initial increase of the coefficient of drug impurity, *η*, affects the value of *A*_*cur*_ only slightly until the amount of injected antibodies, [*η* + 1]*A*_*cur*_, approaches the total amount of specific receptors expressed on cancer cells. The latter measure accounts for both *γN*_0_ of receptors at the moment of drug injection and the new receptors expressed during treatment.

**Figure 3:**
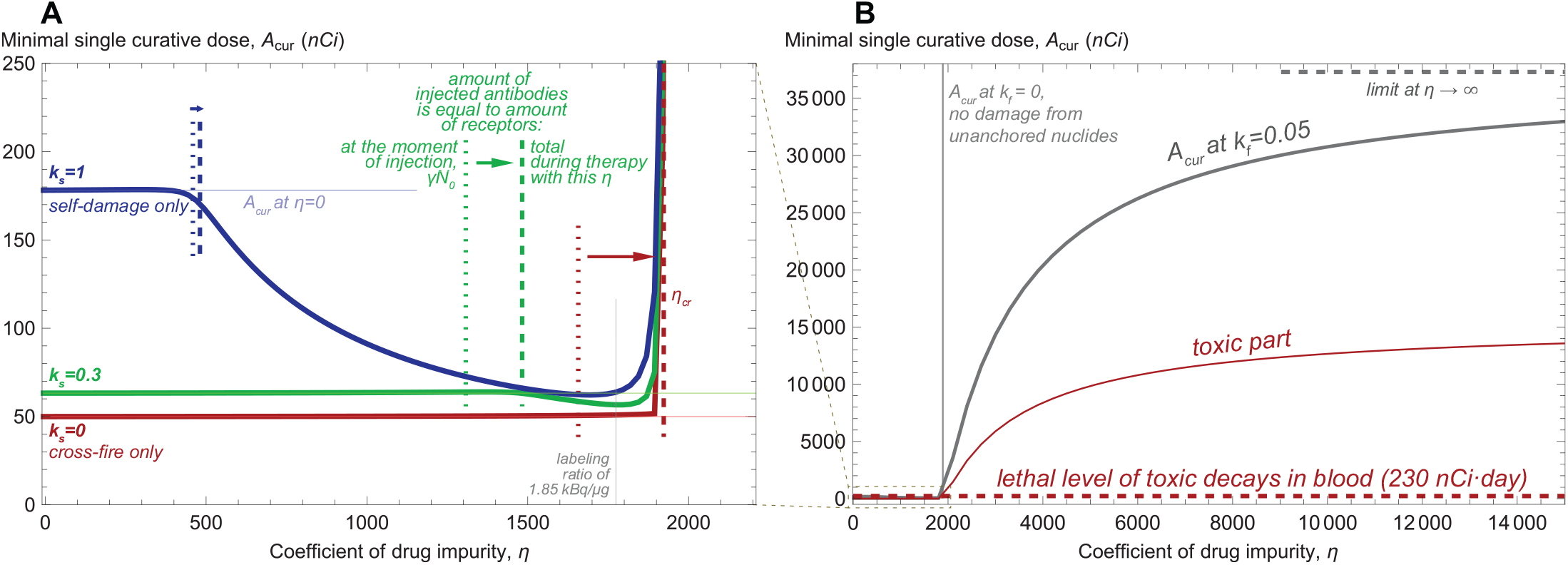
Dependence of minimal single curative dose on the coefficient of drug impurity, *η*, i.e., the ratio of unlabeled antibodies to radioconjugates. **A**, moderate variation of *η*. **B**, extended variation of *η*.

For cross-fire-only case, *A*_*cur*_ remains almost constant until achieving the corresponding threshold, after which it rapidly grows since the anchored nuclides can no longer guarantee elimination of cancer cells. Self-damage-only case shows qualitatively different behavior: as the amount of injected antibodies approaches the total amount of specific receptors, the minimal single curative dose starts decreasing. This effect arises from the combination of dynamic processes taking place when cancer cells are nearly saturated, yet a substantial amount of radioconjugates remains in the bloodstream. New free receptors are produced only by still viable cancer cells, and therefore they attract radioconjugates at a faster rate than damaged cells. In the case of significant self-damage, the radiation energy is therefore redirected towards viable cells, contributing to increase of treatment efficacy and overall to reduced curative dose.

With the decrease of self-damage significance, *k*_*s*_, this effect weakens. However, under any value of *η* the cross-fire-only case demands lower minimal single curative dose than the cases with *k*_*s*_ *>* 0 and, as supplementary Fig. S.15 shows, remains less toxic and delivers greater fraction of injected dose to viable cells. This is explained by the fact that, independently of the value of *k*_*s*_, during single-dose treatment the efficacy of radiation damage cannot decrease with time slower than *e*^*−λt*^, with the slowest decrease accompanying cross-fire-only case.

Notably, the decrease of *A*_*cur*_ under moderate drug impurity does not guarantee the accompanying decrease of treatment toxicity. When specific receptors are close to saturation, radioconjugates spend significant time in the bloodstream before binding, thus amplifying the toxic effect. At *k*_*s*_ = 0.3 the amount of toxic decays in blood grows monotonically with the increase of drug impurity, while at *k*_*s*_ = 1 it falls no more than twice compared to the pure radioconjugates case. It should be noted that the choice of optimal coefficient of drug impurity for increasing efficacy to toxicity ratio demands precise knowledge of values of model parameters. In real-life conditions, due to the inherent uncertainty and variability of characteristics, using the lowest possible coefficient of drug impurity may represent a reasonable strategy for single-dose treatment (see supplementary Fig. S.16).

**Figure 3B** corresponds to the behavior of *A*_*cur*_ upon further increase of drug impurity, where the curves for different values of *k*_*s*_ become indiscernible from each other. Under neglect of damage from unanchored nuclides, *k*_*f*_ = 0, the graph for *A*_*cur*_ skyrockets when the amount of injected antibodies exceeds the total amount of cancer receptors, implying that achieving cure becomes impossible. Under the value of *k*_*f*_, estimated by our experimental data, *A*_*cur*_ tends to a certain limit that leads to the amount of toxic decays in blood exceeding their estimated lethal level by a factor of *≈*70 (see supplementary Section S.3.1). For mathematical convenience we will refer to all the doses that allow decreasing viable cancer cell number to *N*_*cur*_ as curative, implying that they can be curative but lethally toxic simultaneously.

### Parameter sweep for labeling ratio of 1.85 kBq/μg

The above discussed results show that *cancer binding capacity*, i.e., the total number of specific receptors on cancer cells, plays a decisive role in determining whether cancer can be cured by radionuclides without exceeding lethal toxicity. This is confirmed by the global parameter sweep with coefficient of drug impurity *η* = 1780, corresponding to labeling ratio of 1.85 kBq/μg. **Fig. 4A** shows the dependence of minimal single curative dose, *A*_*cur*_, on cancer binding capacity at the moment of drug injection, *γN*_0_. All the dots in the region *A*_*cur*_ *< γN*_0_*/*[*η* + 1] correspond to low-toxic cases. In this region the amount of injected antibodies is lower than cancer binding capacity, and the analytical estimation of *A*_*cur*_, obtained under assumption of pure radioconjugates, still remains accurate (see supplementary Section S.3.2). As outlined above, during treatment with *A*_*cur*_ the number of newborn cancer cells is unlikely to exceed half of their initial number. This justifies low toxicity of cases, for which the amount of injected antibodies [*η* + 1]*A*_*cur*_ is lower than 1.5*γN*_0_ + 3 pmol. The latter coefficient compensates for slow drug binding under small amount of specific receptors.

**Figure 4:**
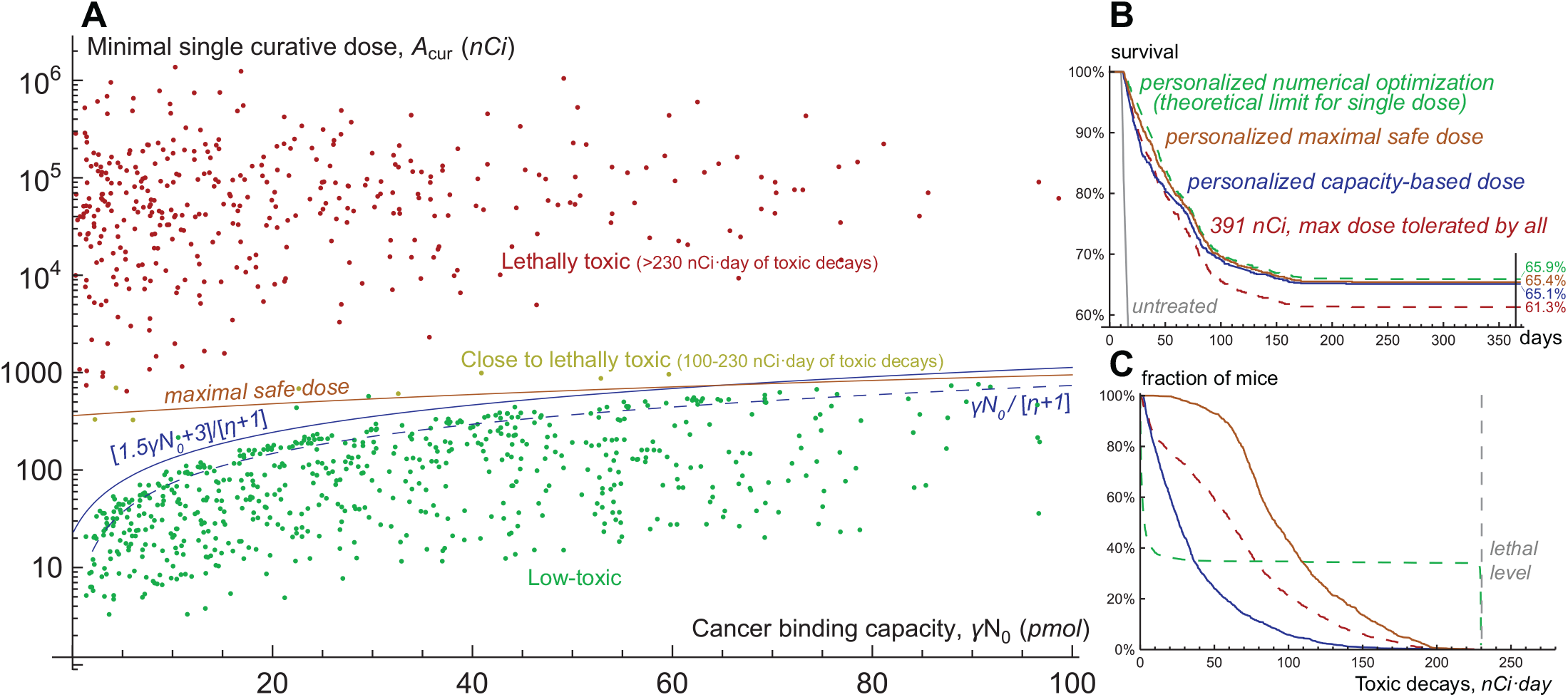
**A**, scatter plot of cancer binding capacity, i.e., total amount of specific receptors on cancer cells at the moment of drug injection, and minimal single curative dose produced by the global parameter sweep within the training set of one thousand virtual mice. Coefficient of drug impurity, *η*, corresponds to the labeling ratio of 1.85 kBq/μg. **B**, survival curves and **C**, toxicity curves for a single test set of one thousand virtual mice, treated by different approaches.

This provides a basis for personalized dosing strategy based on only one parameter, cancer binding capacity, *γN*_0_. In theory, it can be estimated before treatment. As **Fig. 2A** shows, the concentration of drug in blood after its injection follows exponential decrease, the rate of which depends on drug binding rate, *k*_*on*_; on volume of drug distribution, *V*, which by itself can be pre-estimated; and on cancer binding capacity, *γN*_0_. The latter therefore can be assessed from the pharmacokinetic curve of a preliminary small diagnostic dose, which by itself would occupy a negligible fraction of receptors.

The personalized strategy was tested in *in silico* trial within a set of another one thousand virtual mice. Its results are illustrated in **Fig. 4B**,**C**. We consider that a virtual mouse dies at the moment when the number of cancer cells reaches 10^11^, or at the moment when the total amount of decays in blood reaches 230 nCi·day. Direct numerical simulations identified 391 nCi as the maximal one-size-fits-all dose not leading to toxicity-related deaths and thus tolerated by all virtual mice. For determining the theoretical limit of single-dose treatment efficacy, we performed personalized numerical optimization, assuming precise knowledge of all model parameters for each virtual mouse, with the goal of curing a virtual mouse if it is feasible, otherwise prolonging its overall survival as much as possible. The treatment with personalized doses adjusted to cancer binding capacity showed only marginally less efficiency than such explicit numerical optimization and was accompanied by less toxicity for non-cured mice.

The acceptable toxicity levels within the considered parameter ranges are ensured by moderate amount and fast clearance of radioactive fragments of antibodies, released to the bloodstream from dead cancer cells. In a more general case injection of higher doses guided by greater cancer binding capacities can exacerbate treatment-associated toxicity. Its prediction in real-life scenario is complicated by uncertainties in relevant parameters. However, the use of reasonable physiologic ranges of corresponding parameters allows estimating *maximal safe doses*, that will definitely not result in unacceptable toxicity for any specific parameter set (see supplementary Section S.3.2). For negligible cancer binding capacity the maximal safe dose is *≈*362 nCi, being determined only by the decays of nuclides attached to intact antibodies. With the increase of cancer binding capacity maximal safe dose tends to *≈*1763 nCi, being defined by the decays of nuclides attached to fragmented antibodies. Notably, these border values are independent of the coefficient of drug impurity, but its lower values would allow using greater doses safely under lower cancer binding capacity.

As noted above, our experimental data suggests significant intrapopulation heterogeneity of cancer cells, not accounted for herein. Supplementary Section S.4 contains similar modeling study for heterogeneous cancers, which confirms the optimization potential of personalized dosing strategy based on cancer binding capacity. Further improvement of treatment efficacy can be achieved by the use of multiple doses of radioconjugates.

### Optimization of multi-dose treatment

The extension of above-discussed results that includes multi-dose treatments is presented in **Fig. 5**, where virtual mice are stratified into three groups, corresponding to low, intermediate, and high values of relative significance of self-damage, *k*_*s*_. The administration of one-size-fits-all maximum tolerated dose results in comparable one-year overall survival rates for these groups. Personalized single-dose numerical optimization assuming precise knowledge of all model parameters yields the greatest increase in treatment efficacy for the group with intermediate values of *k*_*s*_. These mice benefit from both slow decrease of cross-fire irradiation rate and redistribution of a part of radioconjugates to newborn viable cells that promotes their prolonged self-damage irradiation.

**Figure 5:**
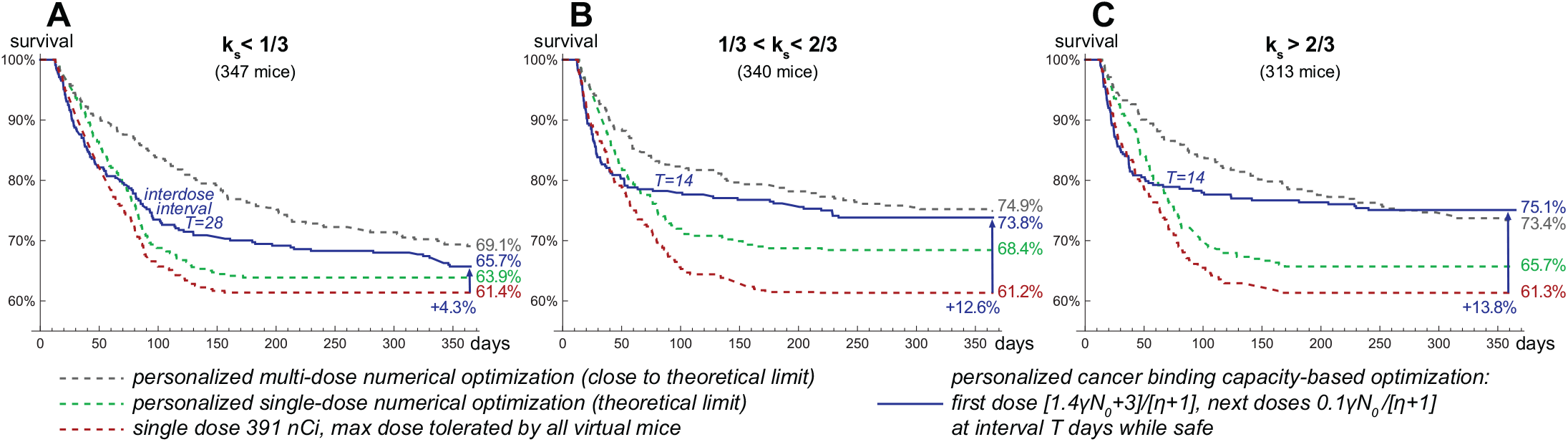
Survival curves for the test set of virtual mice, grouped by relative significance of self-damage, *k*_*s*_: **A**, low *k*_*s*_; **B**, intermediate *k*_*s*_; **C**, high *k*_*s*_. Coefficient of drug impurity, *η*, corresponds to the labeling ratio of 1.85 kBq/μg.

The algorithm of similar personalized multi-dose numerical optimization is described in supplementary Section S.5. It is based on the analysis and simulations of the auxiliary idealized version of the model wherein injected antibodies immediately bind to cancer receptors and ensure their constant saturation, with further discretization of the corresponding administered dose over variable time intervals. Analytical study of this auxiliary model shows that the necessary, but not sufficient, condition for cancer curability in the long term is the requirement of maximal possible rate of cancer cells damage due to irradiation from anchored nuclides to be greater than cell proliferation rate: *αλγ/{ν*[*η* + 1]*} > ρ*. During the prolonged treatment, however, the irradiation rate of cancer cells is impaired by inevitable accumulation of inert antibodies on their receptors. The account for this effect allows deriving more strict conditions that indicate a positive impact of self-damage irradiation efficiency on increasing the likelihood of cancer cure in the long term. Consistent with this finding, personalized multi-dose numerical optimization yields greater increase in one-year survival rate for the groups with intermediate and high values of *k*_*s*_.

The simulations of curative multi-dose treatments, applied to the virtual mice that cannot be cured by a single dose, suggest that the crucial factor enabling treatment optimization is redistribution of activity towards viable cancer cells. For some of the curative multi-dose treatments the fraction of injected activity spent on viable cancer cells exceeds the theoretical threshold for a single-dose setting, estimated above to be *<*5%. Redistribution of activity towards viable cells can be achieved via two methods, which relevance depends on the value of *k*_*s*_. Under its high values treatment efficacy benefits from maintaining near-saturation levels of cancer binding capacity, enabling redirection of further injected radioconjugates and deposited dose to newborn cancer cells. Under low values of *k*_*s*_, when cross-fire dominates, optimal treatments aim to keep a considerable fraction of cancer receptors free but initiate shrinkage of cancer mass with the first dose in order to distribute further injected radioconjugates over fewer cancer cells.

From the practical point of view, the crucial question is whether these methods can be used in a robust way given the uncertainty of parameters. For the study of this question, we followed our assumptions introduced in the previous section. The only parameter assumed to be known is cancer binding capacity at the moment of first drug injection, *γN*_0_. Based on it, the upper limit on injected doses can be set, guaranteeing that they cannot lead to lethal toxicity for any specific parameter set. Also, we assume that the range of emitted particles and approximate knowledge of cancer cell density enable stratification of mice into three groups, corresponding to low, intermediate, and high values of *k*_*s*_. Using these assumptions, within the training set of virtual mice we found optimized universal forms of schedules with doses tailored to *γN*_0_.

The administration of corresponding universal schedules in the test set of virtual mice yields an increase in treatment efficacy comparable to that in the training set, which confirms the universality of these schedules (see supplementary Section S.5.3). The greatest increase in one-year survival rate, compared to outcome of administration of maximum tolerated dose, is achieved for the group with high values of *k*_*s*_. This shows that saturation of cancer receptors enabling redistribution of further injected radioconjugates towards viable cancer cells represents a robust method of increasing treatment efficacy, valid for the use of short-range particle emitters.

The universal schedules for the group with low values of *k*_*s*_ also initially saturate cancer receptors, as their undersaturation turns out to be beneficial only for a subset of virtual mice, hampering treatment efficacy for the others. The optimal inter-dose intervals are longer for this group, as they allow getting advantage of the relatively slow process of cancer mass shrinkage to facilitate cross-fire irradiation of viable cells. This, however, allows gaining only a moderate increase in one-year survival rate, suggesting that this approach, valid for emitters of long-range particles, is less robust.

## Discussion

### Optimization of targeted radionuclide therapy in experimental setting

This study highlights the general principles that can allow increasing the efficacy of targeted radionuclide therapy (TRT) in mouse models of multiple myeloma and can be extrapolated on another cancer models. In particular, the general form of optimal multi-dose strategy is suggested for the use of high-affinity or rapidly internalizing antibodies and short-range particle emitters which provide high efficiency of self-damage irradiation of cancer cells. Namely, the first dose should aim at saturating cancer binding capacity in order to enable redistribution of significantly lower subsequent doses towards still viable cells that continue producing specific receptors. It is noteworthy, that eradication of cancer cells via self-damage irradiation may be a necessary condition for cure of disseminated multiple myeloma, since during prolonged treatment the distance between its remaining cells should gradually increase, hindering the efficiency of cross-fire [33].

The crucial factor governing the rate of cancer radiation damage and overall TRT efficacy is the density of radionuclides attached to cancer cells, which maximum value is achieved upon saturation of specific receptors and is largely determined by the labeling ratio of antibodies. The simulations performed herein considered daratumumab incubated with ^225^Ac at the labeling ratio of 1.85 kBq/μg. It is technically feasible to achieve at least 20 times higher labeling ratios in this setting, resulting in proportional increase of maximal radiation damage rate [11, 12]. However, for the first dose saturating cancer receptors the use of higher labeling ratios implies injecting greater amount of radioconjugates, which may result in unacceptable toxicity. Such risk can be prevented by dilution of radioconjugates for the first dose. The effect of the following low doses, distributed over lower number of remaining viable cells, should on contrary benefit from high labeling ratio. Such deliberate regulation of nuclide to antibody ratio should have additional advantage in case of sufficiently heterogeneous cancer cell composition, providing initial damage of relatively radiosensitive cells by low radiation energy density, with its further increase for eradication of remaining radioresistant cells.

Another factor that determines the rate of cancer radiation damage is the rate of nuclide decay. From this perspective, the use of radium isotope ^224^Ra, which decay chain yields *≈*2.5 faster release of *α*-particle energy, comparable to that of ^225^Ac [31], represents a potentially more effective option for radiopharmaceutical therapy of blood cancers, that should mitigate the negative impact of cancer cell proliferation on treatment efficacy.

### Application in clinical setting

Multiple myeloma is currently considered as a not curable disease [43], spurring the debates on curative versus control doctrines towards its treatment [44]. High and uniform expression of CD38 in its cells, however, provides great potential for its effective treatment with targeted radioconjugates, which can be optimized based on the principles derived herein. Labeling of daratumumab with ^225^Ac leads to significant increase of its anticancer potency, suggesting that radiation damage should be regarded as the major factor for the effect of ^225^Ac-DOTA-daratumumab [11, 12]. Clinical trials with daratumumab did not identify its maximum tolerated dose, which suggests feasibility of using this antibody for saturation of CD38 [4]. However, injections of the amounts of antibodies significantly exceeding cancer binding capacity should be avoided as impeding the binding of further injected radioconjugates to cancer cells. Importantly, prolonged saturation of CD38 was witnessed in clinical setting several weeks after termination of daratumumab treatment, underscoring the necessity of tailoring the first dose to cancer binding capacity for efficient prolonged TRT [45].

In this work we suggest that estimation of cancer binding capacity can be performed via preliminary injection of a small dose of diagnostic radioconjugates and measurement of their pharmacokinetic curve. This method implies that the pool of CD38 receptors on cancer cells acts as the major attractor of radioconjugates. However, as we have shown previously, daratumumab initially preferentially binds to FcRn receptors, that ensure its recycling and maintaining in circulation [13]. Preliminary cold dosing by unlabeled daratumumab for saturation of FcRn receptors allows redistributing further injected radioconjugates towards cancer cells, however, it is as well associated with the risk of prolonged saturation of CD38 by antibodies devoid of radionuclides. One option to overcome this problem is to saturate FcRn by antibodies that do not bind to CD38. Several FcRn-blocking antibodies are currently in clinical development, with efgartigimod alfa having obtained FDA approval as the first-in-class medication of this type [46]. Another option is to use engineered antibodies with mutated Fc-FcRn binding sites for radiolabeling, as they would display significantly faster targeting of cancer cells [47].

Another complication accompanying clinical setting is on-target off-site toxicity of radioconjugates binding to CD38 expressed in healthy cells. However, low levels of CD38 expression can protect them from excessive irradiation [32]. Differential expression of receptors in healthy and cancer cells may guide the optimal ratio of radioconjugates and antibodies during TRT, while the rate of healthy cells repopulation may be utilized to adjust the timing of injections, allowing for toxicity recovery.

The mentioned complications can be addressed via quantitative systems pharmacology modeling incorporating clinical data [48]. With that approach, the solutions of treatment optimization tasks, based on collection of patient data and leveraging on the mechanistic insights gained in this study, can facilitate treatment optimization in clinical setting.

The principles derived in this study can be adapted to treatment of extramedullary lesions of multiple myeloma as well as to solid cancers. The penetration of drug to their sites is impeded by lower permeability of associated capillaries, compared to bone marrow site, resulting in only a moderate fraction of injected activity reaching cancer cells [49]. The denser arrangement of cells in solid tumors, however, should promote their effective cross-fire irradiation, as implied by complete remission of solid tumors reported for experimental setting using *β*-particle emitters [50]. In light of this, combination of initial administration of *β*-particle emitters or external beam irradiation to solid tumors with subsequent injections of *α*-particle emitters for redirection of deposited dose towards still viable cells represents an intriguing concept for further investigation.

## Supporting information

Computational codes

## Authors’ Contributions

**Conception and design:** M. Kuznetsov, V. Adhikarla, R.C. Rockne

**Development of methodology:** M. Kuznetsov, V. Adhikarla, R.C. Rockne

**Acquisition of data (provided animals, acquired and managed patients, provided facilities, etc**.): E. Caserta

**Analysis and interpretation of data (e**.**g**., **statistical analysis, biostatistics, computational analysis):** M. Kuznetsov, V. Adhikarla, R.C. Rockne

**Writing, review, and/or revision of the manuscript:** M. Kuznetsov, V. Adhikarla, R.C. Rockne

**Administrative, technical, or material support (i**.**e**., **reporting or organizing data, constructing databases):** J.E. Shively, X. Wang, F. Pichiorri, R.C. Rockne

**Study supervision:** J.E. Shively, X. Wang, F. Pichiorri, R.C. Rockne

## Acknowledgments

Research reported in this publication was supported by the National Cancer Institute of the National Institutes of Health under grant numbers P30CA033572, R01CA238429. The content is solely the responsibility of the authors and does not necessarily represent the official views of the National Institutes of Health. The authors thank Dr. S. Peter Wu for fruitful discussions.

## References

[1] Sacha Zinn, Rodrigo Vazquez-Lombardi, Carsten Zimmermann, Puja Sapra, Lutz Jermutus, and Daniel Christ. Advances in antibody-based therapy in oncology. Nature Cancer, 4(2):165–180, 2023.

[2] Haidong Wang, Mohsen Naghavi, Christine Allen, Ryan M Barber, Zulfiqar A Bhutta, Austin Carter, et al. Global, regional, and national life expectancy, all-cause mortality, and cause-specific mortality for 249 causes of death, 1980– 2015: a systematic analysis for the Global Burden of Disease Study 2015. The lancet, 388(10053):1459–1544, 2016.

[3] Colin Phipps, Yunxin Chen, Sathish Gopalakrishnan, and Daryl Tan. Daratumumab and its potential in the treatment of multiple myeloma: overview of the preclinical and clinical development. Therapeutic advances in hematology, 6(3):120–127, 2015.

[4] Danai Dima, Joshua Dower, Raymond L Comenzo, and Cindy Varga. Evaluating daratumumab in the treatment of multiple myeloma: safety, efficacy and place in therapy. Cancer management and research, pages 7891–7903, 2020.

[5] Ilaria Saltarella, Vanessa Desantis, Assunta Melaccio, Antonio Giovanni Solimando, Aurelia Lamanuzzi, Roberto Ria, et al. Mechanisms of resistance to anti-CD38 daratumumab in multiple myeloma. Cells, 9(1):167, 2020.

[6] Domenico Viola, Ada Dona, Enrico Caserta, Estelle Troadec, Francesca Besi, Tinisha McDonald, et al. Daratumumab induces mechanisms of immune activation through CD38+ NK cell targeting. Leukemia, 35(1):189–200, 2021.

[7] Kilian E Salerno, Soumyajit Roy, Cathy Ribaudo, Teresa Fisher, Ravi B Patel, Esther Mena, and Freddy E Escorcia. A primer on radiopharmaceutical therapy. International Journal of Radiation Oncology* Biology* Physics, 2022.

[8] Richard L Wahl, George Sgouros, Amir Iravani, Heather Jacene, Daniel Pryma, Babak Saboury, et al. Normal-tissue tolerance to radiopharmaceutical therapies, the knowns and the unknowns. Journal of Nuclear Medicine, 62(Supplement 3):23S–35S, 2021.

[9] Cecilia Hindorf, Gerhard Glatting, Carlo Chiesa, Ola Lindén, and Glenn Flux. EANM Dosimetry Committee guidelines for bone marrow and whole-body dosimetry. European journal of nuclear medicine and molecular imaging, 37:1238–1250, 2010.

[10] Megan Minnix, Vikram Adhikarla, Enrico Caserta, Erasmus Poku, Russell Rockne, John E Shively, and Flavia Pichiorri. Comparison of CD38-targeted α-versus β-radionuclide therapy of disseminated multiple myeloma in an animal model. Journal of Nuclear Medicine, 62(6):795–801, 2021.

[11] Mark S Berger, Keisha Thomas, Dadachova Ekaterina, Mackenzie Malo, Rubin Jiao, Kevin Allen, and Colin Clarke. Actinium labeled daratumumab demonstrates enhanced killing of multiple myeloma cells over naked daratumumab. Blood, 130:4427, 2017.

[12] Wojciech Dawicki, Kevin JH Allen, Rubin Jiao, Mackenzie E Malo, Muath Helal, Mark S Berger, et al. Daratumumab-225Actinium conjugate demonstrates greatly enhanced antitumor activity against experimental multiple myeloma tumors. Oncoimmunology, 8(8):1607673, 2019.

[13] Amrita Krishnan, Vikram Adhikarla, Erasmus K Poku, Joycelynne Palmer, Ammar Chaudhry, Van Eric Biglang-awa, et al. Identifying CD38+ cells in patients with multiple myeloma: First-in-human imaging using copper-64-labeled daratumumab. Blood Advances, 4(20):5194–5202, 2020.

[14] Maxim Kuznetsov, Jean Clairambault, and Vitaly Volpert. Improving cancer treatments via dynamical biophysical models. Physics of life reviews, 39:1–48, 2021.

[15] Simon Arsène, Yves Parès, Eliott Tixier, Solène Granjeon-Noriot, Bastien Martin, Lara Bruezière, et al. In silico clinical trials: Is it possible? In High Performance Computing for Drug Discovery and Biomedicine, pages 51–99. Springer, 2023.

[16] Thomas O McDonald, Yu-Chen Cheng, Christopher Graser, Phillip B Nicol, Daniel Temko, and Franziska Michor. Computational approaches to modelling and optimizing cancer treatment. Nature Reviews Bioengineering, 1(10):695–711, 2023.

[17] Huixi Zou, Parikshit Banerjee, Sharon Shui Yee Leung, and Xiaoyu Yan. Application of pharmacokinetic-pharmacodynamic modeling in drug delivery: development and challenges. Frontiers in pharmacology, 11:997, 2020.

[18] David J Brenner, LR Hlatky, PJ Hahnfeldt, Y Huang, and RK Sachs. The linear-quadratic model and most other common radiobiological models result in similar predictions of time-dose relationships. Radiation research, 150(1):83–91, 1998.

[19] Peter Kletting, Christiane Schuchardt, Harshad R Kulkarni, Mostafa Shahinfar, Aviral Singh, Gerhard Glatting, et al. Investigating the effect of ligand amount and injected therapeutic activity: a simulation study for 177Lu-labeled PSMA-targeting peptides. PLoS One, 11(9):e0162303, 2016.

[20] Peter Kletting, Thomas Kull, Christian Maaß, Noeen Malik, Markus Luster, Ambros J Beer, and Gerhard Glatting. Optimized peptide amount and activity for 90Y-labeled DOTATATE therapy. Journal of Nuclear Medicine, 57(4):503–508, 2016.

[21] Peter Kletting, Anne Thieme, Nina Eberhardt, Andreas Rinscheid, Calogero D’Alessandria, Jakob Allmann, et al. Modeling and predicting tumor response in radioligand therapy. Journal of nuclear medicine, 60(1):65–70, 2019.

[22] Nouran RR Zaid, Peter Kletting, Gordon Winter, Vikas Prasad, Ambros J Beer, and Gerhard Glatting. A physiologically based pharmacokinetic model for in vivo alpha particle generators targeting neuroendocrine tumors in mice. Pharmaceutics, 13(12):2132, 2021.

[23] Yoshiharu Yonekura, Hiroshi Toki, Tadashi Watabe, Kazuko Kaneda-Nakashima, Yoshifumi Shirakami, Ooe, et al. Mathematical model for evaluation of tumor response in targeted radionuclide therapy with 211At using implanted mouse tumor. International Journal of Molecular Sciences, 23(24):15966, 2022.

[24] Alireza Karimian, Nathan T Ji, Hong Song, and George Sgouros. Mathematical modeling of preclinical alpha-emitter radiopharmaceutical therapy. Cancer research, 80(4):868–876, 2020.

[25] Vikram Adhikarla, Dennis Awuah, Alexander B Brummer, Enrico Caserta, Amrita Krishnan, Flavia Pichiorri, et al. A mathematical modeling approach for targeted radionuclide and chimeric antigen receptor T cell combination therapy. Cancers, 13(20):5171, 2021.

[26] Vikram Adhikarla, Dennis Awuah, Enrico Caserta, Megan Minnix, Maxim Kuznetsov, Amrita Krishnan, et al. Designing combination therapies for cancer treatment: Application of a mathematical framework combining CAR T-cell immunotherapy and targeted radionuclide therapy. Frontiers in Immunology, 15:1358478, 2023.

[27] David A Scheinberg and Michael R McDevitt. Actinium-225 in targeted alpha-particle therapeutic applications. Current radiopharmaceuticals, 4(4):306–320, 2011.

[28] Natascia Di Iorgi, Ashley O Mo, Kate Grimm, Tishya AL Wren, Frederick Dorey, and Vicente Gilsanz. Bone acquisition in healthy young females is reciprocally related to marrow adiposity. The Journal of Clinical Endocrinology & Metabolism, 95(6):2977–2982, 2010.

[29] Michael C Joiner and Albert J van der Kogel. Basic clinical radiobiology. CRC press, 2018.

[30] Maxim Kuznetsov and Andrey Kolobov. Optimization of antitumor radiotherapy fractionation via mathematical modeling with account of 4 R’s of radiobiology. Journal of Theoretical Biology, 558:111371, 2023.

[31] Sophie Poty, Lynn C Francesconi, Michael R McDevitt, Michael J Morris, and Jason S Lewis. α-Emitters for radiotherapy: from basic radiochemistry to clinical studies – Part 1. Journal of Nuclear Medicine, 59(6):878–884, 2018.

[32] Jayeeta Ghose, Domenico Viola, Cesar Terrazas, Enrico Caserta, Estelle Troadec, Jihane Khalife, et al. Daratumumab induces CD38 internalization and impairs myeloma cell adhesion. Oncoimmunology, 7(10):e1486948, 2018.

[33] HE Daldrup-Link, TM Link, EJ Rummeny, C August, S Könemann, H Jürgens, and W Heindel. Assessing permeability alterations of the blood–bone marrow barrier due to total body irradiation: in vivo quantification with contrast enhanced magnetic resonance imaging. Bone marrow transplantation, 25(1):71–78, 2000.

[34] Amin I Kassis. Therapeutic radionuclides: biophysical and radiobiologic principles. In Seminars in nuclear medicine, volume 38, pages 358–366. Elsevier, 2008.

[35] Gert Luurtsema, Verena Pichler, Salvatore Bongarzone, Yann Seimbille, Philip Elsinga, Antony Gee, and Johnny Vercouillie. EANM guideline for harmonisation on molar activity or specific activity of radiopharmaceuticals: Impact on safety and imaging quality. EJNMMI Radiopharmacy and Chemistry, 6:1–16, 2021.

[36] Andrew Brown and Herman Suit. The centenary of the discovery of the Bragg peak. Radiotherapy and Oncology, 73(3):265–268, 2004.

[37] Btissam Chami, Makoto Okuda, Morvarid Moayeri, France Pirenne, Yoko Hidaka Nambiar, et al. Anti-CD38 monoclonal antibody interference with blood compatibility testing: Differentiating isatuximab and daratumumab via functional epitope mapping. Transfusion, 62(11):2334–2348, 2022.

[38] Ryan L Kelly, Tingwan Sun, Tushar Jain, Isabelle Caffry, Yao Yu, Yuan Cao, et al. High throughput cross-interaction measures for human IgG1 antibodies correlate with clearance rates in mice. In MAbs, volume 7, pages 770–777. Taylor & Francis, 2015.

[39] Ian D Ferguson, Bonell Patiño-Escobar, Sami T Tuomivaara, Yu-Hsiu T Lin, Matthew A Nix, Kevin K Leung, et al. The surfaceome of multiple myeloma cells suggests potential immunotherapeutic strategies and protein markers of drug resistance. Nature communications, 13(1):4121, 2022.

[40] Jaroslaw Wajs and Waldemar Sawicki. The morphology of myeloma cells changes with the progression of the disease. Contemporary Oncology/Wspólczesna Onkologia, 17(3):272–275, 2013.

[41] Estefanía García-Guerrero, Ralph Götz, Sören Doose, Markus Sauer, Alfonso Rodríguez-Gil, Thomas Nerreter, et al. Upregulation of CD38 expression on multiple myeloma cells by novel HDAC6 inhibitors is a class effect and augments the efficacy of daratumumab. Leukemia, 35(1):201–214, 2021.

[42] Xili Liu, Maria Moscvin, Seungeun Oh, Tianzeng Chen, Wonshik Choi, Benjamin Evans, et al. Characterizing dry mass and volume changes in human multiple myeloma cells upon treatment with proteotoxic and genotoxic drugs. Clinical and Experimental Medicine, 23(7):3821–3832, 2023.

[43] Mohamad Mohty, Hervé Avet-Loiseau, Florent Malard, and Jean-Luc Harousseau. Potential future direction of measurable residual disease evaluation in multiple myeloma. Blood, 142(18):1509–1517, 2023.

[44] Mark Roschewski, Neha Korde, S Peter Wu, and Ola Landgren. Pursuing the curative blueprint for early myeloma. Blood, The Journal of the American Society of Hematology, 122(4):486–490, 2013.

[45] Anna Oberle, Anna Brandt, Malik Alawi, Claudia Langebrake, Snjezana Janjetovic, Christine Wolschke, et al. Long-term CD38 saturation by daratumumab interferes with diagnostic myeloma cell detection. Haematologica, 102(9):e368, 2017.

[46] Marco Meglio. FDA approves subcutaneous efgartigimod as treatment for generalized myasthenia gravis. Neurology Live, 2023.

[47] Vania Kenanova, Tove Olafsen, Desiree M Crow, Gobalakrishnan Sundaresan, Murugesan Subbarayan, Nora H Carter, et al. Tailoring the pharmacokinetics and positron emission tomography imaging properties of anti– carcinoembryonic antigen single-chain Fv-Fc antibody fragments. Cancer research, 65(2):622–631, 2005.

[48] Bruna Scheuher, Khem Raj Ghusinga, Kimiko McGirr, Maksymilian Nowak, Sheetal Panday, Joshua Apgar, et al. Towards a platform quantitative systems pharmacology (QSP) model for preclinical to clinical translation of antibody drug conjugates (ADCs). Journal of Pharmacokinetics and Pharmacodynamics, pages 1–19, 2023.

[49] Megan Minnix, Lin Li, Paul J Yazaki, Aaron D Miller, Junie Chea, Erasmus Poku, et al. TAG-72–targeted αradionuclide therapy of ovarian cancer using 225Ac-labeled DOTAylated-huCC49 antibody. Journal of Nuclear Medicine, 62(1):55–61, 2021.

[50] Sara Lundsten, Diana Spiegelberg, Nakul R Raval, and Marika Nestor. The radiosensitizer onalespib increases complete remission in 177Lu-DOTATATE-treated mice bearing neuroendocrine tumor xenografts. European journal of nuclear medicine and molecular imaging, 47:980–990, 2020.

