## Supplemental material for "Mathematical Modeling Unveils Optimization Strategies for Targeted Radionuclide Therapy of Blood Cancers"

### S.1 Materials and Methods

#### S.1.1 Animal studies

Briefly, the previously reported animal studies [1] were performed as follows.

Daratumumab (Janssen Biotech Inc.) or control trastuzumab antibodies were reacted with a 30 M excess of the chelator DOTA-mono-N-hydroxysuccinimide ester (Macrocyclics, Inc.). DOTA-conjugated antibody (50 μg) was incubated with $^{225}$Ac (Oak Ridge National Laboratory) at a labeling ratio of 1.85 kBq/μg for 45 min at 43°C and chased with 1 mM diethylenetriamine pentaacetic acid.

Animal studies were conducted on 6- to 10-week-old NOD.Cg-Prkdc$^{scid}$Il2rg$^{tm1Wjl}$/SzJ mice (NSG; Jackson Laboratory). Green fluorescent protein luciferase-positive MM.1S cells (Dana-Farber Cancer Institute) were injected intravenously, at $5{\cdot}10^6$ cells/200 μL of phosphate-buffered saline per mouse. Tumor distribution and growth were followed by serial whole-body imaging on the Lago X (Spectral Instruments Imaging). Before imaging, the animals were anesthetized with 4% isoflurane and injected intraperitoneally with 200 μL of D-luciferin (15 mg/mL) in sterile phosphate-buffered saline. After nine days, mice were randomized and treated with saline; 22.2 kBq of $^{225}$Ac-DOTA-trastuzumab; and 0.925 kBq, 1.85 kBq, 3.7 kBq, 11.1 kBq or 22.2 kBq of $^{225}$Ac-DOTA-daratumumab. Mice were given intravenous immunoglobulin by intraperitoneal injection 2 hours before the injection of radioconjugates. All therapy doses were made up to 30 μg of antibody.

The experimental data is reproduced in Fig. S.1. We assume that total radiance signal over the mouse body is proportional to the current number of alive, both viable and damaged, cancer cells in it.

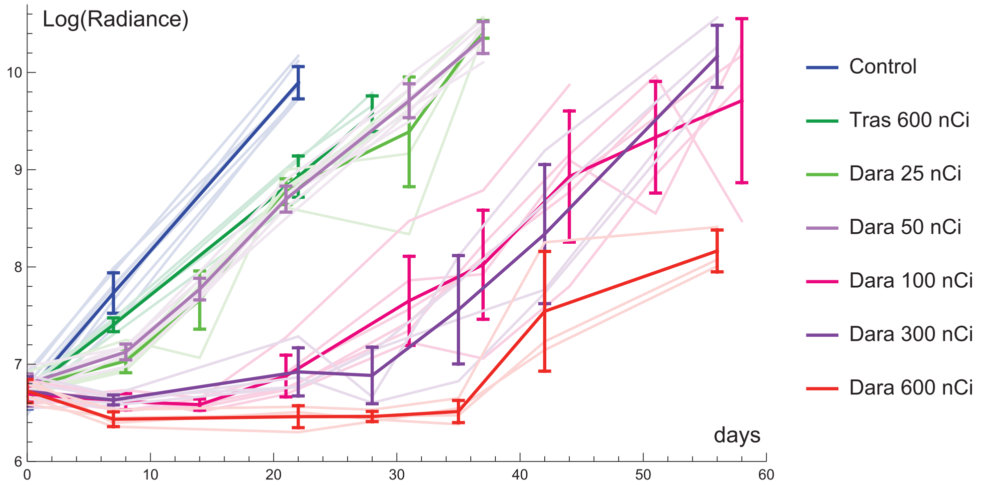


Figure S.1: Dose response for treatment of disseminated multiple myeloma mouse model, reproduced by data of the work [1]. Injected radioactivity values are indicated in the legend. Lighter lines correspond to individual mice. Tras=$^{225}$Ac-DOTA-trastuzumab, Dara=$^{225}$Ac-DOTA-daratumumab.

#### S.1.2 Derivation of equations for the receptors of cancer cells

For convenience, let's specify what quantities and measures are followed in the model. The numbers of viable and damaged cancer cells are denoted as $N$ and $D$. Since they can achieve high orders of magnitude, transient normalization parameters are introduced for them in this study for numerical convenience.

Concentrations of active and inert antibodies in blood, measured in $nM = pmol/ml$, are designated as $a$ and $b$. Total volume of drug distribution, further referred to as plasma volume for simplicity, is $V$, measured in $ml$. $A = aV$ and $B = bV$ are the total amounts of active and inert antibodies in the system, measured in $pmol$. Each normalized number of cancer cells has $\gamma$ pmols of specific receptors. The total number of receptors in the system is $\gamma[N + D]$. The concentration of receptors in plasma is $\frac{\gamma}{V}[N + D]$.

Fractions of certain types of receptors of cancer cells, which are numbers in the range $[0, 1]$, are denoted as $f_{FN}$, $f_{AN}$, $f_{BN}$ for the fractions of free (unbound), active and inert receptors of viable cancer cells, and $f_{FD}$, $f_{AD}$, $f_{BD}$ for the fractions or free, active and inert receptors of damaged cancer cells. Note that $f_{FN} + f_{AN} + f_{BN} = 1$ and $f_{FD} + f_{AD} + f_{BD} = 1$.

Since concentrations of antibodies in blood plasma and cancer cells microenvironment are assumed to be equal, we can introduce the measures of concentration of free receptors of viable and damaged cells in blood: $n_F = f_{FN}\frac{\gamma}{V}N$ and $d_F = f_{FD}\frac{\gamma}{V}D$. Total concentration of free receptors is $r_F = \frac{\gamma}{V}[f_{FN}N + f_{FD}D]$.

Following the model assumptions, the equations for the concentrations of receptors of viable cancer cells are as follows, where $R_D \equiv R_D(N, D, f_{AN}, f_{AD}, a, p)$ for brevity.

$$\begin{aligned}
\text{Free receptors of viable cells:}\quad n'_F &= \overbrace{\rho[n_F + n_A + n_B]}^{\text{proliferation}} \overbrace{-k_{on}[a + b]n_F}^{\text{binding}} \overbrace{-R_D n_F}^{\text{irradiation}};\\
\text{Active receptors of viable cells:}\quad n'_A &= \overbrace{k_{on} a n_F}^{\text{binding}} \overbrace{-\lambda n_A}^{\text{decay}} \overbrace{-R_D n_A}^{\text{irradiation}};\\
\text{Inert receptors of viable cells:}\quad n'_B &= \overbrace{k_{on} b n_F}^{\text{binding}} \overbrace{+\lambda n_A}^{\text{decay}} \overbrace{-R_D n_B}^{\text{irradiation}}.
\end{aligned}$$

Using $n_F = \frac{\gamma}{V} f_{FN} N$ as well as similar substitutions for $n_A$ and $n_B$, and reducing $\frac{\gamma}{V}$, we get:

$$\begin{aligned}
f_{FN}N' + f'_{FN}N &= \overbrace{\rho N}^{\text{proliferation}} \overbrace{-k_{on}[a + b]f_{FN}N}^{\text{binding}} \overbrace{-R_D f_{FN}N}^{\text{irradiation}};\\
f_{AN}N' + f'_{AN}N &= \overbrace{k_{on} a f_{FN}N}^{\text{binding}} \overbrace{-\lambda f_{AN}N}^{\text{decay}} \overbrace{-R_D f_{AN}N}^{\text{irradiation}};\\
f_{BN}N' + f'_{BN}N &= \overbrace{k_{on} b f_{FN}N}^{\text{binding}} \overbrace{+\lambda f_{AN}N}^{\text{decay}} \overbrace{-R_D f_{BN}N}^{\text{irradiation}}.
\end{aligned}$$

And using the equation for the number of viable cells ($N' = \rho N - R_D N$) and reducing $N$, we obtain:

$$\begin{aligned}
f'_{FN} &= [1 - f_{FN}]\rho - k_{on}[a + b]f_{FN};\\
f'_{AN} &= k_{on} a f_{FN} - \lambda f_{AN} - \rho f_{AN};\\
f'_{BN} &= k_{on} b f_{FN} + \lambda f_{AN} - \rho f_{BN}.
\end{aligned}$$

There is no need to consider equation for $f_{BN}$ explicitly since it is always equal to $1 - f_{FN} - f_{AN}$. The other two equations are present in the model.

Similarly, the equations for the concentrations of receptors of damaged cancer cells are:

$$\begin{aligned}
\text{Free receptors of damaged cells:}\quad d'_F &= \overbrace{R_D n_F}^{\text{irradiation}} \overbrace{-k_{on}[a + b]d_F}^{\text{binding}} \overbrace{-\omega d_F}^{\text{degradation}};\\
\text{Active receptors of damaged cells:}\quad d'_A &= \overbrace{R_D n_A}^{\text{irradiation}} \overbrace{+k_{on} a d_F}^{\text{binding}} \overbrace{-\lambda d_A}^{\text{decay}} \overbrace{-\omega d_A}^{\text{degradation}};\\
\text{Inert receptors of damaged cells:}\quad d'_B &= \overbrace{R_D n_B}^{\text{irradiation}} \overbrace{+k_{on} b d_F}^{\text{binding}} \overbrace{+\lambda d_A}^{\text{decay}} \overbrace{-\omega d_B}^{\text{degradation}}.
\end{aligned}$$

Using $d_F = \frac{\gamma}{V} f_{FD} D$ as well as similar substitutions for other receptors and reducing $\frac{\gamma}{V}$, we get:

$$\begin{aligned}
f_{FD}D' + f'_{FD}D &= R_D f_{FN}N - k_{on}[a + b]f_{FD}D - \omega f_{FD}D;\\
f_{AD}D' + f'_{AD}D &= R_D f_{AN}N + k_{on} a f_{FD}D - \lambda f_{AD}D - \omega f_{AD}D;\\
f_{BD}D' + f'_{BD}D &= R_D f_{BN}N + k_{on} b f_{FD}D + \lambda f_{AD}D - \omega f_{BD}D.
\end{aligned}$$

Using the equation for the number of damaged cells ($D' = R_D N - \omega D$) it can be presented as:

$$f'_{FD} = [f_{FN} - f_{FD}]R_D \frac{N}{D} - k_{on}[a+b]f_{FD};$$

$$f'_{AD} = [f_{AN} - f_{AD}]R_D \frac{N}{D} + k_{on} a f_{FD} - \lambda f_{AD};$$

$$f'_{BD} = [f_{BN} - f_{BD}]R_D \frac{N}{D} + k_{on} b f_{FD} + \lambda f_{AD}.$$

The first two equations are present in the main model.

##### S.1.3 Fitting of experimental data and estimation of corresponding parameters

The computational codes for this section are provided in supplementary file "01-Fitting-data.nb". All codes for this study were designed in Wolfram Mathematica, version 13.3.1.0.

###### S.1.3.1 Model parameters

The clearance rate of daratumumab was assessed based on the data of clearance of human IgG1 antibodies, to the type of which it belongs, in mice [2]. The clearance rate of fragments of antibodies is assumed to be higher than that of intact antibodies.

The study [3] explicitly quantified the number of CD38 receptors on cells of three experimental myeloma cell lines and five patients by glycoprotein cell surface capture approach. That yielded the range of 16,251–613,422 receptors per cell. The value of ≈126,000 receptors per cell was obtained for MM.1S cell line that was used in our experiments. Therefore, this value is used as the basic one. Another estimation of the number of CD38 receptors on a MM.1S cell can be obtained by multiplying their average surface density, which was estimated as 40 per μm$^2$ in the work [4], and the surface area of a myeloma cell, which was estimated as ≈200 μm$^2$ for newly diagnosed patients in the work [5]. Such implicit approach yields notably lower number of ≈8000 receptors per cell. The variation range, used herein, spans from the minimal to the maximal of the obtained values.

Total volume of drug distribution $V$ is estimated to be close to mouse plasma volume, given the typical mouse blood volume of 1.5-2.5 ml and hematocrit of ≈0.5. The volume occupied by cancer cells is roughly estimated on the basis of experimentally measured volume of a multiple myeloma cell in the work [6], which is ≈1.3 pl, under the assumption that cancer cells occupy about half of the lesion volume. Note that from the mathematical point of view this parameter can be reduced by redefining $\alpha = \alpha/\nu$ and $k_f = \nu k_f$. However, it is maintained for the sake of clarity of physical meaning.

The radiosensitivity parameter $\alpha$ is formally measured in nM$^{-1}$, which implies sensitivity of cancer cells to a certain amount of decays within a unit volume. Calculation of the coefficient of conversion of radiosensitivity parameter into traditional reversed Grays can be performed as follows.

The cumulative energy $E$ released from all $\alpha$-particles emitted from a single disintegration of $^{225}$Ac and their daughter nuclei is 27.65 MeV or $E = 4.42 \cdot 10^{-12}$ J [7]. We assume red marrow density to be constant and estimate it as $P \approx 1.06$ g/ml [8].

Let's consider 1 pmol, or $10^{-12} \cdot N_A$ (where $N_A$ is Avogadro constant), of $^{225}$Ac decays happening within 1 ml of red marrow. The concentration of decays under consideration is thus 1 nM. The concentration of released energy, expressed in $Gy = J/kg$ is

$$\epsilon = \frac{10^{-12} \cdot N_A \cdot E}{10^{-3} \cdot P},$$

which is ≈2500 Gy. Therefore, in terms of radiosensitivity, 1 nM$^{-1}$ corresponds to ≈0.0004 Gy$^{-1}$.

The initial number of cancer cells $N(0) = N_0$ in this work corresponds to the moment of time, when the first drug injection is performed. In the experimental setting, the mice were injected $5 \cdot 10^6$ multiple myeloma cells intravenously 9 days before the administration of radioconjugates. The upper boundary for the initial number of cancer cells was obtained under the assumption that during this period the cells cannot proliferate faster than they do in control mice group after engraftment.

The radioactivity dose of injected $^{225}$Ac in the experiments was in the range $25 - 600$ nCi. The corresponding numbers of injected radioactive molecules were calculated straightforwardly via dividing the radioactivity by $N_A\lambda$. During the model simulations we did not set the limits for injected doses.

A virtual mouse is considered cured if the number of viable cancer cells falls down to $N_{cur} = 0.01$. Since we deal with continuous modeling, the number of cells is expressed in real numbers rather than in integers. The classical approach towards interpreting low numbers of cancer cells during such type of modeling is considering cancer cure probability as $CP(N_m) = e^{-N_m}$ [9]. This formula follows from the Poisson dose–response model, and it roughly means that if an infinite number of cancers on average have $N_{cur}$ viable cells, provided that cells are distributed randomly, then $CP(N_m)$ of them contain no viable cells. Thus, we take the level of cure probability $CP(N_{cur}) \approx 99\%$ as sufficient for our study.

Following our previous modeling [7], we consider that a virtual mouse dies at the moment when the number of cancer cells reaches $C_d = 10^{11}$. Also a mouse can die of toxicity at the moment when the total amount of decays in blood reaches $A_{bl}^{cr} = 230$ nCi·day. This level is estimated from our experimental data, see Section S.4.1.

###### S.1.3.2 Coefficient of drug impurity

For the experiments under consideration, radioconjugates were prepared by incubation of antibody (daratumumab or trastuzumab) with $^{225}$Ac at the labeling ratio of 1.85 kBq/μg. The molecular mass of daratumumab is ≈148 kg/mol, which is within a common range of weights of monoclonal antibodies. Therefore, 1 μg of daratumumab corresponds to ≈6.76 pmol. At that, 1.85 kBq of $^{225}$Ac corresponds to 0.0038 pmol of it. Thus, this process of drug preparation yielded ≈1780 unlabeled antibody molecules per every radioconjugate.

Further, all doses were made up to 30 μg of antibody, which corresponds to ≈202.7 pmol of daratumumab. The resulting coefficients of drug impurity $\eta$ for experimental settings with daratumumab are listed in Table S.1.

Table S.1: Coefficient of drug impurity $\eta$ for groups treated with $^{225}$Ac-DOTA-daratumumab.

| **dose, nCi** | **dose, kBq** | **Ac, pmol** | **Dara, pmol** | $\eta$ |
|---|---|---|---|---|
| 25 | 0.925 | 0.0019 | 202.7 | 106,683 |
| 50 | 1.85 | 0.0038 | 202.7 | 53,341 |
| 100 | 3.7 | 0.0076 | 202.7 | 26,670 |
| 300 | 11.1 | 0.0228 | 202.7 | 8889 |
| 600 | 22.2 | 0.0456 | 202.7 | 4444 |

###### S.1.3.3 Cancer cells proliferation rate

For estimation of cancer cell proliferation rate, we use the data of the control group. Since the growth of cancer cells is assumed to be exponential, the logarithm of radiance $LR$ should grow linearly, $LR = LR_0 \cdot \rho t$, where $LR_0 \approx 6.7$ is the average logarithm of radiance on day 0. By varying $\rho$ and minimizing the sum of squared deviations of logarithms of radiance for experimental and simulated data, we obtained $\rho \approx 0.34$ as the optimal value, see Fig. S.2.

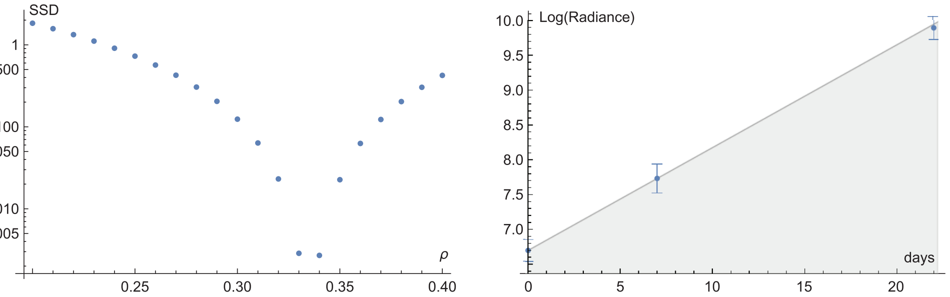


Figure S.2: Fitting of cancer cell proliferation rate, $\rho$. *Left*, sum of squared deviations (SSD) of logarithms of radiance between simulations and experimental data under variation of $\rho$. *Right*, simulation with optimal $\rho = 0.34$ along with the experimental data of control mice group.

##### S.1.3.4 Cancer cells radiosensitivity to unanchored nuclides

To estimate radiosensitivity of cancer cells to unanchored nuclides, we rely on the data of experiment with trastuzumab, which does not bind to cancer cells. The model in this case effectively boils down to the following system of equations. Note that the equation for inert antibodies is omitted here, as it does not influence the other variables. Since the coefficient of proportionality between radiance and cell number does not affect the dynamics here, it is set it to unity.

$$\begin{aligned}
&\text{Concentration of active antibodies in plasma:} && a' = \overbrace{\frac{A_1}{V}\cdot\delta(t-t_1)}^{\text{injection}}\overbrace{-\lambda a}^{\text{decay}}\ \overbrace{-\kappa_c a}^{\text{clearance}}\ ;\\
&\text{Number of viable cancer cells:} && N' = \overbrace{\rho N}^{\text{proliferation}}\ \ \overbrace{-k_f\alpha\lambda\cdot aN}^{\text{damage}};\\
&\text{Number of damaged cancer cells:} && D' = \overbrace{k_f\alpha\lambda\cdot aN}^{\text{damage}}\overbrace{-\omega D}^{\text{death}}.
\end{aligned}$$

The radioconjugates were injected once, 1 day after the first imaging. Therefore, only one injection is considered, with $t_1 = 1$ day and $A_1 = 0.0456$ pmol (assuming equal molecular masses of trastuzumab and daratumumab). We are going to fit the value of the product $k_f\alpha$ using the already estimated value of $\rho = 0.34$. However, the values of other parameters are linked with notable uncertainty, except for the radionuclide decay rate $\lambda$, which is a physical constant. At first, we use the basic values for the antibody clearance rate $\kappa_c = 0.1$ and the volume of drug distribution $V = 1$, and further we compare the optimal values of $k_f\alpha$ under their variation in physiologically justified ranges $\kappa_c \in [0.04, 0.28]$ and $V \in [0.75, 1.5]$. As the death of cancer cells should mainly happen upon mitosis, its rate can be expected to be lower than cell proliferation rate. Therefore, we set its variation range as $\omega \in [0.034, 0.34]$ and at first use the basic value of 0.05.

The chosen basic values of these parameters yield the optimal value of $k_f\alpha = 150$, as depicted in Fig. S.3. Under variation of $\kappa_c$, $V$ and $\omega$ the most probable optimal values of $k_f\alpha$ lie in the range $[80, 160]$ (see supplementary code).

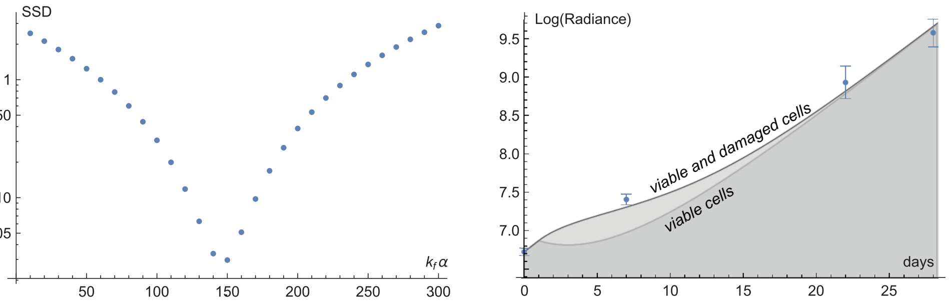


Figure S.3: Fitting of cancer cell radiosensitivity to unanchored nuclides, $k_f\alpha$. *Left*, sum of squared deviations (SSD) between simulated and experimental data under variation of $k_f\alpha$. *Right*, simulation with optimal $k_f\alpha = 150$ along with the experimental data with untargeted $^{225}$Ac-DOTA-trastuzumab.

##### S.1.3.5 Cancer cells radiosensitivity and damaged cells death rate

Further we use the data for the mice groups treated by $^{225}$Ac-DOTA-daratumumab to fit two parameters: cancer cell radiosensitivity, $\alpha$, and damaged cells death rate, $\omega$. Full model is used for this purpose as well as the coefficients of drug impurity, defined in Section S.1.3.2. Note that for the basic set of parameters the total amount of specific receptors on cancer cells at the moment of drug injection is 6.3 pmol. This is more than an order of magnitude less than the total amount of injected daratumumab in each setting. The upper value of the amount of specific receptors on cancer cells is 100 pmol, which is still twice less that the amount of used daratumumab. This suggests that cancer cells did not have enough capacity to attract all the injected drug, and its considerable fraction should have remained in the bloodstream during experiments. According to Fig. S.1, at lower doses cancers continued to grow, suggesting that they may have eventually attracted the major part of the injected drug. At higher doses it is likely that not enough new receptors were produced for that.

Like previously, the values of many other parameters are linked with uncertainties. At first we use the basic values of parameters, provided in Table 1 of main text, and $k_f \alpha = 150$, as estimated in previous section.

Since the death rate of damaged cells $\omega$ is expected to notably affect cell dynamics only in the high-dose settings, we at first use the 300 nCi and 600 nCi experiments to estimate $\omega$. Figure S.4 shows the discrepancy between experimental data and simulation results under variation of $\alpha$ and $\omega$, with red dots marking the optimal pairs of values: $(403, 0.043)$ for 300 nCi and $(660, 0.05)$ for 600 nCi. Based on that, we further use the basic value of $\omega = 0.05$ and we allow its further variation by an order of magnitude higher and lower.

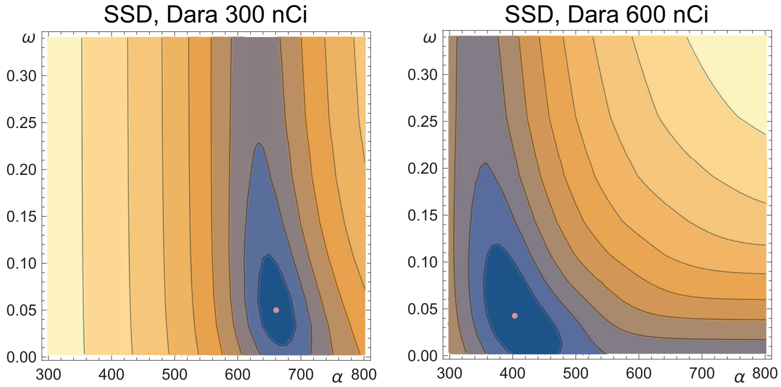


Figure S.4: Fitting of cancer cell radiosensitivity, $\alpha$, and damaged cancer cells death rate, $\omega$, in high-dose settings. SSD=sum of squared deviations. Pink dots mark the optimal pairs of values.

These results indicate that the experimental data for these mice groups are better reproduced with the use of notably different values of cancer cell radiosensitivity. Fitting of the data for each group of mice treated with daratumumab further exacerbates this fact, as Fig. S.5 shows. The upper left plot demonstrates notable decrease of fitted value of cancer cell radiosensitivity under the increase of injected dose. The optimal values of $\alpha$ in 25 nCi and 600 nCi settings differ more than ten-fold under the basic values of other parameters.

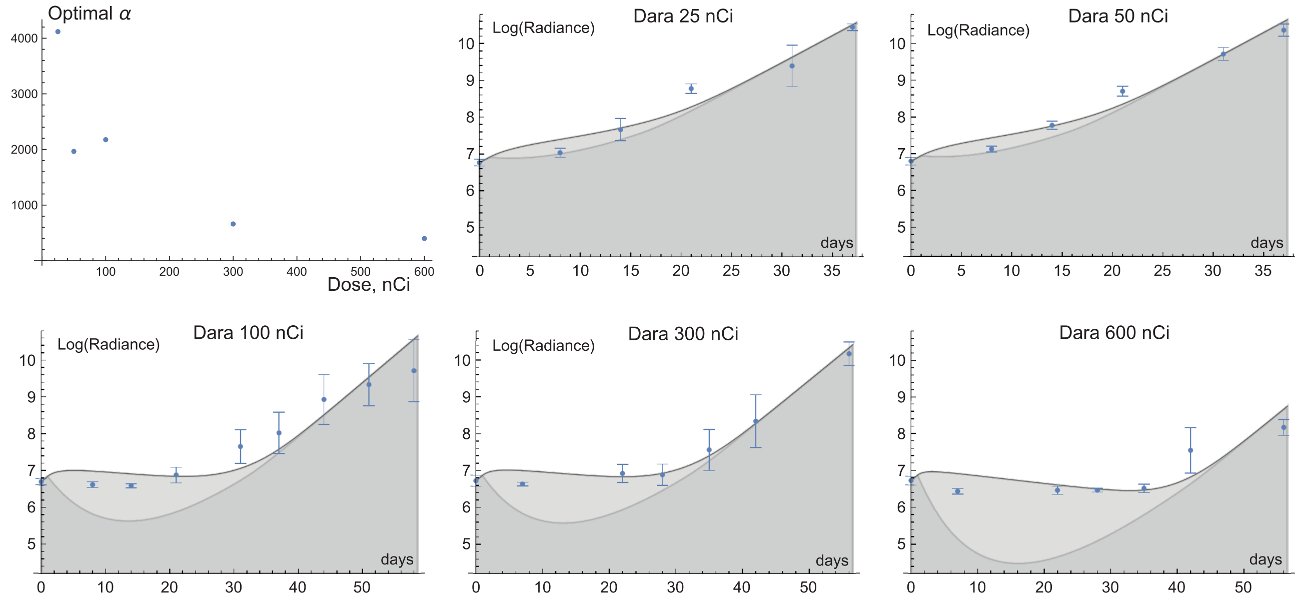


Figure S.5: Fitting of cancer cell radiosensitivity, $\alpha$, to experimental data with $^{225}$Ac-DOTA-daratumumab. Upper left, optimal values of $\alpha$ for experiments with different doses. Other plots, simulations with corresponding optimal values of $\alpha$ along with the experimental data.

This pattern of change of optimal $\alpha$ with injected dose persists under variation of values of other model parameters. Global variation of $\kappa_c$, $\kappa_p$, $\gamma$, $V$, $k_s$, $N_0$ and $k_f\alpha$ yielded ≈9.6 as the average ratio of optimal $\alpha$ in 25 nCi and 600 nCi settings, with its range being $[7, 12]$ (see supplementary code).

Therefore, the homogeneous cancer model is not sufficient to reasonably approximate the experimental data with a single set of parameters. A notable heterogeneity of cancer cells response to treatment is present, which will be discussed further, and the augmentation of the current model will be introduced. However, the current model still holds value for theoretical investigation, which is performed within this study. For it, we choose the basic value of $\alpha$ to be equal to 500 and allow its variation by an order of magnitude higher and lower. Since the significance of unanchored nuclides decays was estimated in the previous section on the basis of experimental data where cancer growth was perturbed comparably slightly, cancer cells dynamics there should correspond to rather high radiosensitivity. Given that, we choose the basic value for $k_f = 150/3000 = 0.05$ and allow its variation in the range $[0.01, 0.25]$ for numerical investigation.

### S.2 Single-dose treatment by pure radioconjugates

Computational codes for this section are provided in supplementary file '02-Pure-drug.nb".

#### S.2.1 Analytical investigation of the model in close-to-curative setting under low amount of injected radioconjugates

For the basic set of parameters, the total amount of specific receptors on cancer cells is $\gamma N_0 = 6.3$ pmol. The same amount of pure radioconjugates corresponds to ≈83,000 nCi of activity, which exceeds the greater dose used in our experiments by more than two orders of magnitude. If the amount of injected antibodies is significantly less than $\gamma N_0$, an approximated version of the model can be derived which behavior is similar to that of the full model. It allows for analytical estimations of several crucial measures for close-to-curative settings. By this term we imply that the minimum number of viable cancer cells achieved during treatment is much lower than their initial number: $N_m << N_0$. At that, the cancer cure probability, e.g., determined via the classical equation $e^{-N_m}$ [9], does not have to be high by itself. The formulas derived in this section can provide good approximations even for the cases with the minimum viable cancer cell numbers of the order of hundreds of thousands, that corresponds to absolutely negligible cure probabilities. For comparison of analytical formulas with simulation results in this section we consider a single injection of pure radioconjugates with $A = 50$ nCi of activity at $t_0 = 0$.

##### S.2.1.1 Estimation of total amount of toxic decays

Due to high antibody-receptor binding rate, the vast majority of the injected drug binds to specific receptors very rapidly. The amount of activity, that is lost due to clearance and decays in blood, can be estimated from the relative rates of the three exponentially decaying processes comprising antibodies dynamics:

$$\frac{\kappa_c + \lambda}{\kappa_c + \lambda + k_{on}\gamma N_0/V} \cdot A,$$

which is ≈0.12 nCi for basic parameter set. This estimation is only ≈0.4% greater than the simulation result.

So, we can make a reasonable approximation that at $t_0 = 0$ all the drug injected at this moment instantaneously occupies cancer cell receptors, setting

$$f_{AN}(0) = f_{AD}(0) \approx f_0 \equiv \frac{A}{\gamma N_0}.$$

For basic set of parameters, it yields $f_0 \approx 0.0006$, which is ≈2% greater than the maximum value of $f_{AD}$ attained in the simulation.

The amount of radioactivity lost due to clearance and decays of antibody fragments can be estimated analytically for close-to-curative setting, in which the vast majority of cancer cells eventually die. A reasonable approximation for this purpose is that all cancer cells become damaged starting from $t = 0$. Thus, $D(t) \approx N_0 e^{-\omega t}$ and $f_{AD}(t) \approx f_0 e^{-\lambda t}$. Approximated equation for the concentration of active fragments in blood therefore is:

$$p' \approx \omega \frac{A}{V} e^{-[\lambda+\omega]t} - [\lambda + \kappa_p]p.$$

Given that $p(0) = 0$, it is solved as

$$p(t) \approx \frac{\omega}{\kappa_p - \omega} \frac{A}{V} \cdot \{e^{-[\omega+\lambda][t-t_0]} - e^{-[\kappa_p+\lambda][t-t_0]}\}.$$

The amount of radioactivity that is lost due to decays in blood is $V \cdot \lambda \int_0^T p(t)\mathrm{d}t$, which is equal to

$$\frac{\omega \cdot \lambda}{[\lambda + \omega][\lambda + \kappa_p]} \cdot A.$$

For the basic parameter set this is ≈1.4 nCi, which is only ≈3% greater than the simulation result.

The estimated value of the total amount of toxic radioactivity, spent during decays in blood is as follows, being proportional to the injected radioactivity:

$$\lambda A \cdot \Big\{ \frac{1}{\lambda + \kappa_c + k_{on}\gamma N_0/V} + \frac{\omega}{[\lambda + \omega][\lambda + \kappa_p]} \Big\}. \tag{S.1}$$

###### S.2.1.2 Simplification of the system for the analysis of cancer cell dynamics

For analytical estimations, regarding the viable cancer cell dynamics and therefore related to the curative effect of a single injection, we can continue using the approximation that all the injected drug instantaneously occupies cancer cell receptors. Thus, the equations for antibodies can be discarded, and the terms including $a$ and $b$ can be nullified. That renders the equations for free receptors redundant as well. Let's also notice that the radiation damage due to the unanchored nuclides is negligible in the corresponding simulation. In it, after the antibodies are bound, the maximum value of the term of dose from unanchored nuclides is more than four orders of magnitude smaller than the maximal sum of the other terms of radiation damage function. This suggests their negligibility under parameters variation as well. Since this assumption leads to the fact that antibody fragments do not influence cancer cells, their equations can be omitted as well.

The term of cross-fire can be simplified due to the following observations. During the first hours of treatment, when damaged cells begin to emerge, the fractions of active receptors of viable and damaged cells, $f_{AN}$ and $f_{AD}$, are close to each other. Therefore, they can be approximated as one variable, taken out of the brackets. After already one day, the number of viable cells is very small compared to damaged cells number and thus can be neglected. Therefore, we can approximate the total number of active receptors of cancer cells as $f_{AD}[N + D]$, at least from the moment of drug injection until the minimal number of viable cancer cells is achieved.

Recall from Section S.1.2, that the processes that active receptors participate in are: binding, which we neglect in this approximation; decay; transfer from viable to damaged cells, which does not change their total number; and degradation with cell death. Overall, this leads to the following approximate equation:

$$(f_{AD}[N + D])' = -\lambda f_{AD}[N + D] - \omega f_{AD} D.$$

All the approximations result in the following system. For convenience we introduce $\chi \equiv \alpha\lambda\gamma/\nu$, which corresponds to the maximal possible rate of damage of cancer cells due to irradiation from nuclides anchored to them, which is achieved when all their receptors are occupied by radioconjugates.

$$
\begin{aligned}
&\text{Number of viable cancer cells:} && N' = \overbrace{\rho N}^{\text{proliferation}} \overbrace{-R_D(f_{AN}, f_{AD}) \cdot N}^{\text{damage}};\\
&\text{Number of damaged cancer cells:} && D' = \overbrace{R_D(f_{AN}, f_{AD}) \cdot N}^{\text{damage}} \overbrace{-\omega D}^{\text{death}};\\
&\text{Fraction of active receptors of viable cancer cells:} && f'_{AN} = -[\lambda + \rho] f_{AN};\\
&\text{Fraction of active receptors of damaged cancer cells:} && f'_{AD} = -\lambda f_{AD} - \frac{\rho N}{N + D} f_{AD};\\
&\text{Radiation damage function:} && R_D(f_{AN}, f_{AD}) = \chi \cdot \left\{ \overbrace{k_S \cdot f_{AN}}^{\text{self-damage}} \overbrace{+[1 - k_S] \cdot f_{AD}}^{\text{cross-fire}} \right\}.
\end{aligned}
\tag{S.2}
$$

For the analysis of this system, it is convenient to consider at first the opposite extreme cases regarding the relative significance of self-damage and cross-fire.

###### S.2.1.3 The case of negligible cross-fire, $k_s = 1$ (self-damage-only)

In case when only self-damage contributes to curative effect of treatment ($k_s = 1$), Eqs. (S.2) boil down to

$$
\begin{aligned}
N' &= \rho N - \chi f_{AN} N;\\
f'_{AN} &= -[\lambda + \rho] f_{AN}.
\end{aligned}
\tag{S.3}
$$

The equation for the damaged cells can be omitted here since they do not affect the dynamics of viable cells. The equation for active receptors of viable cancer cells is independent and can be solved as

$$
f_{AN}(t) = \frac{A}{\gamma N_0} e^{-[\lambda+\rho]t}.
$$

That leads to the analytical solution for the number of viable cancer cells:

$$
N^{SD}(t) = N_0 \cdot exp(\rho t - \frac{\alpha}{\nu} \cdot \frac{\lambda}{\lambda + \rho} \cdot \frac{A}{N_0}[1 - e^{-[\lambda+\rho]t}]),
$$

where superscript $SD$ highlights the validity of this formula for the self-damage-only case. When the derivative of this function is zero, the minimal number of viable cancer cells $N_m^{SD} = N^{SD}(t_m^{SD})$ is achieved. That yields

$$
t_m^{SD} = \frac{ln(\frac{\alpha d}{P})}{\lambda + \rho}, \quad N_m^{SD} = N_0 \cdot [\frac{\alpha d}{P}]^{\frac{P}{1+P}} \cdot e^{\frac{P - \alpha d}{1+P}}, \quad \text{where } d \equiv \frac{A}{\nu N_0}, \quad P = \frac{\rho}{\lambda}.
\tag{S.4}
$$

The case when $\alpha d = P$ and therefore $A = \rho\nu N_0/[\alpha\lambda]$ corresponds to the situation in which $t_m^{SD} = 0$ and therefore $N'(0) = 0$, i.e., the border case for which the minimum number of viable cells is $N_m^{SD} = N_0$. Therefore, in this case the number of viable cells does not decrease at all during treatment. The form of Eqs. (S.2) suggests that this notion is valid for any value of $k_s$. For the basic parameter set the corresponding dose is ≈5.75 nCi. Thus, at close and greater doses the used approximation of close-to-curative setting loses its validity.

For the basic set of parameters with $A = 50$ nCi and $k_s = 1$, $t_m^{SD} \approx 5.3$ days, less than 0.04% greater than the actual simulation result, and $N_m^{SD} \approx 305772$ cells, which is only ≈5% less than the actual result.

Let's find the minimal curative dose for this case, i.e., the amount of injected active antibodies $A_{cur}^{SD}$ that would correspond to $N_m^{SD} = N_{cur}$. This can be done with the use of Lambert function $W(z)$, which is defined as the solution of equation $W(z)e^{W(z)} = z$. Generally, this function has two branches for real numbers, from which the solution has to be chosen on reasonable ground. After straightforward transformations we obtain

$$
\alpha d_{cur}^{SD} = -P \cdot W_{-1}(-e^{-1} \cdot [\frac{N_{cur}}{N_0}]^{[1+P]/P}), \quad \text{thus} \quad A_{cur}^{SD} = -N_0 \cdot \frac{\rho}{\lambda} \cdot \frac{\nu}{\alpha} \cdot W_{-1}(-e^{-1} \cdot [\frac{N_{cur}}{N_0}]^{[\lambda+\rho]/\rho}),
$$

where $W_{-1}$ branch is chosen, since $W_0$ would always provide $\alpha d < P$, which, as follows from the above reasoning, would not lead to any decrease of viable cancer cell number.

For the basic set of parameters with $k_s = 1$, $A^{SD}_{cur} \approx 176.8$ nCi, which is less than 0.7% smaller than the simulation result. Thus, this estimation provides a good degree of approximation.

From these estimations we can obtain the minimal number of molecules of radioconjugates, that should anchor on each cancer cell in order to achieve cure. It is $M_m = A^{SD}_{cur}/N_0$, where the dose is expressed in the number of molecules of injected antibodies. That is ≈272 molecules for the basic set.

###### S.2.1.4 The case of negligible self-damage, $k_s = 0$ (cross-fire-only)

In the case when only cross-fire contributes to the curative effect, Eqs. (S.2) boil down to:

$$\begin{aligned} N' &= \rho N - \chi f_{AD} N; \\ D' &= \chi f_{AD} N - \omega D; \\ f'_{AD} &= -\lambda f_{AD} - \frac{\rho N}{N+D} f_{AD}. \end{aligned}$$

Although this model is not solved analytically, the minimal number of viable cancer cells can be reasonably estimated, without obtaining an explicit formula for $N(t)$. For this goal, the second term of the equation for $f_{AD}$ has to be approximated. We can use the fact that during the initial period of active damaging of cancer cells their total number does not change drastically – e.g., in the basic set simulation with $A = 50$ nCi it increases by only about 10%. So, let's approximate $N(t) + D(t) \approx N_0$. Then, disregarding the equation for the damaged cells as effectively not affecting the viable cells, we get:

$$\begin{aligned} N' &= \rho \cdot N - \chi \cdot N f_{AD}; \\ f'_{AD} &= -\lambda \cdot f_{AD} - \frac{\rho}{N_0} \cdot N f_{AD}. \end{aligned}$$

Let's represent this system in the following form, where $T = \lambda t$, $\tilde{N} = \frac{\rho}{\lambda} \cdot \frac{N}{N_0}$, $\tilde{f} = \frac{\chi}{\lambda} \cdot f_{AD}$, $P = \frac{\rho}{\lambda}$:

$$\begin{aligned} \frac{d\tilde{N}}{dT} &= P \cdot \tilde{N} - \tilde{f}\tilde{N}; \\ \frac{d\tilde{f}}{dT} &= -\tilde{f} - \tilde{f}\tilde{N}. \end{aligned}$$

Equating $dT$, we get:

$$dT = \frac{d\tilde{N}}{\tilde{N}[P - \tilde{f}]} = \frac{d\tilde{f}}{-\tilde{f}[1 + \tilde{N}]},$$

or

$$\frac{[1 + \tilde{N}]}{\tilde{N}} d\tilde{N} = \frac{[\tilde{f} - P]}{\tilde{f}} d\tilde{f}.$$

After integrating:

$$ln(\tilde{N}) + \tilde{N} = E + \tilde{f} - P \cdot ln(\tilde{f}),$$

where $E$ is a conserved quantity, which can be found via initial values of $\tilde{N}_0$ and $\tilde{f}_0$:

$$E = ln(\tilde{N}_0 \tilde{f}_0^P) + \tilde{N}_0 - \tilde{f}_0 = ln(\frac{\rho}{\lambda} \cdot [\alpha \cdot \frac{A}{\nu N_0}]^{\rho/\lambda}) + \frac{\rho}{\lambda} - \alpha \cdot \frac{A}{\nu N_0}.$$

After exponentiating:

$$\tilde{N} e^{\tilde{N}} = \tilde{f}^{-P} e^{E + \tilde{f}},$$

from which we can express one variable through another, once again using the Lambert function:

$$\tilde{N}(\tilde{f}) = W(\tilde{f}^{-P} e^{E + \tilde{f}}).$$

Since the argument of this function is positive, it is single valued.

The minimum number of viable cancer cells is achieved when the derivative of this function by $\tilde{f}$ is zero:

$$\frac{\tilde{f}-P}{\tilde{f}} \cdot \frac{W(\tilde{f}^{-P}e^{E+\tilde{f}})}{1+W(\tilde{f}^{-P}e^{E+\tilde{f}})} = 0,$$

which happens when and only when $\tilde{f} = P$. At that point

$$\tilde{N}_m = W(P^{-P}e^{E+P}).$$

Therefore:

$$N_m^{CF} = \frac{N_0}{P} \cdot W(P \cdot [\frac{\alpha d}{P}]^P \cdot e^{2P-\alpha d}), \quad \text{where as previously } d \equiv \frac{A}{\nu N_0}, \quad P \equiv \frac{\rho}{\lambda}, \tag{S.5}$$

where superscript $CF$ highlights the validity of this formula for the case of negligible self-damage. In the close-to-curative case, we expect $\alpha d$ to be notably higher than $P$ (recall that $\alpha d = P$ corresponds to the case with no decrease of viable cancer cell number during treatment). Therefore, the argument of the Lambert function is expected to be very small and we can approximate this function by the first term of its Taylor expansion:

$$N_m^{CF} = N_0 \cdot [\frac{\alpha d}{P}]^P \cdot e^{2P-\alpha d}.$$

For the basic set of parameters with $k_s = 0$ and $A = 50$ nCi, $N_m^{CF}$ ≈0.008 cells, with expanding of Lambert function resulting in negligible change. This is ≈5% less than the corresponding simulation result.

Although the explicit function for $N(t)$ cannot be obtained, the moment of achieving minimum viable cancer cell number can be estimated using the approximation that neglects the quadratic term in the equation for active receptors. Then the model takes the form similar to Eqs. (S.3) and it can be deduced that

$$t_m^{CF} \approx ln(\frac{\alpha d}{P})/\lambda,$$

which is ≈30.9 days for the basic set, ≈7% greater than the simulation result.

The minimal curative dose, $A_{cur}^{CF}$, can be estimated similarly to the previous section:

$$\alpha d_{cur}^{CF} = -P \cdot W_{-1}(-e^{-2} \cdot [\frac{N_{cur}}{N_0}]^{1/P}), \quad \text{thus} \quad A_{cur}^{CF} = -N_0 \cdot \frac{\rho}{\lambda} \cdot \frac{\nu}{\alpha} \cdot W_{-1}(-e^{-2} \cdot [\frac{N_{cur}}{N_0}]^{\lambda/\rho}).$$

For the basic set of parameters with $k_s = 0$, $A_{cur}^{CF} \approx 49.8$ nCi, which is only ≈0.1% smaller than the simulation result. The minimal number of radioconjugate molecules, that should anchor on each cancer cell in order to achieve cure, is about 76 in this case.

Notably, the analytical formulas for $A_{cur}^{CF}$ and $A_{cur}^{SD}$ are similar, and there are only two differences in them. They are manifested in different forms of exponents within the Lambert function. For self-damage-only case the fraction $[\lambda + \rho]/\rho$ reflects the decrease of self-damage efficiency in result of both decay of nuclides and proliferation of cancer cells. In cross-fire-only case after a transient period the rate of energy deposition decreases independently of cell proliferation, therefore the fraction $\lambda/\rho$ is present in this case. Cell proliferation, however, still affects the formula for $A_{cur}^{CF}$, yielding the factor $e^{-2}$ rather than $e^{-1}$ as in self-damage-only case. Neglect of this alteration would lead to ≈13% error in estimation of curative dose.

With increase of $N_0$, the minimal curative dose grows faster than a linear function, which is valid for both $k_s = 0$ and $k_s = 1$. Therefore, a delay of treatment beginning will cause the required minimal single curative dose to grow even faster than the number of cancer cells, which is by itself an exponential function.

###### S.2.1.5 Influence of nuclide decay rate on minimal surviving fraction of cells

As Eqs. (S.4) and (S.5) suggest, the estimated minimal surviving fractions of cancer cells in both cross-fire-only case, $N^{CF}/N_0$, and self-damage-only case, $N^{SD}/N_0$, effectively depend on only two parameters. One of them is $P = \rho/\lambda$, the ratio of cell proliferation rate and nuclide decay rate. When it tends to zero, both formulas tend to $e^{-\alpha \cdot A/[\nu N_0]}$. This value corresponds to the classical formula for instantaneous irradiation, widely used in external beam setting, since $A$ here corresponds to the total activity, affecting cancer cells, while $\nu N_0$ represents the volume occupied by them, and their ratio can be straightforwardly converted into the energy deposited per unit of cancer mass, measured in Grays.

The dependence of minimal surviving fraction of cancer cells on $\rho/\lambda$ is illustrated in Fig. S.6. As it shows, treatment efficacy grows both with the decrease of cancer cell proliferation rate $\rho$ and with the decrease of nuclide half-life. Analytical estimations correspond well to numerical simulations of the full system under variation of $\lambda$ until nuclide decay rate becomes very fast, so that a large fraction of injected nuclides decay in the bloodstream before anchoring to cancer cells.

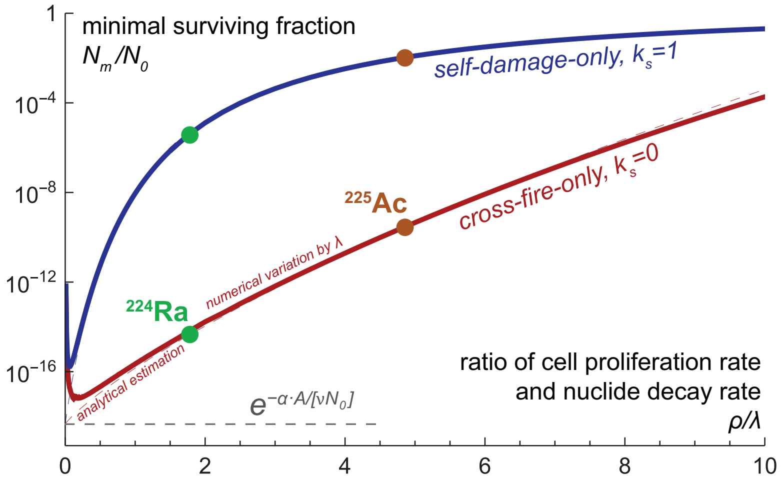


Figure S.6: Dependence of minimal surviving fraction of cancer cells on the ratio of cell proliferation rate and nuclide decay rate.

These results suggests that the negative influence of cancer cell repopulation can be mitigated by faster decays of nuclides. From this perspective, the use of radium isotope $^{224}$Ra, which decay chain yields $\approx$2.5 faster release of $\alpha$-particle energy, comparable to that of $^{225}$Ac, represents a potentially more effective option for radiopharmaceutical therapy of blood cancers [10].

##### S.2.1.6 The case of non-negligible self-damage and cross-fire, $0 < k_s < 1$

The more general case when both self-damage and cross-fire have notable effect is not amendable to the types of analytical investigation used above for the extreme cases. Additional reasoning nevertheless allows to deduce the analytical function $A_{cur}(k_s)$, that approximates the minimal curative dose for a general value of $k_s$.

First of all, it is obvious that it should coincide with the obtained formulas for the border cases $k_s = 0$ and $k_s = 1$. Let's represent them the following way:

$$\begin{aligned} A_{cur}(0) = \quad & A_{cur}^{CF} = -N_0 \cdot \frac{\rho}{\lambda} \cdot \frac{\nu}{\alpha} \cdot W_{-1}(-e^{-1} \cdot [\frac{N_{cur}}{N_0}]^{\frac{\lambda}{\rho}} \cdot e^{-1}), \\ A_{cur}(1) = \quad & A_{cur}^{SD} = -N_0 \cdot \frac{\rho}{\lambda} \cdot \frac{\nu}{\alpha} \cdot W_{-1}(-e^{-1} \cdot [\frac{N_{cur}}{N_0}]^{\frac{\lambda}{\rho}} \cdot \frac{N_{cur}}{N_0}). \end{aligned} \tag{S.6}$$

From the fact that $N_{cur}/N_0 << e^{-1}$ it follows that $A_{cur}(1)$ is always greater than $A_{cur}(0)$. It also follows from Eqs. (S.6), that $A_{cur}(1)/A_{cur}(0)$ grows monotonically with the increase of $\rho$ (see supplementary code).

Since $[1-k_s]$ participates as a multiplier in the cross-fire term, it can be expected that in the limit of $k_s \to 0$ cross-fire effect by itself would demand scaling of the minimal curative dose as $A_{cur}^{CF}/[1-k_s]$. This function however has asymptotic behavior at $k_s \to 1$, and thus it is expected to significantly diverge from the numeric result as $k_s$ increases, since this function completely neglects the effect of self-damage. A way to minimize this discrepancy is to use the value of $A_{cur}^{SD}$ for rescaling this function the following way:

$$A_{cur} = A_{cur}^{CF}/[1 - k_s \cdot \{1 - \frac{A_{cur}^{CF}}{A_{cur}^{SD}}\}], \tag{S.7}$$

thus forcing it to be equal to $A^{SD}_{cur}$ at $k_s = 1$. This function provides a descent approximation, as Fig. S.7 shows. At greater values of $k_s$ it however goes notably below the the numerical curve. The relative error, maximal by its absolute value, is ≈-5%, which is attained at $k_s \approx 0.89$. For $k_s < 0.5$ the correspondence of analytical estimation to the numerical results is better. The maximum error in that region is only ≈0.8%, at $k_s \approx 0.29$.

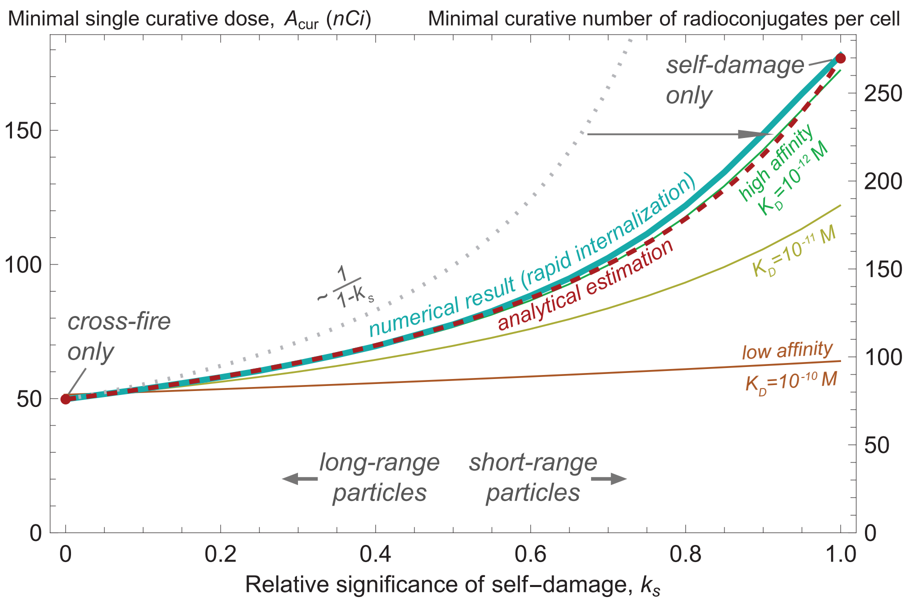


Figure S.7: Dependence of minimal single curative dose $A_{cur}$ on the particle range for the basic set of parameters. Analytical curve is governed by Eqs. (S.6)-(S.7). Thin curves refer to the case of non-internalizing antibody with varying affinity.

For a more general case, especially under moderate initial number of specific receptors and/or fast antibody clearance, the estimation of minimal single curative dose can be improved by accounting for the fact that only a fraction of injected drug anchors to cancer cells, and therefore by multiplying Eq. (S.7) by

$$\Omega = \frac{\kappa_c + \lambda + k_{on}\gamma N_0/V}{k_{on}\gamma N_0/V}.$$

Importantly, for these results we neglect the unbinding of antibodies, which is an assumption justified by the fact that daratumumab is internalized in multiple myeloma cells within only several hours [11]. Figure S.7 also illustrates the cases for which the used antibodies are assumed to not internalize and to bind to specific receptors reversibly. The corresponding graphs are produced during simulations of the following augmentation of the full model, where ellipses stand for its original terms.

$$a' = ... + k_{off}\frac{\gamma}{V}[f_{AN}N + f_{AD}D], \quad b' = ... + k_{off}\frac{\gamma}{V}[\{1 - f_{FN} - f_{AN}\}N + \{1 - f_{FD} - f_{AD}\}D],$$
$$f'_{FN} = ... + k_{off}[1 - f_{FN}], \quad f'_{AN} = ... - k_{off}f_{AN},$$
$$f'_{FD} = ... + k_{off}[1 - f_{FD}], \quad f'_{AD} = ... - k_{off}f_{AD}.$$

The range of particles becomes less significant with decrease of antibodies affinity, characterized by dissociation constant $K_D = k_{off}/k_{on}$, since the redistribution of radioconjugates from damaged to viable cells mitigates the decrease of nuclide density on them caused by cell proliferation.

#### S.2.2 Global parameter sweep

##### S.2.2.1 Curative dose and toxicity: highly idealized *in silico* trial

For the sake of mathematical curiosity, let's regard the global parameter sweep as a part of the following problem. Let's assume that the cancer dynamics for each mouse can be exactly represented via the developed model, and for each mouse all the corresponding individual parameter values are known. The task is to cure each mouse by using one injection of pure radioconjugates, with keeping treatment toxicity as low as possible.

In theory, the setting of this problem can be further gradually deidealized and brought closer to a realistic human scenario. At that, the solution of this highly idealized problem should serve as the foundation for the guiding lines of treatment administration suggested for a clinical setting.

We put the current problem in the context of *in silico* trial. Below we use the results of the global parameter sweep as the outcome of investigation with a training set of virtual mice. Then we validate the obtained results with a test set of virtual mice. Such conceptual method will be further repeatedly used throughout this work.

For parameter sweep one thousand sets of values of ten parameters – $\kappa_c$, $\kappa_p$, $\gamma$, $V$, $k_s$, $\rho$, $\omega$, $\alpha$, $k_f$, $N_0$ – were selected randomly within the ranges designated in Table 1 of main text, with $\eta$ kept to be equal to 0 to correspond to the injections of pure drug. Importantly, for the designated ranges of parameters the injected amount of antibodies for the minimal single curative dose is always less than the total amount of receptors of cancer cells. In the performed simulations the maximum receptors occupancy of damaged cancer cells did not exceed 10.5%, the average value being about only about 0.3%.

*Minimal single curative dose.* Upper graphs in Fig. S.8 summarize the influence of parameter variation on the values of minimal single curative dose $A_{cur}$, defined numerically by straightforward bisection method.

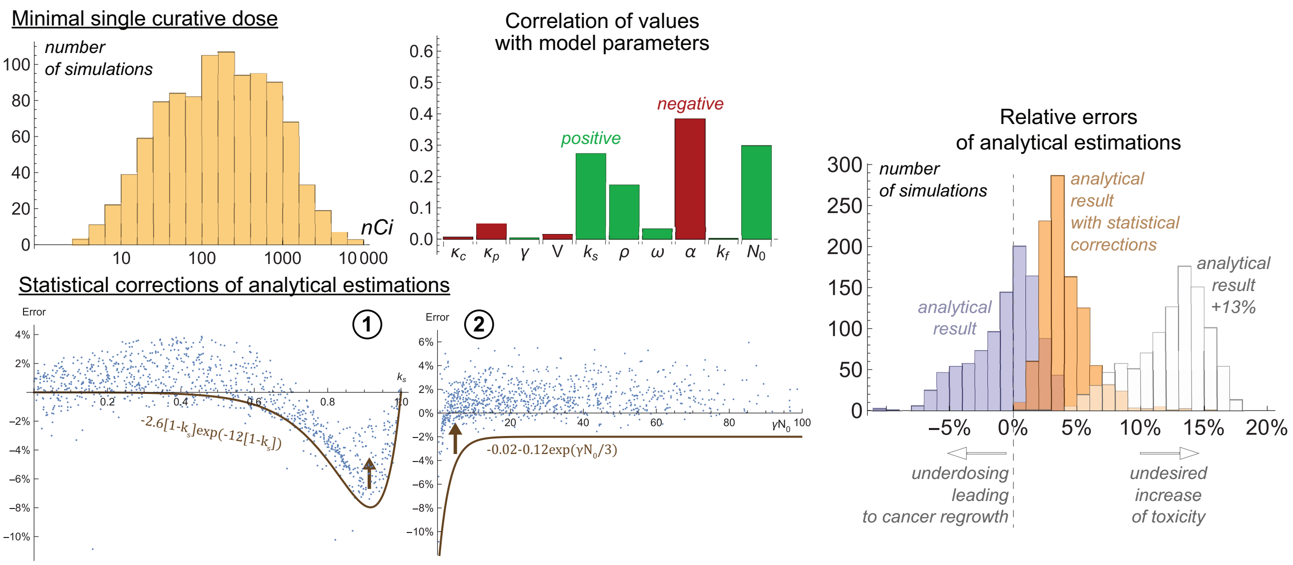


Figure S.8: *Upper*, influence of parameter variation on the minimal single curative dose of pure radioconjugates. *Lower*, illustration for the two-step correction of analytical estimation of minimal single curative doses, leading to absence of underestimations. *Right*, accuracy of analytical estimations without and with statistical corrections.

Performed simulations yield a wide range of minimal single curative doses, spanning more than three orders of magnitude. As correlation chart shows, the four crucial parameters, that determine the value of $A_{cur}$ are cancer cells radiosensitivity, $\alpha$; initial number of cancer cells, $N_0$; relative significance of self-damage, $k_s$; and cancer cells proliferation rate, $\rho$. This corresponds well to the analytical results. Importantly, these numerical results alone are insufficient to infer the nonlinear type of functions, provided in Eqs. (S.6). They can be derived only by analytical investigation.

Analytical formulas provide both underestimations and overestimations of curative doses for different virtual mice. From the treatment point of view, a slight overestimation does not present a significant problem, since the corresponding values of injected activity still would lead to cancer cure, albeit with a slight increase in the accompanying toxic effects, compared to treatment by $A_{cur}$. Underdosing, however, would be a fatal mistake yielding cancer regrowth, and therefore it is desirable to avoid it. A straightforward way to avoid underdosing is to account for maximum relative error via multiplying the estimated values of $A_{cur}$ by a single coefficient. The generated data suggests that adding 13% to each estimated dose ensures cure of all the considered virtual mice in the training set.

A more personalized approach that allows minimizing toxicity while keeping the treatment curative is introduction of a specific correction coefficient for each mouse. This can be done on statistical basis since the values of errors correlate with some of the model parameters. As the lower left graph shows, the error of formula for minimal single curative dose strongly depends on significance of self-damage, $k_s$, and the obtained data can be used to introduce a statistically based correction. When it is applied, we come up with the new distribution of errors that notably depends on the initial total number of specific receptors of cancer cells, $\gamma N_0$, as the middle bottom graph shows. The remaining error can be corrected by introducing the second correction coefficient.

This procedure overall yields the orange histogram of relative errors in the right plot, which shift to the left compared to the one-size-fits-all approach confirms the decrease of used doses and therefore the decrease of treatment toxicity, while maintaining its curative effect since these corrections eventually yield no underestimations. The obtained analytical formulas with statistical corrections will further be used for the test set of virtual mice. It is however still necessary to evaluate whether the assessed doses can be lethally toxic for some mice. The training set contains four lethal cases. The corresponding mice thus cannot survive after a single-dose treatment, but proper adjustment of dose can prolong mouse survival. Therefore, the injected dose has to be decreased in order to comply to the constraint on toxicity. For this purpose, the amount of toxic decays in blood also has to be estimated. As it was noted previously, it consists of two separate effects.

*Activity released in blood from antibodies.* Since almost all of the injected antibodies bind to receptors rapidly, they release only a small fraction of activity in blood before binding, as the upper row in Fig. S.9 shows. The correlators reflect the fact that this fraction decreases with the increase of the effective rate of binding of drug to cancer cells, which corresponds to the previously derived analytical formula

$$\frac{\lambda}{\lambda + \kappa_c + k_{on}\gamma N_0/V}, \tag{S.8}$$

where antibody clearance rate $\kappa_c$ plays minor role, being small compared to the last term in denominator.

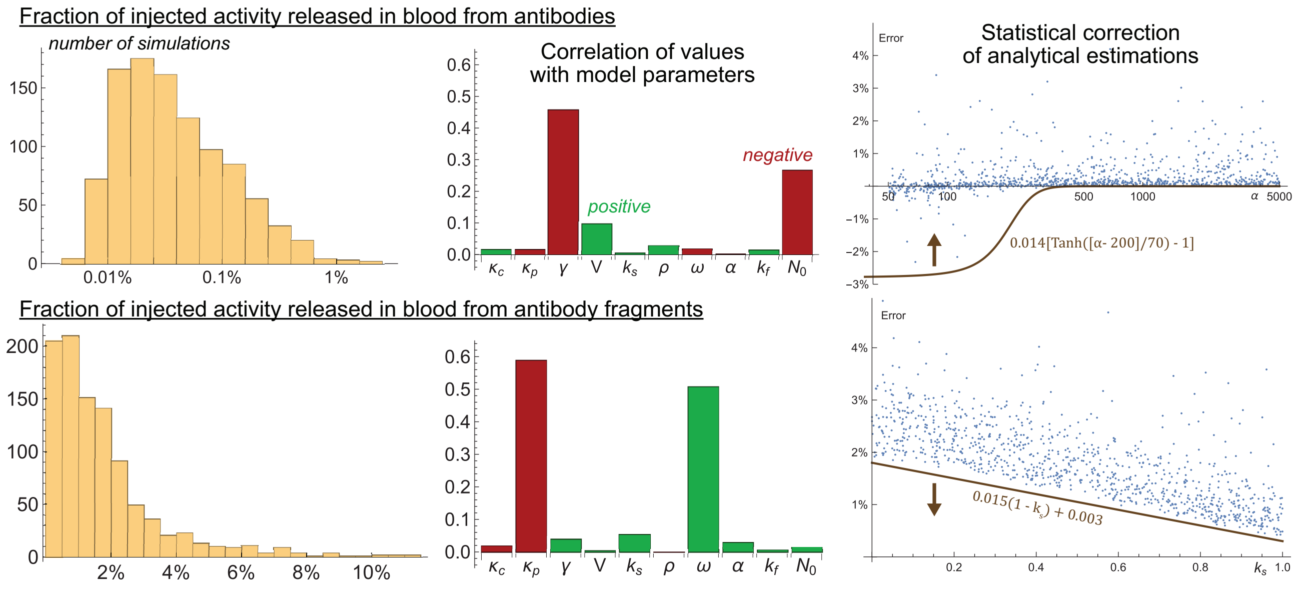


Figure S.9: *Left and middle*, influence of parameter variation on the fraction of injected activity, released in blood (without drug impurities, $\eta = 0$). *Right*, illustrations for the correction of analytical estimations.

Once again, analytical formulas provide both underestimations and overestimations of the actual values of fractions of activity released in blood from antibodies. As well, underestimations have to be avoided, since conducting the treatment with underestimated toxicity level may result in unexpected death. The upper right figure shows that all of the underestimated cases correspond to low values of cancer cell radiosensitivity and illustrates the way to correct the undesired error.

*Activity released in blood from antibody fragments.* This path of activity is generally more significant. Among all the decays in blood in all the simulations of this parameter sweep, ≈97.2% of them happen via this path. The fraction of activity released in blood from antibody fragments has been previously estimated as

$$\frac{\omega \cdot \lambda}{[\lambda + \omega][\lambda + \kappa_p]}, \tag{S.9}$$

with the decisive role of $\omega$ and $\kappa_p$ clearly confirmed by numerical simulations, as the lower row in Fig. S.9 shows.

This formula is obtained under the assumption of immediate damage of effectively all cancer cells. Therefore, it always overestimates the value obtained in numerical simulations. The accuracy of this formula falls with slower killing of cancer cells, which happens with the decrease of the applied dose. Due to that, the analytical error correlates with the values of parameters that determine the value of the curative dose, with the coefficient of self-damage significance $k_s$ being the strongest correlator. The right lower graph illustrates how the overestimation of this part of toxic activity can be minimized depending on the value of this parameter.

*Total activity, released in blood.* Straightforward combination of all the above-discussed analytical formulas and statistical corrections yields the set of predicted amounts of toxic decays in blood, which good correspondence to the actual simulated values is illustrated in Fig. S.10. Notably, the analytical predictions correctly discern the four lethally toxic cases, in which the amount of toxic decays in blood exceeds $A_{bl}^{cr} = 230$ nCi·day.

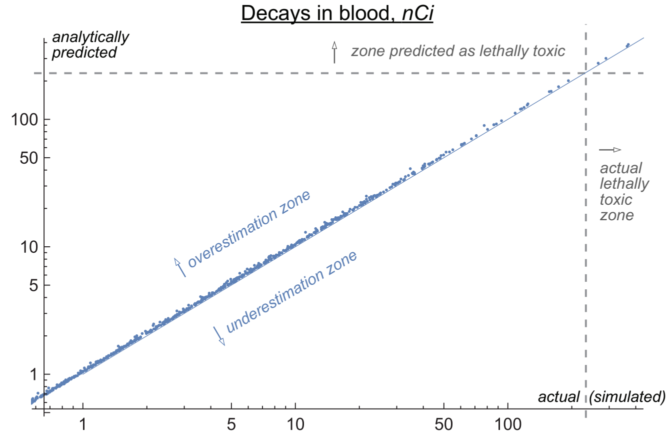


Figure S.10: Simulated versus predicted values of the amounts of decays in blood during parameter sweep for a single-dose treatment by pure radioconjugates.

*Validation of results in a test set of virtual mice.* The obtained formulas and their statistical corrections were used for estimations of personalized doses for one thousand virtual mice in a test set. As Fig. S.11 shows, such approach is only marginally less efficient than straightforward numerical optimization in terms of both survival and toxicity. Due to low toxicity of treatment by pure radioconjugates, administration of a single maximal dose not leading to toxicity-related deaths is also efficient but results in greater overall toxicity.

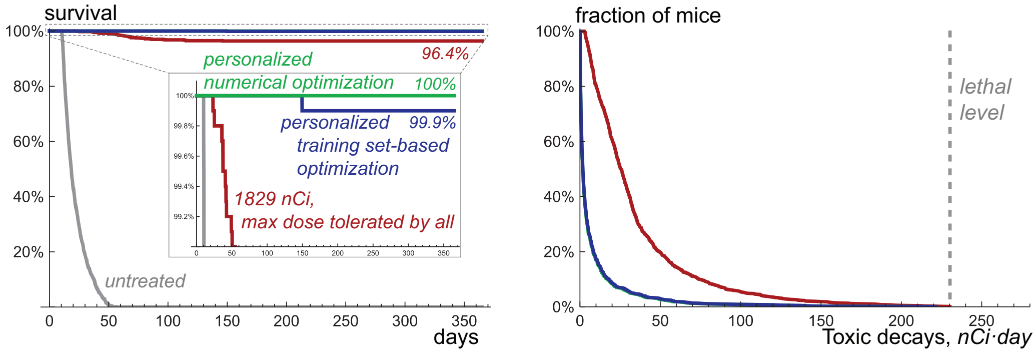


Figure S.11: Results of *in silico* trial for a single injection of pure radioconjugates in the single test set of one thousand virtual mice, treated by: a numerically optimized dose, a dose derived by analysis of a training set, and a single dose, tolerated by all mice. *Left*, survival curves. *Right*, toxicity curves.

Such modeling results may seem unreasonably optimistic, but they are obtained under idealized assumptions that by themselves render the estimation of personalized optimal doses possible and the modeled treatment largely curative. The following sections are devoted to modeling of treatment and suggesting its optimization under sufficiently less idealized conditions, which include the non-purity of used radioconjugates.

The training set generated herein allows analyzing two other treatment-related measures that will be important for that part of the study.

##### S.2.2.2 Other treatment-related measures

*Activity spent on viable cancer cells.* An important parameter related to treatment efficacy is the fraction of activity spent in curative setting on damaging of viable cancer cells rather than on irradiation of already damaged cells or on the loss due to the decays in blood and clearance. This parameter is defined via the following formula:

$$\frac{A_{via}}{A_{cur}} = \Big\{ \overbrace{k_s \cdot \lambda \cdot \int_0^T f_{AN}(t) \cdot \gamma N(t) \mathrm{d}t}^{\text{self-damage}} + \overbrace{[1-k_s]\lambda \cdot \int_0^T [f_{AN}(t) \cdot \gamma N(t) + f_{AD}(t) \cdot \gamma D(t)] \cdot \frac{N(t)}{N(t)+D(t)} \mathrm{d}t}^{\text{cross-fire}}$$
$$\overbrace{+k_f \lambda \cdot \int_0^T \nu N(t) \cdot [a(t) + p(t)] \mathrm{d}t}^{\text{dose from unanchored nuclides}} \Big\} / A_{cur}.$$

As Fig. S.12 shows, this parameters varies in range 0.4%-3.3% in the training set.

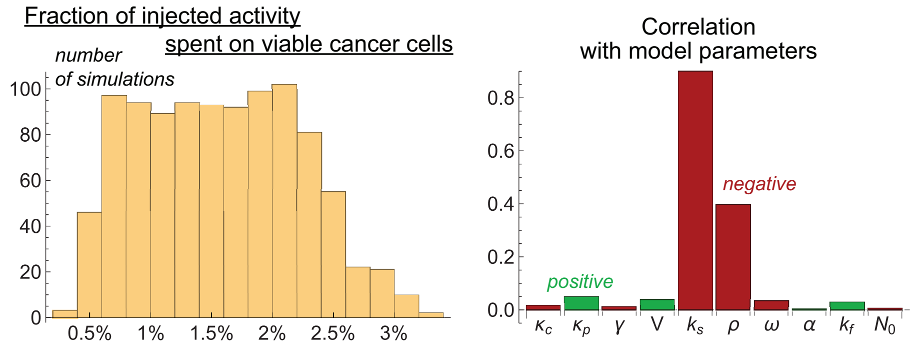


Figure S.12: Influence of parameter variation on the fraction of injected activity spent on irradiation of viable cancer cells after injection of a minimal single curative dose of pure radioconjugates.

Correlation coefficients show that the fraction of activity spent of viable cells drastically decreases with the increase of relative significance of self-damage, $k_s$. In this case the rate of irradiation of viable cells falls rapidly not only due to the decay of nuclides but also due to proliferation of cells. The increase of cell proliferation rate exacerbates this effect, which is also reflected in the correlation chart.

It is hard if possible to estimate analytically the fraction of injected activity spent on viable cancer cells in curative setting. However, it is not difficult to estimate its upper bound for a single-dose setting, corresponding to idealized situation in absence of cell proliferation and death. In this case, the solution of equation for viable cells yields the classical formula for the surviving fraction of cells:

$$\frac{N_m}{N_0} = e^{-\alpha d},$$

where, as previously, $N_m$ is the minimal surviving number of viable cells, and $d$ is total energy released per cancer mass.

Now let's imagine that we gradually increase $d$ starting from zero up to the value that would yield cancer cure, i.e., $d_{cur} = ln(N_0/N_{cur})/\alpha$. As we do it, the viable fraction of cells decreases and increasingly greater

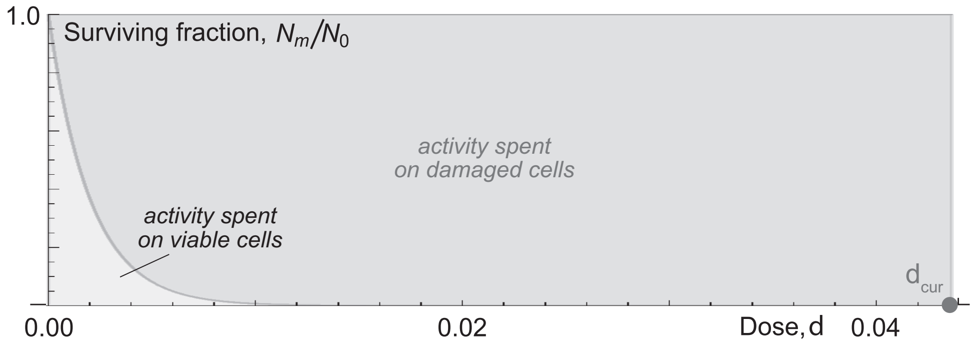


Figure S.13: Illustration for estimation of the upper bound of fraction of injected activity spent on viable cells.

fraction of irradiation is spent on already damaged cells, as Fig. S.13 shows. To obtain the total fraction of activity spent on viable cells, we should use the area under surviving fraction curve from 0 to $d_{cur}$:

$$\frac{A_{via}}{A_{cur}} = \frac{\int_0^{d_{cur}} e^{-\alpha d} \mathrm{d}d}{d_{cur}} = \frac{1 - e^{-\alpha d_{cur}}}{\alpha \cdot d_{cur}} \approx \frac{1}{ln(N_0/N_{cur})}.$$

That is ≈4.6% for the basic values of $N_0$ and $N_{cur}$. Simulations with very low cancer cell proliferation rate and negligible drug clearance yield very close values of the fraction of injected activity spent on viable cells. As was shown above, proliferation of cells, especially under high significance of self-damage, leads to the values notably less than this upper bound.

*Number of newborn cells during treatment.* The results of the training set also allow deriving an approximate formula for the number of cancer cells $N_{new}$, that are born during the treatment by the minimal single curative dose. This measure is important, since newborn cancer cells influence the overall drug binding capacity of cancer. Formally, implicit formulas for $N_{new}$ can be obtained for the approximated systems in the self-damage-only and cross-fire-only settings, introduced above in Sections S.2.1.3-S.2.1.4. However, they would represent the integrals which are hard if possible to take analytically.

Rough estimations of $N_{new}$ can be performed when considering irradiation of cancer cells by constant rate, reflected in the simple system

$$N' = \rho N - \chi f N; \\ f' = 0,$$

where $f$ is the initial fraction of active receptors, estimated as $A_{cur}/[\gamma N_0]$, and $\chi \equiv \alpha\lambda\gamma/\nu$, as defined earlier.

In this case $N_{new}$ can be calculated as

$$N_{new} = \int_0^{\infty} \rho N(t)dt = N_0 \frac{\rho}{\chi f - \rho} = N_0 \cdot \frac{1}{[A_{cur}/N_0] \cdot [\alpha/\nu] \cdot [\lambda/\rho] - 1}.$$

The comparison of estimated and predicted values of $N_{new}$ is provided in Fig. S.14.

This approximation always results in an underestimation of the actual number of newborn cells, since in the full model the rate of irradiation decreases with time. Using Eqs. S.6 for estimation of minimal single curative doses, simplified formulas can be obtained for the cross-fire-only and self-damage-only cases:

$$\begin{aligned} N_{new}^{CF} &= N_0/[-W_{-1}(-e^{-2} \cdot [\tfrac{N_{cur}}{N_0}]^{\frac{\lambda}{\rho}}) - 1], \\ N_{new}^{SD} &= N_0/[-W_{-1}(-e^{-1} \cdot [\tfrac{N_{cur}}{N_0}]^{\frac{\lambda+\rho}{\rho}}) - 1]. \end{aligned} \qquad (S.10)$$

Notably, these formulas are independent of $\alpha$. Therefore, cell radiosensitivity does not influence, to a reasonable degree of approximation, the extent of total proliferation of cancer cells during curative treatment.

The degree of underestimation can be corrected using the training set data, similarly to the previously discussed corrections (see supplementary code). It is important to notice that the ratio $N_{new}/N_0$ varies in a moderate range for the training set, exceeding 0.25 in only three cases and never exceeding 0.5. Therefore, the overall drug binding capacity of cancer during a curative treatment can be generally expected to increase by less than a half. The use of doses greater than $A_{cur}$ will furthermore result in the decrease of this parameter.

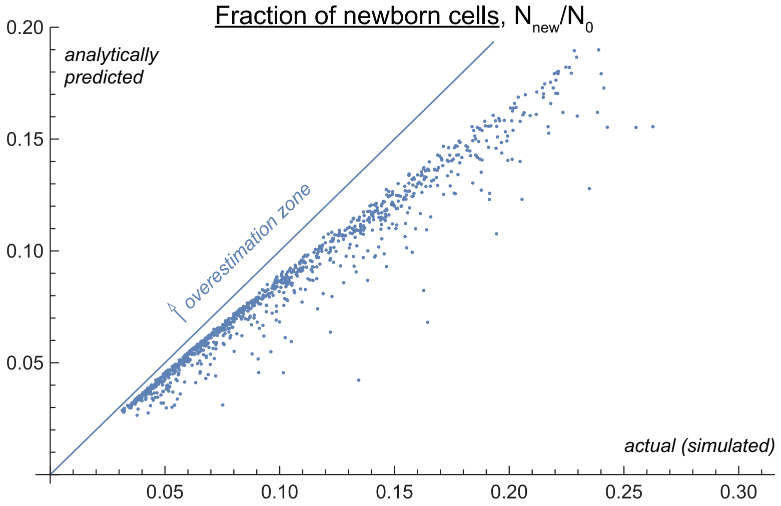


Figure S.14: Simulated versus predicted values of the number of newborn cancer cells during treatment by minimal single curative dose of pure radioconjugates.

#### S.3 Single-dose treatment by impure radioconjugates

The computational codes for this section are provided in supplementary file “03-Impure-drug.nb”.

##### S.3.1 Variation by radioconjugates impurity

Figure S.15 serves as an augmentation to Fig. 3A of main paper, showing the amount of toxic decays and the fraction of activity spent of viable cells during treatment by minimal single curative dose upon increase of the coefficient of drug impurity, $\eta$.

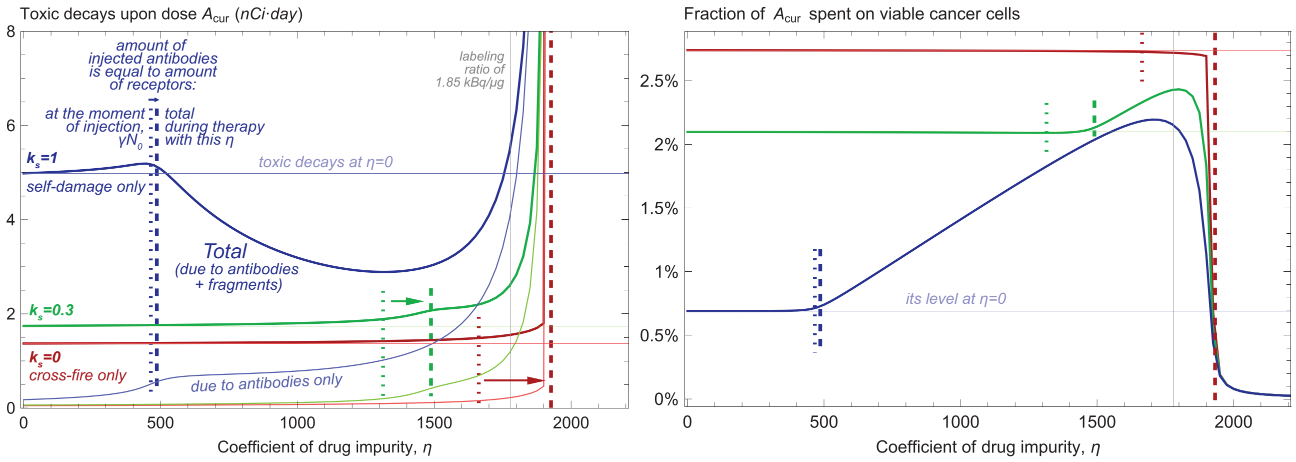


Figure S.15: Effect of the coefficient of drug impurity, $\eta$, on the toxicity of treatment by minimal single curative dose, $A_{cur}$, and on the fraction of it spent on viable cancer cells.

Figure S.16 illustrates the notion that in real-life conditions, due to the inherent uncertainly and variability of characteristics within a population, the use of the lowest possible coefficient of drug impurity represents a reasonable strategy for a single-dose treatment. The survival curves correspond to the single group of one thousand virtual mice, treated by 600 nCi of activity with different coefficient of drug impurity.

The limit of $A_{cur}$ at infinite drug impurity, shown in Fig. 3B of main paper, is estimated as follows. As $\eta \to \infty$, the number of radioconjugates attracted by cancer cells tends to zero, and the damage from unanchored

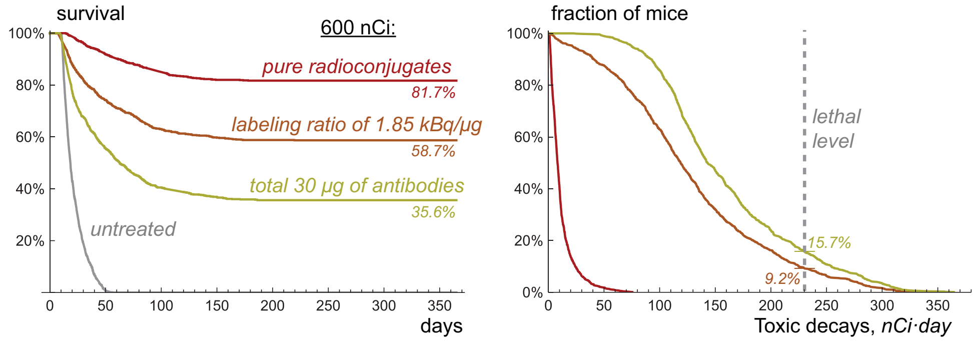


Figure S.16: Results of *in silico* trial for a single injection of radioconjugates with activity 600 nCi and coefficients of drug impurity: 0 (pure radioconjugates); 1780, (labeling ratio of 1.85 kBq/μg); 4444, (30 μg of daratumumab in total), for a single group of one thousand virtual mice. *Left*, survival curves. *Right*, toxicity curves.

nuclides eventually begins to play decisive role. In the limit, when it is the only contributor to the curative effect of treatment, the dynamics of viable cancer cells is governed by the following system:

$$\begin{aligned} &\text{Concentration of active antibodies in plasma:} && a' = \overbrace{-\lambda a}^{\text{decay}} \overbrace{-\kappa_c a}^{\text{clearance}}; \\ &\text{Number of viable cancer cells:} && N' = \overbrace{\rho N}^{\text{proliferation}} \overbrace{-k_f \alpha\lambda \cdot aN}^{\text{damage}}. \end{aligned} \tag{S.11}$$

This system is solved similarly to Eqs. (S.3). The equation for antibodies is independent, its solution is

$$a(t) = \frac{A}{V} e^{-[\kappa_c+\lambda]t},$$

where $A$ is the dose injected at $t = 0$. That leads to the analytical solution for the number of viable cancer cells:

$$N^{UN}(t) = N_0 \cdot exp(\rho t - k_f\alpha \cdot \frac{\lambda}{\lambda+\kappa_c} \cdot \frac{A}{V}[1 - e^{-[\lambda+\kappa_c]t}]),$$

superscript UN highlighting the validity of this formula for unanchored-nuclides-only case. When the derivative of this function is zero, the minimal number of viable cancer cells, $N_m^{UN} = N^{UN}(t_m^{UN})$, is achieved. That yields

$$t_m^{UN} = \frac{ln(\frac{A}{V} \cdot \frac{k_f\alpha\lambda}{\rho})}{\kappa_c+\lambda}, \; N_m^{UN} = N_0 \cdot [\frac{A}{V} \cdot \frac{k_f\alpha\lambda}{\rho}]^{\frac{\rho}{\kappa_c+\lambda}} \cdot exp([\rho - \frac{A}{V}k_f\alpha\lambda]/[\kappa_c+\lambda]).$$

The minimal curative dose that corresponds to $N_m^{UN} = N_{cur}$ is

$$A_{cur}^{UN} = -\frac{V\rho}{k_f\alpha\lambda} \cdot W_{-1}(-e^{-1}[\frac{N_{cur}}{N_0}]^{[\kappa_c+\lambda]/\rho}).$$

For the basic set of parameters, it is ≈37,300 nCi.

##### S.3.2 Parameter sweep for labeling ratio of 1.85 kBq/μg

As Fig. S.17 shows, analytical Eqs. (S.6)-(S.7), obtained for estimation of minimal single curative dose under assumption of pure radioconjugates, remain accurate as long as the amount of injected antibodies does not exceed cancer binding capacity at the moment of drug injection: $\xi \equiv [\eta+1]A_{cur}/[\gamma N_0] < 1$. Further overestimations correspond to the effect of decrease of curative dose due to the redistribution of radioconjugates towards viable cells, which is more pronounced when relative significance of self-damage, $k_s$, is close to one. Until $\xi \approx 2$ the formulas for $A_{cur}$ yield only moderate underestimations which can be corrected on statistical basis using the formulas introduced in Section S.2.2.1 (see supplementary code). For large $\xi$ the nuclides on cancer cells cannot provide cure, and corresponding estimations are inapplicable.

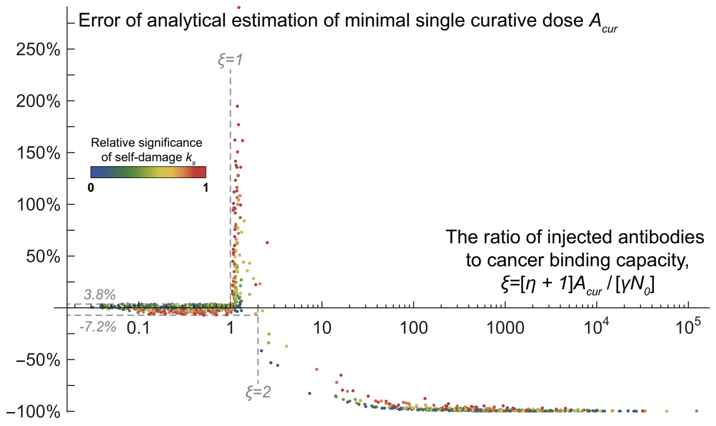


Figure S.17: Errors of analytical estimation of minimal single curative dose $A_{cur}$, Eqs. (S.6)-(S.7), in dependence of the ratio of injected antibodies to cancer binding capacity for a global parameter sweep with $\eta = 1780$.

The estimation of maximal safe dose, $A_{sf}$, is based on the following reasoning. In absence of cancer cells all the toxic decays would be due to the nuclides attached to intact antibodies, as they are not broken down to fragments by cancer cells. The overall level of toxic decays thus can be estimated from the relative rates of clearance of antibodies and decays of nuclides, from which it follows that

$$A_{bl}|_{\gamma N_0=0} = \frac{\lambda}{\lambda + \kappa_c} \cdot A.$$

Given $\lambda = 0.07$ and $\kappa_c^{min} = 0.04$, up to ≈64% of injected dose would contribute to toxicity for the considered parameter range. The following dose is the maximal dose that is guaranteed to be safe:

$$A_{sf}|_{\gamma N_0=0} = A_{bl}^{cr} \cdot \frac{\lambda + \kappa_c^{min}}{\lambda},$$

which is ≈362 nCi. As cancer capacity grows, an increasing amount of nuclides is retrieved from the bloodstream. They either decay on cancer cells or return to blood being attached to fragments of antibodies, which are rapidly cleared out. Each of these paths contributes to decrease of treatment toxicity. As long as cancer binding capacity is smaller than the amount of injected antibodies, in the worst-case scenario cancer attracts only $\gamma N_0/[\eta + 1]$ nuclides from blood, the remaining part staying on intact antibodies. The toxicity due to the backflush of radioactive fragments can be estimated via Eq. S.9, which always overestimates it, as it was shown in Section S.2.2.1. When cancer binding capacity exceeds the amount of injected antibodies, toxicity can be assessed via Eq. S.1, which should also overestimate it. Thus, the overall level of toxic decays cannot exceed

$$A_{bl}^{M} = \max\Big(\frac{\lambda}{\lambda + \kappa_c} \cdot [A - \frac{\gamma N_0}{\eta + 1}] + \frac{\omega \cdot \lambda}{[\lambda + \omega][\lambda + \kappa_p]} \cdot \frac{\gamma N_0}{\eta + 1}, \frac{\lambda}{\lambda + \kappa_c + k_{on}\gamma N_0/V} \cdot A + \frac{\omega \cdot \lambda}{[\lambda + \omega][\lambda + \kappa_p]} \cdot A\Big). \tag{S.12}$$

The worst scenario is realized under the slowest possible clearance rates $\kappa_c^{min} = 0.04$ and $\kappa_p^{min} = 0.4$ and highest possible death rate of damaged cancer cells $\omega^{max} = 0.5$. Under considered coefficient of drug impurity $\eta = 1780$ the second expression becomes greater than the first one only for $\gamma N_0 \approx 238$, which allows simplifying the formula and results in the following estimation of maximal safe dose:

$$A_{sf} = \min\Big(\frac{\lambda + \kappa_c^{min}}{\lambda} \cdot \Big\{A_{bl}^{cr} - \frac{\omega^{max} \cdot \lambda}{[\lambda + \omega^{max}][\lambda + \kappa_p^{min}]} \cdot \frac{\gamma N_0}{\eta + 1}\Big\} + \frac{\gamma N_0}{\eta + 1}, \frac{[\lambda + \omega^{max}][\lambda + \kappa_p^{min}]}{\omega^{max} \cdot \lambda} \cdot A_{bl}^{cr}\Big). \tag{S.13}$$

For sufficiently high cancer binding capacity, it is equal to the second expression, which is ≈1763 nCi for the used parameter values. This value is independent of the coefficient of drug impurity $\eta$. However, as Fig. S.18 shows, the use of lower labeling ratio would allow using greater doses safely under lower cancer binding capacity, while dilution of drug in unlabeled antibodies would have the opposite effect.

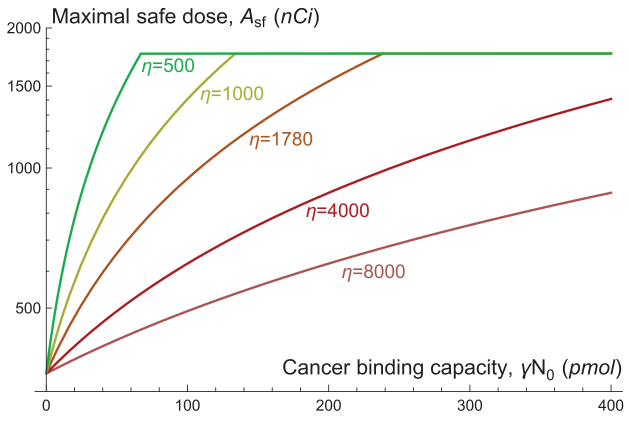


Figure S.18: Dependence of maximal safe dose $A_{sf}$ on cancer binding capacity $\gamma N_0$ under variation of coefficient of drug impurity $\eta$.

### S.4 Investigation of the model of heterogeneous cancer

Computational codes for this section are provided in supplementary file "04-Impure-drug-Hetero.nb".

#### S.4.1 Augmentation of the model, accounting for heterogeneity of cancer cells

As it was noted in Section S.1.3.5, our experimental data suggest notable intrapopulation heterogeneity of cancer cells. In particular, the experimental data are better reproduced with the use of notably different values of cancer cell radiosensitivity, which decreases with the increase of applied dose. It is reasonable to assume that such pattern is due to the inherent heterogeneity of radiosensitivity of cancer cells. More sensitive cells are the first ones to be killed, rendering the remaining cell population increasingly radioresistant along the course of therapy.

Another aspect that our experimental data suggests is that the cancer cells remaining after administration of high doses proliferate slower than the untreated population, even after the vast majority of radioactivity has already been deposited. This is in accordance with the generally accepted notion that radiosensitivity of cells correlates with their proliferation rate. Namely, for 300 nCi and 600 nCi of injected $^{225}$Ac-DOTA-daratumumab, the parts of the experimental data showing cancer regrowth are better fitted by exponential growth of cancer cells with $\rho = 0.28$ and 0.2, respectively, starting from day 35, by which less than 9% of activity should remain. This is about 82% and 59% of the estimated value of proliferation rate of untreated cancer cells. Therefore, we also introduce variable proliferation rate of cancer cells within the model, which is linked with their variable radiosensitivity. The following equations represent the model augmentation, aimed to capture heterogeneous response of cancer cells:

$$\begin{aligned} \text{Proliferation rate:} \quad & \rho \equiv \rho(t) = -k_D \cdot \rho \cdot ln(\frac{\rho}{\rho_m}) \cdot R_D(N, D, f_{AN}, f_{AD}, a, p); \\ \text{Radiosensitivity:} \quad & \alpha \equiv \alpha(t) = \frac{\rho - \rho_m}{\rho_0 - \rho_m} \cdot [\alpha_0 - \alpha_m] + \alpha_m. \end{aligned} \tag{S.14}$$

Four values of parameters were optimized by simultaneous fitting of simulation results to experimental data of all five mice groups, treated with different doses of $^{225}$Ac-DOTA-daratumumab. They are: minimum cancer cell proliferation rate, $\rho_m$; minimum cancer cell radiosensitivity, $\alpha_m$; initial (and maximum) cancer cell radiosensitivity, $\alpha_0$; and sensitivity of population-based parameters to radiation $k_D$. Their rounded up basic values are presented in Table S.2, where initial cancer cells proliferation rate, $\rho_0$ is transferred from the homogeneous model. The variation ranges are suggested with account of the fitting results of Section S.1.3.

Figure S.19 illustrates the simulations of the augmented model with basic values of parameters. Obviously, the quality of the fitting can be further improved by the introduction of additional processes in the model. The most prominent discrepancies are overestimation of treatment efficacy in 50 nCi setting and its underestimation in 100 nCi setting. This is linked with the apparently non-monotonic nature of decrease of cancer cell radiosensitivity in the experiments, which was illustrated above for homogeneous model in Fig. S.5. Its

Table S.2: Parameters for augmented model accounting for cell heterogeneity.

| | Parameter | Basic value | Range |
|---|---|---|---|
| $k_D$ | sensitivity of population-based parameters to radiation | 0.6 | 0.1-1.5 |
| $\rho_0$ | initial (and maximum) cancer cells proliferation rate | 0.34 | 0.15-0.7 |
| $\rho_m$ | minimum cancer cell proliferation rate | 0.2 | 0.05-min(0.25,$\rho_0$) |
| $\alpha_0$ | initial (and maximum) cancer cell radiosensitivity | 2400 | 500-4000 |
| $\alpha_m$ | minimum cancer cell radiosensitivity | 90 | 20-400 |

upper left plot suggests that the effective values of radiosensitivity are close to each other in these two settings, while for 25 nCi the value of optimal $\alpha$ is about twice greater. A possible explanation for this effect is the phenomenon of increased radiosensitivity of cells under low doses, which was demonstrated experimentally for a number of cell lines for external beam irradiation, with the use of doses lower than 1 Gy [9]. Although the cancer cells in our experiment are expected to receive higher total doses even for the lowest injected amount of drug, their irradiation in our setting is effectively prolonged for weeks. This leads to an even lower effective rate of irradiation compared to the external beam setting, in which the dose is delivered within minutes. However, to the best of our knowledge, no specific experiments have been performed to quantitatively study the effect of increased radiosensitivity under low doses for radiopharmaceutical therapy.

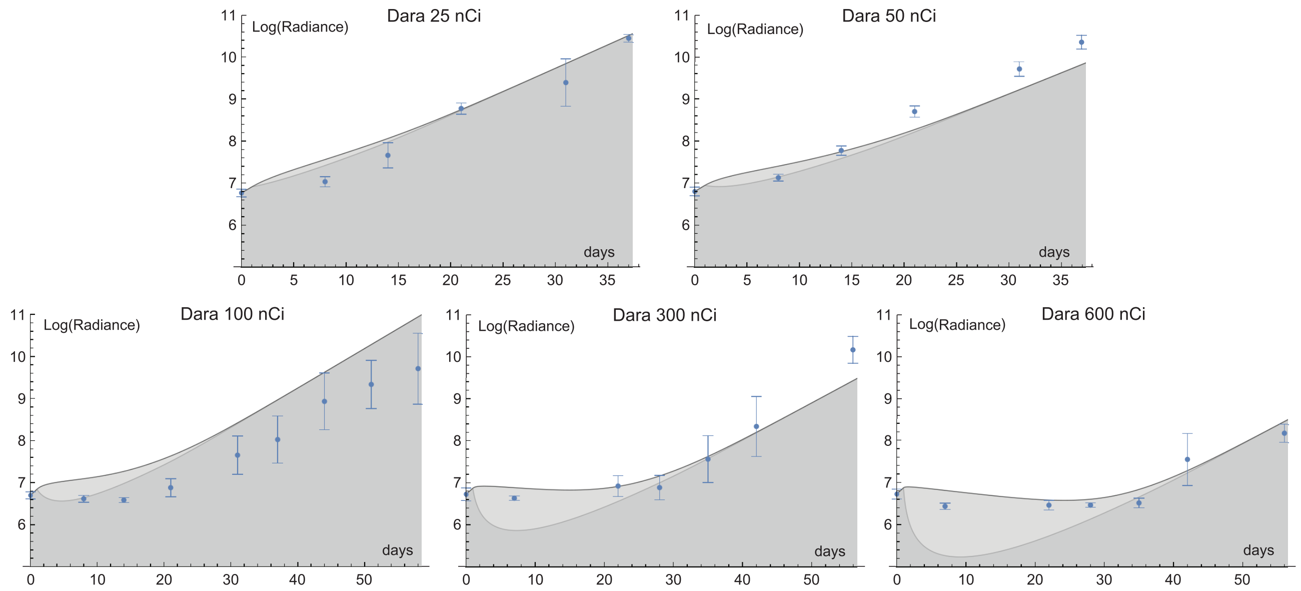


Figure S.19: Simulations with optimal values of parameters of augmented model, capturing cancer cell heterogeneity, along with the experimental data with $^{225}$Ac-DOTA-daratumumab.

Another discrepancy of simulated and experimental data is the overestimation of cancer cell numbers during the first days after drug injection. This could be attributed to various possible reasons, including strong manifestation of early forms of cell death in response to irradiation (e.g., apoptosis prior to mitosis [9]) and deceleration of luciferase production by damaged cells which would affect the observed radiance.

In this study we assign all the observed cancer cell heterogeneity to the variation of their inherent cell radiosensitivity and proliferation rate. It should be noted that the experimentally observed non-homogeneous response of cancer cells to irradiation can be due to other reasons as well, and more probably a combination of them underlies cancer cell response. In particular, it is reasonable to expect that cancer cells should have different number of specific receptors and thus attract different number of nuclides, yielding differential exposure to their decays. Moreover, under low injected doses, accompanied in our experimental setting with high coefficients of drug impurity, each cancer cell is expected to anchor few radionuclides, with some fraction of cells remaining devoid of them (as can be straightforwardly estimated from Poisson distribution). Furthermore, radiotherapy should affect the cellular composition of bone marrow and exacerbate its non-uniformity with time, with separated cancer cells being harder to eliminate [12]. However, the amount of experimental data at hand is insufficient for the introduction of new terms in the model that would describe additional processes in a physiologically reasonable way, improving the quality of fitting. The developed model already provides a good framework for a theoretical investigation that yields potentially valuable insights for clinical setting. Therefore, these fitting results are overall satisfactory for our purposes.

Simulations of 600 nCi setting suggest that in it about 39% of injected activity is spent in form of decays in blood. These decays are considered to be linked with the damage of the crucial organ at risk, bone marrow. Since this amount of radioactivity has led to one mouse death due to toxicity, we've taken take 230 nCi·day as the lethal level of decays in blood.

The introduced modifications worsen cancer susceptibility to treatment, compared to homogeneous model. The basic set of parameters, provided in Table S.2, corresponds to minimal single curative dose of 220 nCi for the case of pure radioconjugates. While the corresponding amount of them still can occupy less than one percent of cancer receptors, the use of labeling ratio of 1.85 kBq/μg prevents their fitting onto the cancer cells and escalates the minimal single curative dose by a factor of ≈500, rendering the treatment lethally toxic.

##### S.4.2 Global parameter sweep with pure radioconjugates

The global parameter sweep for heterogeneous cancer model was performed similarly to the sweep in Section S.2.2 in order to determine how this augmentation alters model behavior. One thousand sets of values of thirteen parameters – $\kappa_c$, $\kappa_p$, $\gamma$, $V$, $k_s$, $\rho_0$, $\rho_m$, $\omega$, $\alpha_0$, $\alpha_m$, $k_f$, $N_0$, $k_D$ – were selected randomly within the ranges designated in Table 1 of main paper and Table S.2, with $\eta$ kept to 0.

Like in the case of homogeneous model, the majority of cancer cell receptors remained free during each treatment, with the maximum receptors occupancy not exceeding 6.3%. Only one case was lethally toxic due to the backflush of radionuclides to the bloodstream. Both parameter sweeps for homogeneous and heterogeneous cancer model yielded similar distributions for the fractions of activity released in blood, with Eqs. (S.8) and (S.9) maintaining good accuracy for the estimation of the level of toxic decays in heterogeneous model (see supplementary code).

The value of minimal single curative dose, $A_{cur}$, for heterogeneous cancer model in the majority of simulated cases can be reasonably estimated with the use of modified Eqs. (S.6)-(S.7), derived for homogeneous cancer, as illustrated in Fig. S.20. Parameters $A^0_{est}$ and $A^m_{est}$ refer to the estimated values of doses, obtained correspondingly for the most radiosensitive cell population, characterized by parameters $\rho_0$ and $\alpha_0$, and the most radioresistant population, characterized by $\rho_m$ and $\alpha_m$. As the left plot shows, in most cases the actual value of $A_{cur}$ is largely determined by the latter population, being close to $A^m_{est}$. This fact has a simple intuitive explanation, since in heterogeneous cancer setting the curative density of nuclides on cancer cells should be sufficient to eliminate each of them, and therefore is overall determined by the most radioresistant cells.

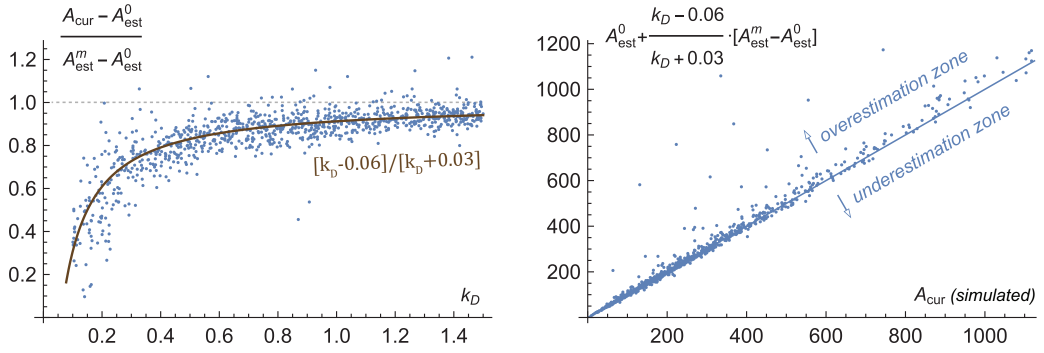


Figure S.20: Method of estimation of minimal single curative dose $A_{cur}$ for heterogeneous cancer model.

This implies that during treatment by minimal single curative dose the radiosensitive cancer cells should receive greater amount of radiation energy, than is sufficient for their elimination. Due to that, in heterogeneous cancer model the fraction of injected activity spent on viable cancer cells is generally lower than in homogeneous model, as the left plot in Fig. S.21 illustrates. Furthermore, the accompanying fast killing of radiosensitive cancer cells leads to strong inhibition of cancer cell proliferation in the beginning of treatment. This yields overall lower amounts of newborn cancer cells during treatment in heterogeneous model, as right plot in Fig. S.21 shows.

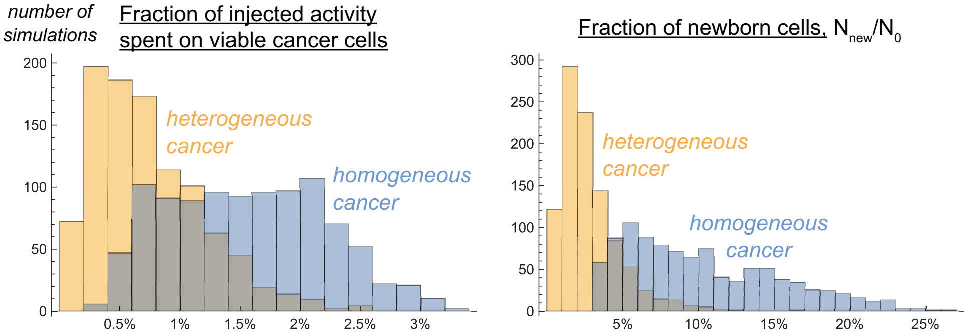


Figure S.21: Comparison of the fractions of injected activity spent on viable cancer cells and the fractions of newborn cells during treatment for homogeneous and heterogeneous cancer models.

##### S.4.3 Parameter sweep for labeling ratio of 1.85 kBq/µg

Figure S.22 shows the results of global parameter sweep of heterogeneous cancer model with the use of coefficient of drug impurity $\eta = 1780$. It is similar to Fig. 4 of main paper, corresponding to homogeneous model, and confirms the previously obtained qualitative result for it.

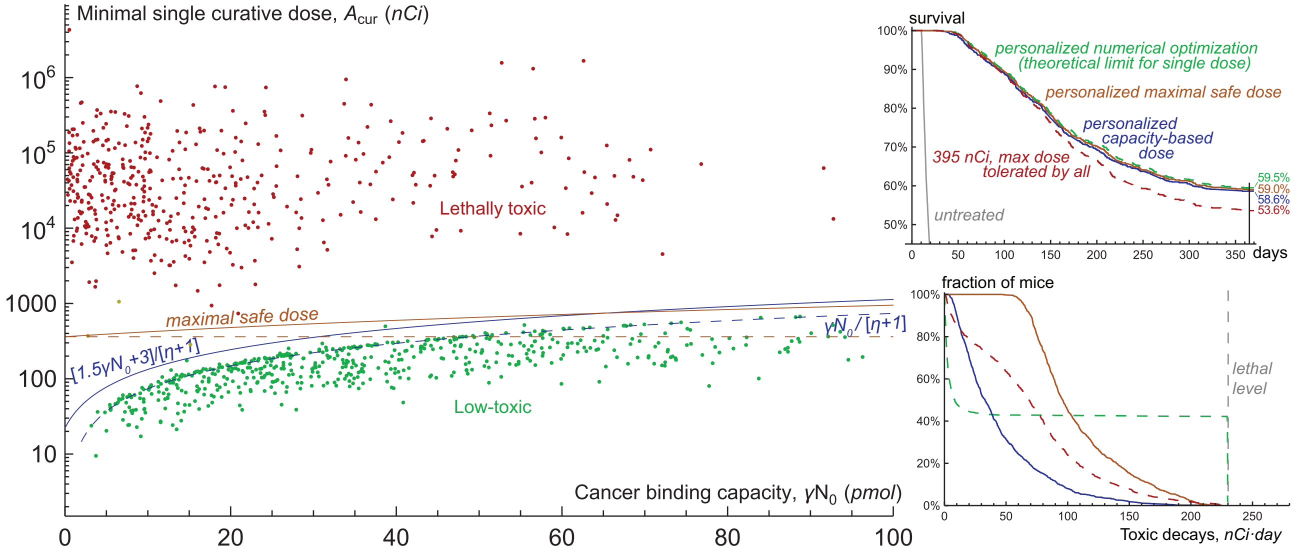


Figure S.22: *Left*, scatter plot of cancer binding capacity, i.e., total amount of specific receptors on cancer cells at the moment of drug injection, and minimal single curative dose produced by the global parameter sweep in heterogeneous cancer model within the training set of one thousand virtual mice. Coefficient of drug impurity, $\eta$, corresponds to the labeling ratio of 1.85 kBq/µg. *Right*, survival curves and toxicity curves for a single test set of one thousand virtual mice, treated by different approaches.

One notable difference in the results for two models is that the fraction of newborn cells and therefore the increase of cancer binding capacity during single-dose curative treatment is generally more moderate in heterogeneous cancer model. Therefore, the low-toxic cases in the left plot tend to concentrate closer to the blue dashed curve for which the amount of injected antibodies, $[\eta + 1]A_{cur}$, is equal to cancer binding capacity $\gamma N_0$ at the moment of drug injection.

#### S.5 Optimization of multi-dose treatment

We pose the task of optimization of multi-dose treatment as the problem of finding the schedule of injections that would cure a virtual mouse yielding the lowest possible amount of toxic decays in blood, if cure is feasible, otherwise would prolong the mouse survival as long as possible. For that, we use the homogeneous cancer model, presented in main paper. As previously, we assume that a mouse dies either at the moment when the number of cancer cells reaches $C_d = 10^{11}$ or the amount of toxic decays in its blood reaches $A_{bl}^{cr} = 230$ nCi·day. For the sake of computational convenience, we use no more than ten injections and allow administering them at equal intervals at the moments of time, corresponding to integer numbers of days, i.e., $t_i = 0, 1, 2, \ldots$. Also, we limit the maximum duration of simulations by 365 days for non-cured mice. Thus, a mouse can be alive at this moment if either it is cured, or the schedule of drug injections allows suppressing cancer growth up to this moment without reaching lethal toxicity level. We keep the coefficient of drug impurity $\eta = 1780$, and we use the training and test parameter sets containing one thousand virtual mice each, which have already been generated within single-dose treatment study.

The optimization algorithm in described in Section S.5.2.2. Within it, for curative cases we rely on three solutions of different optimization tasks. The first one is finding the optimal single-dose schedule, this task having already been performed. The second task is finding the optimal two-dose schedule, the corresponding algorithm is described in Section S.5.1. The third task is finding the optimal multi-dose schedule without predefined number of doses, which solution relies on additional simulations of the model under certain approximations, described further. A separate solution for the schedules containing two doses was used since this restriction assures the proximity of solution to the global optimum, which is hard to guarantee in a general multidimensional optimization task. Therefore, in some cases such two-dose schedules indeed represent a more optimal solution than the one found without restriction on the number of doses.

The computational codes for this section are provided in supplementary file "05-Multi-dose.nb".

##### S.5.1 Optimization of two-dose treatment

###### S.5.1.1 Algorithm for optimization of two-dose treatment

The search for two-dose schedules, leading to cancer cure with the lowest possible amount of toxic decays in blood, was performed via the following algorithm based on classical gradient descent. It is thus implicitly assumed that the optimization problem is convex, which was suggested empirically based on model behavior.

The search for the optimal schedule takes place in a two-dimensional parameter space. The first parameter represents the ratio of the first dose in the two-dose schedule and the minimal single curative dose. Its admissible range was set as [0.05, 0.95]. The first dose is always applied at $t = 0$ since the previous results suggest that delaying the beginning of treatment can lead only to the increase of the amount of drug needed for cancer cure. The second parameter is the interval between two doses, which cannot be less than 1 day. These two parameters identify a two-dose schedule in an unambiguous manner, since its second dose represents the uniquely defined minimal dose, leading to cancer cure when injected at the specific moment of time. Thus defined *current minimal curative dose* can be viewed as a latent variable, which evolves with ongoing system dynamics.

The work of the algorithm of two-dose optimization for the basic set of parameters is illustrated in Fig. S.23. Briefly, it involves a series of steps within this parameter space, each of which is performed if it leads to a decrease of the amount of accompanying toxic decays. In the beginning of algorithm operation relatively big steps are taken, and their size decreases when no steps can be taken in any of the eight possible directions in terms of the used parametric plane, which are up, down, left, right and four diagonals. The minimal size of the step was set as 1% on the dose-ratio axis and 1 day on the interval axis, or the same simultaneous changes in any of the diagonal directions. For the optimizations within the parameter sweep the search began at initial point (0.8, $2t_m$), where $t_m$ is the moment at which viable cancer cell number reaches its minimum for a single-dose schedule with minimal single curative dose, rounded up to an integer number of days.

###### S.5.1.2 Optimized two-dose schedules under variation of particle range

Figure S.24 illustrates the model simulations with two-dose optimal schedules for the basic set of parameters and varied significance of self-damage, $k_s$. These simulations highlight the general factors that make optimization

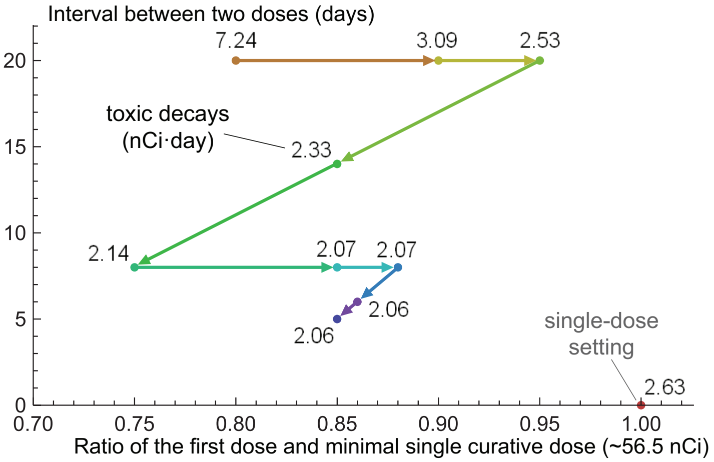


Figure S.23: Illustration of work of the algorithm of two-dose treatment optimization for basic parameter set.

of targeted radionuclide therapy possible. Three factors of that kind can be distinguished, which relative importance depends on the value of $k_s$.

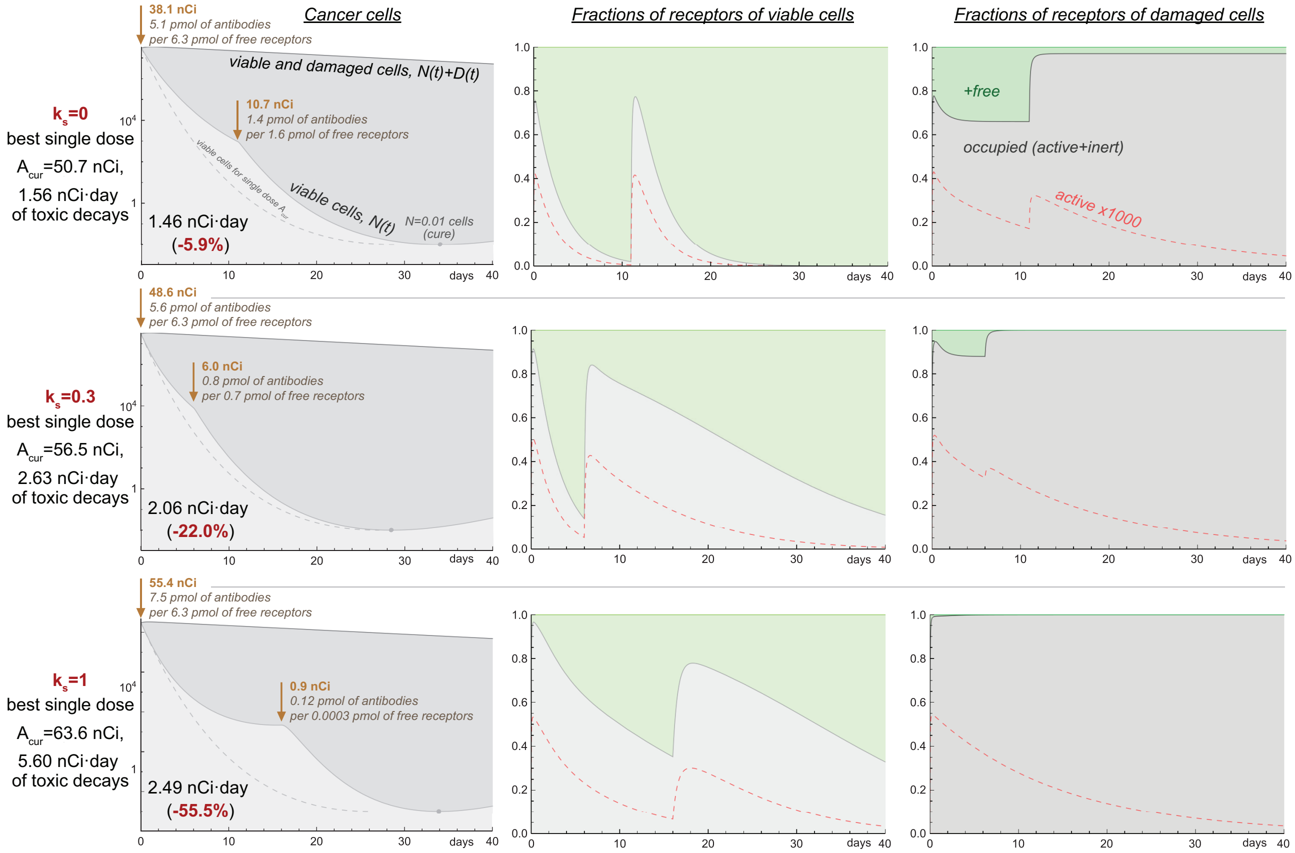


Figure S.24: Simulations of treatments by optimal two-dose schedules for the basic set of parameters and relative significance of self-damage $k_s = 0$ (top row); 0.3 (middle row) and 1 (bottom row).

*Inability of minimal single curative dose to fit on cancer cells.* In each of the considered simulations the amount of antibodies to be injected with the corresponding minimal single curative dose, $[\eta + 1]A_{cur}$, is greater than the initial cancer binding capacity, $\gamma N_0$. During a single-dose treatment that leads to the fact that initially a notable fraction of injected antibodies remains in the bloodstream until they are cleared out or the newborn cancer cells attract them. At that period the decays of nuclides, attached to the radioconjugates floating in the bloodstream, significantly contribute to treatment toxicity. In contrast, the nuclides that are attached to cancer cells, are eventually less toxic, since only a fraction of them returns in the bloodstream attached to antibody fragments, that are furthermore cleared out faster than intact antibodies.

This factor suggests avoiding oversaturation of cancer binding capacity in order to decrease toxicity. This method is efficient in the cross-fire-only setting, the optimal schedule outcome for which is illustrated in the upper plots. In this schedule the amounts of antibodies in both doses are lower than the total amounts of free receptors on cancer cells at the moments of their injection.

*Decrease of total cancer cell number during treatment.* Each of the first doses in the optimized schedules represents only a fraction of a corresponding minimal single curative dose, and therefore provides slower initial decrease of the number of viable cancer cells, as confirmed by the left three plots. Nevertheless, the first doses in the optimized schedules are sufficiently high to yield the continuous decrease of the total number of cancer cells, due to the death of damaged ones. The second injected doses are thus distributed over fewer cancer cells. This fact is of particular importance for cross-fire, which efficiency is proportional to the average density of nuclides on all cancer cells, both viable and damaged.

Therefore, the decrease of cancel cell number with time justifies delaying injection of the second dose, despite the counteracting ongoing repopulation of viable cancer cells and despite the decrease of activity remaining on their receptors.

*Blockage of receptors on damaged cancer cells.* When cancer cells are nearly saturated by antibodies, the radioconjugates remaining in the bloodstream are predominantly attracted by new free receptors which are produced only by viable cancer cells. In the case of significant self-damage this contributes to the redistribution of dose towards viable cells.

This represents an effective method for increasing efficacy of treatment using short-range particles. The lower row in Fig. S.24 corresponds to the self-damage-only case, for which the optimal two-dose schedule yields over twice less toxic decays than the optimal single-dose schedule. In contrast with cross-fire-only case, in self-damage-only case the amounts of injected antibodies in both doses exceed the current amounts of free receptors, yielding rapid blockage of receptors on damaged cells and ensuring prolonged binding of radioconjugates to the newly expressed receptors on viable cells, as witnessed by the lower right and middle plots.

These results show that the range of emitted particles guides qualitatively different methods towards optimization of multi-dose treatments, with self-damage-only setting having the greatest potential for optimization. The optimal schedule for the intermediate value of $k_s = 0.3$, illustrated in the middle plots, provides a compromise achieving maximal benefit from the indicated factors.

#### S.5.2 Optimization of multi-dose treatment without predefined number of doses

The current results suggest that within the studied ranges of model parameters the curative effect of treatment is provided mainly by the nuclides attached to cancer cells, while the nuclides floating in the bloodstream have negligible efficacy to toxicity ratio. This suggests that the extreme setting when cancer cells are fully occupied by antibodies represents an important special case. If cancer cannot be cured in this approximated setting, then in the original model achieving cancer cure without lethal toxicity is highly unlikely. This setting will serve as the foundation for the algorithm of optimization of multi-dose treatment without predefined number of injections. Its analytical study is performed below.

##### S.5.2.1 Analytical estimation of cancer curability under neglect of irradiation from unanchored nuclides

Let's assume that a virtual mouse has already obtained the first injection of antibodies that saturated cancer cells but cannot result in cancer cure. Consider an idealized approximation in which from now on all the receptors, including newly produced ones, are immediately bound to antibodies. Also let's assume that the damage from unanchored nuclides is negligible, $k_f = 0$.

Note that the assumption of absence of free receptors, $f_{FN} \to 0$, $f_{FD} \to 0$, formally implies that the level of radioconjugates in blood tends to infinity, guaranteeing the corresponding treatment to be lethally toxic. Thus, this approximation is valuable only as a limiting case that allows identifying the cancers which can potentially be cured by multi-dose treatment. However, by itself it does not yield insights on how to optimally achieve cure and whether cure can be achieved without exceeding lethal toxicity level.

We assume that initially the active and inert antibodies bind to receptors in proportion of $1 : \eta$, defined by the used coefficient of drug impurity. However, in the limit of negligible number of free receptors this ratio will decrease in the long term, since a fraction of nuclides will inevitably decay before anchoring to cancer cells. The effective long-term ratio of inert to active antibodies binding to cancer cells $\eta^*$ can be estimated as the ratio of stationary solutions of the following system:

$$a' = J - \lambda a - \kappa_c a,$$
$$b' = \eta J + \lambda a - \kappa_c b,$$

which is an approximation of antibodies dynamics under constant rate of their infusion and neglect of their binding. It can be deduced straightforwardly that

$$\frac{b_C}{a_C} = \eta[1 + \frac{\lambda}{\kappa_c}] + \frac{\lambda}{\kappa_c}.$$

For the considered ranges of parameter values, the second term can be neglected and therefore

$$\eta^* \approx \eta[1 + \frac{\lambda}{\kappa_c}],$$

which for the used parameter ranges implies the increase of effective coefficient of drug impurity in long term by 1.25-2.75 times.

In this limiting case, the investigated model effectively boils down to the following system. Note that the terms of radiation damage are regrouped for convenience. Rather than corresponding to self-damage and cross-fire, as in main model equations, here they correspond to the damage of viable cells due to the nuclides attached to themselves and due to the nuclides located on damaged cells. As earlier, $\chi = \alpha\lambda\gamma/\nu$ for brevity.

$$\begin{aligned} N' &= \rho N - R_D \cdot N; \\ D' &= R_D \cdot N - \omega D; \\ f'_{AN} &= \frac{\rho}{\eta^* + 1} - [\lambda + \rho] f_{AN}; \\ f'_{AD} &= [f_{AN} - f_{AD}] \cdot R_D \cdot \frac{N}{D} - \lambda f_{AD}; \\ R_D \equiv R_D(N, D, f_{AN}, f_{AD}) &= \chi \cdot f_{AN} \cdot \frac{N + k_s D}{N + D} + \chi[1 - k_s] \cdot f_{AD} \cdot \frac{D}{N + D}. \end{aligned} \tag{S.15}$$

The fact whether cancer is curable is inferred by the sign of $\rho - R_D$ as $t \to \infty$. If the proliferation rate of cancer cells prevails over the rate of their damage, cancer eventually escapes eradication.

The equation for $f_{AN}$ is independent of other variables. Therefore, for the consideration of long-term behavior of this system we can replace this variable with its stationary state:

$$f^C_{AN} = \frac{1}{\eta^* + 1} \cdot \frac{\rho}{\lambda + \rho}. \tag{S.16}$$

Gradually, $f_{AN}$ should decrease tending to this value, as initially it is equal to $f_{AN}(0) = 1/[\eta + 1] > f^C_{AN}$.

At first, consider the situation in which the first injection has already damaged a significant number of viable cancer cells, so that $N << D$, and this condition is kept for sufficiently long period. In such case, the first term in equation for $f_{AD}$, i.e., the term of appearance of active receptors on damaged cells, can be neglected. Therefore, $f_{AD}$ will tend to zero in the long term, and the rate of radiation damage would fall down to $R_D = \chi k_s \cdot f^C_{AN}$. This expression implies that cross-fire is effectively canceled out and only self-damage of cancer cells takes place. Despite the efficacy of irradiation being now severely limited, the number of viable cells will decrease if $\chi k_s \cdot f^C_{AN} > \rho$. Cancer is thus potentially curable via self-damage only, if

$$C_{SD} \equiv k_s \cdot \frac{\alpha\lambda\gamma}{\nu[\eta^* + 1]} - [\lambda + \rho] > 0.$$

If that inequality is not met, then the number of viable cells grows. It can be deduced straightforwardly from Eqs. (S.15) that

$$[\frac{N}{D}]' = [\rho + \omega - R_D]\frac{N}{D} - R_D[\frac{N}{D}]^2.$$

Under current assumptions $N/D$ is yet negligible and $\rho > R_D$, which means that $N/D$ will grow until

$$\frac{N}{D} = \frac{\rho + \omega - R_D}{R_D}. \tag{S.17}$$

However, $R_D$ depends on $N/D$ by itself. Using for convenience $y = N/D$, we obtain from Eqs. (S.15):

$$f'_{AD} = f^C_{AN} R_D y - [\lambda + R_D y] f_{AD},$$

$$R_D = \chi\frac{y + k_s}{y + 1} f^C_{AN} + \chi\frac{1 - k_s}{y + 1} f_{AD}.$$

From the first equation the stationary state of $f_{AD}$ can be found as

$$f^C_{AD} = \frac{R_D y_C}{\lambda + R_D y_C} \cdot f^C_{AN},$$

where $y_C$ is the stationary state of $N/D$. Note that $f^C_{AD}$ cannot exceed $f^C_{AN}$. From this equation it follows that the stationary state of radiation damage function is defined by equation

$$R^C_D = [\{y_C + k_s\} + \{1 - k_s\}\frac{R^C_D y_C}{\lambda + R^C_D y_C}] \cdot \frac{\chi f^C_{AN}}{y_C + 1}.$$

On the other hand, from Eq. (S.17) it follows that

$$R^C_D = \frac{\rho + \omega}{1 + y_C}.$$

The last two equations result in the quadratic equation for $y_C$:

$$[\rho + \omega + \lambda]y_C^2 + \Big[\rho + \omega + \lambda - \frac{[\rho + \omega]^2}{\chi f^C_{AN}} + \lambda k_s - \frac{\rho + \omega}{\chi f^C_{AN}}\lambda\Big] y_C + \lambda k_s - \frac{\rho + \omega}{\chi f^C_{AN}}\lambda = 0,$$

with its straightforward but quite cumbersome to be written out explicitly positive solution having physical meaning of the stationary ratio of viable and damaged cells. It depends on the effective coefficient of drug impurity $\eta^*$, since $f^C_{AN}$ is a function of it, as indicated in Eqs. (S.16). The existence of a positive real solution is guaranteed by our initial assumption $C_{SD} < 0$, or $\chi f^C_{AN} < \rho/k_s$. Under this condition the free term of this equation is less than $-\lambda k_s \omega/\rho$ and thus is always negative. With the quadratic coefficient being always positive, it implies that this equation always has two real solutions of different signs.

The resulting condition for cancer curability in the long term follows from the requirement $R^C_D > \rho$:

$$C_{LT} \equiv \frac{\omega}{\rho} - y_C > 0.$$

The resulting formula is quite complicated to be straightforwardly interpreted. It is clear, however, that $C_{LT}$ should grow, firstly, with increase of the amount of receptors on cancer cells, $\gamma$, and their radiosensitivity, $\alpha$, which act as multipliers within the parameter $\chi$ and govern the rate of damage of cancer cells, see Eqs. (S.15). Secondly, it should grow with the increase of cancer cells death rate, $\omega$, as it promotes greater viable to damaged cells ratio and leads to faster appearance of active receptors on damaged cells. Thirdly, it should grow with the increase of significance of self-damage, $k_s$, as this demands smaller income from cross-fire irradiation for achieving cure.

Figure S.25 confirms this reasoning showing the regions in parameter space, corresponding to $C_{LT} > 0$, where cancers are estimated as curable in long term. This region always contains the region corresponding to curability by self-damage only, $C_{SD} > 0$, and at $k_s = 1$ they would coincide. These regions are also compared with the region approximately corresponding to cancer curability in single-dose setting:

$$C_0 \equiv \gamma N_0 - [\eta + 1] A_{cur}|_{\eta=0} > 0,$$

where $A_{cur}|_{\eta=0}$ is the estimation of minimal single curative dose of pure radioconjugates by Eqs. (S.6)-(S.7). This condition demands that this dose can fit onto the initial pool of cancer receptors at the moment of drug injection. As discussed above, the drug impurity affects the minimal single curative dose only slightly until it exceeds cancer binding capacity, justifying this estimation. However, moderate exceeding of cancer binding capacity would still maintain the curability of treatment due to the prolonged binding of radioconjugates to newly expressed receptors. Therefore, this condition underestimates the actual parameter space region, corresponding to cancers curable by a single dose.

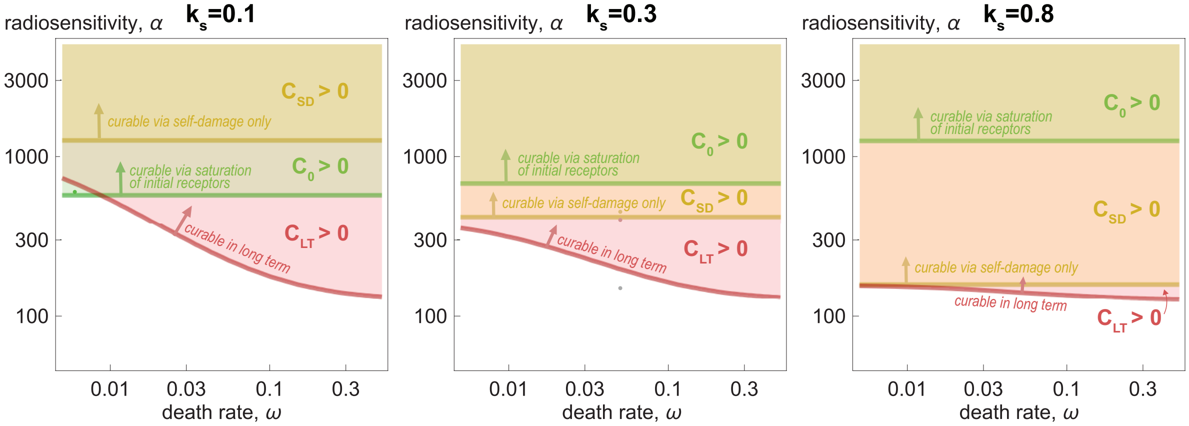


Figure S.25: Regions of potentially curable cancers in parameter space, under variation of $\omega$, $\alpha$ and $k_s$. The values of other parameters are from basic set. Dots marks the cases illustrated in Figs. S.26 and S.27.

The region corresponding to $C_0 > 0$ shrinks with the increase of self-damage significance, $k_s$. For sufficiently high values of $k_s$ it is contained within the regions corresponding to long-term curability. This implies that the use of multiple doses increases the range of curable cancers. For low values of $k_s$ the region with $C_0 > 0$ contains the region with $C_{SD}$ and intersects with the region with $C_{LT} > 0$. This implies that some cancers, that can be cured by a single dose, are incurable in the long term.

Figure S.26 illustrates the validity of analytical results. In the original model cancer can be cured under basic set of parameters by a single dose without achieving lethal toxicity. Basic value of cancer cell radiosensitivity is $\alpha = 500$. Upon its decrease to $\alpha = 450$ the amount of toxic decays in blood accompanying treatment by minimal single curative dose reaches 564 nCi·day, which exceeds lethal level. In the limit of immediate binding of all receptors cancer nevertheless can be cured within a month due to the sufficient efficiency of self-damage.

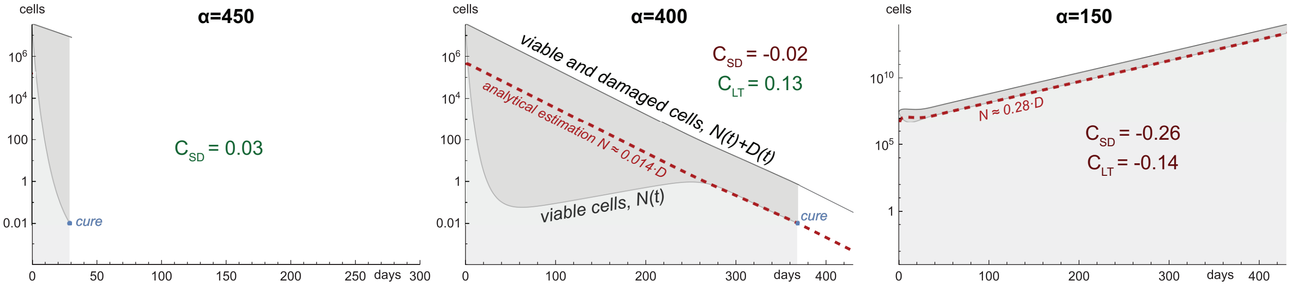


Figure S.26: Simulations of Eqs. (S.15), corresponding to the limiting case of immediate binding of all receptors to antibodies under basic set of parameters and variation of cancer cell radiosensitivity $\alpha$.

Upon further decrease of radiosensitivity down to $\alpha = 400$ self-damage irradiation by itself becomes insufficient to cure cancer, but due to the shrinkage of cancer mass the increasing rate of overall radiation damage eventually becomes high enough to guarantee long-term eradication of cancer cells. At $\alpha = 150$ cancer is incurable, and it continues to grow under full saturation of its receptors, with the ratio of viable to damaged cells corresponding to the analytically estimated value.

Overall, the criteria $C_{SD} > 0$ and $C_{LT} > 0$ suggest the possibility of cancer cure in the original model via a multi-dose schedule, however, it has yet to be checked for each parameter set whether cure can be achieved without exceeding the lethal level of toxic decays in blood. Additionally, as shown above, some cancers can be cured by a single dose, but they remain effectively incurable in the long term. The dynamics of a corresponding case is illustrated in Fig. S.27. Using physical analogies, it may be noted that the response of cancer to the initial dose resembles a pendulum swing which may or may not cross the curative threshold. In contrast, the long-term drug infusion is similar to a push with a constant force which may prove insufficient for cure.

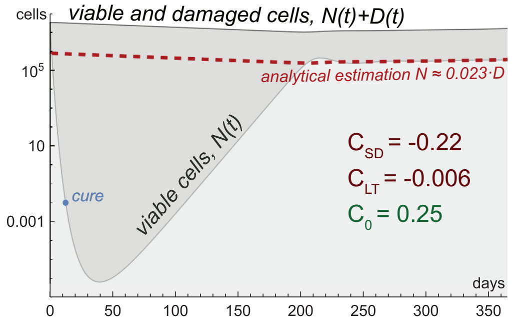


Figure S.27: Simulations of Eqs. (S.15) for $k_s = 0.1$, $\omega = 0.006$, $\alpha = 600$ and other parameters from basic set.

##### S.5.2.2 Resulting algorithm of multi-dose treatment optimization

The high-level scheme of the overall algorithm of multi-dose treatment optimization is presented in Fig. S.28.

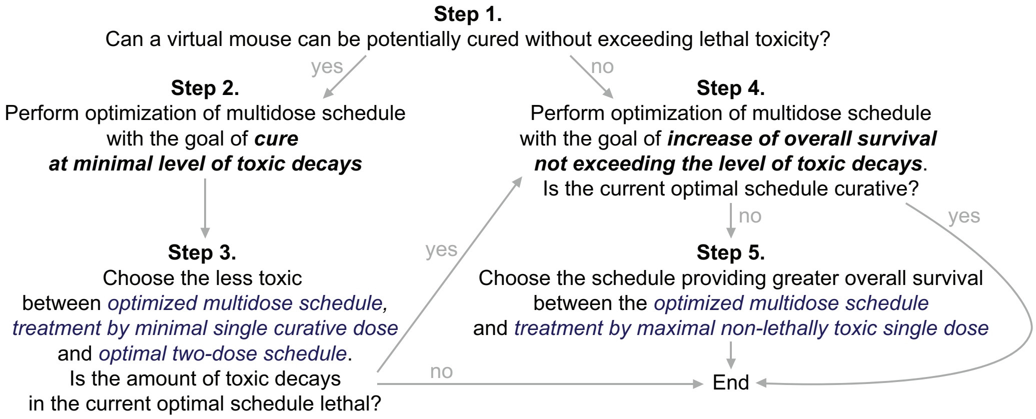


Figure S.28: High-level scheme of the algorithm of multi-dose treatment optimization.

*Step 1.* For each given parameter set, it is initially determined whether it can be reasonably expected that cancer can be cured without exceed of lethal toxicity. The virtual mice, that can be cured by minimal single curative dose with the corresponding amount of toxic decays not exceeding lethal threshold, already fit this criterion. Additionally, those parameter sets are selected, for which the cure is achieved within a year in the approximated model, corresponding to immediate binding of all receptors by antibodies. This model, governed by Eqs. (S.15), was analyzed in the previous subsection. In order to not discard potentially curative cases, we use the value of coefficient of drug impurity $\eta^* = \eta$ during this process.

*Step 2.* For those virtual mice that can potentially be cured, we perform multi-dose optimization without predetermined number of injections but keeping their maximum number equal to ten. The optimization goal is to achieve cure at minimal level of toxic decays.

For initial selection of potentially effective multi-dose schedules to be optimized further, we perform auxiliary simulations of cancer dynamics under varied constant concentration of active antibodies in blood, i.e., setting $a'(t) = 0$ in main model equations. Initial level $a = a_0$ for the variation is selected based on the simulation of Eqs. (S.15), which has already been performed in the previous step. Namely, using this simulation, we estimate the amount of antibodies that should have been injected to approximately correspond to its results. For that, we integrate the corresponding terms of nuclide decays on active receptors and nuclide release in the bloodstream, from initial moment until the achievement of cure $t_{cur}$, and we add the remaining amount of nuclides on cancer cells at that moment. By dividing the result by $V$ and $t_{cur}$, we obtain the estimated value of $a_0$. Starting with it and using bisection method, we find the constant level of active antibodies in blood $a_c$, which provides cure with minimal accompanying toxicity.

The corresponding solution serves as the basis for creating the multi-dose schedules to be tested in the original model. They are obtained by discretization of the continuous infusion term, which keeps concentration of active antibodies in blood at the level $a_c$, over variable time intervals $T$. Initial interval $T_0$ is chosen as $t_{cur}/10$, rounded up to days, ten being the maximal number of injections in considered schedules. Then $T$ is increased as long as the resulting schedule provides decrease of treatment toxicity, or $T$ becomes equal to $t_{cur}/2$, yielding two-dose schedule. Each multi-dose schedule is normalized by applying a common multiplier to all the doses so that the minimal number of viable cancer cells $N_m$ becomes close to $N_{cur}$; by removing last doses, if cure can be achieved without them at lesser toxicity; and finally by replacing the last dose with the current minimal curative dose, so that overall $N_m$ is kept equal to the curative threshold $N_{cur}$. In the resulting optimized multi-dose schedule the first dose is further varied, with applying similar methods of schedule normalization.

It should be noted that this technique overall yields suboptimal solutions, which is a common situation for multidimensional optimization problems. Additional alterations and improvements can be suggested, including introduction of non-equal intervals between doses and implementation of gradient descent with the constraint $N_m = N_{cur}$ and with the level of toxic decays regarded as the function of the array of injections, similarly to how it was performed in the work [13] for optimization of external beam therapy. However, such modifications would come at additional computational cost, and are expected to generally yield minor improvements. From the point of view of the current study, it is more important to find common patterns in the optimized schedules under variation of parameters, so that they can serve as a foundation for further suggestions of treatment optimization under significant uncertainty of model parameters, which is discussed in the next section. The algorithm used herein is thus sufficient for our goals.

*Step 3.* The resulting multi-dose schedule is compared to the optimal two-dose schedule, found by the algorithm, described in Section S.5.1, and to the schedule with minimal single curative dose, found by straightforward bisection method. In contrast with the algorithm implemented at the previous step, these methods yield the solutions which are close to the global optimums in single-dose and two-dose settings. The least toxic of these three schedules is selected at this step. If the corresponding amount of toxic decays is less than the lethal level $A_{bl}^{cr} = 230$ nCi·day, the algorithm terminates at this step for the consider set of parameters.

*Step 4.* If cancer is not defined as potentially curative at step 1, or if the optimized curative schedule produced by steps 2-3 is lethally toxic, we perform multi-dose optimization with the goal of prolonging the overall survival of a virtual mouse without achieving lethal toxicity. A virtual mouse is considered dead at the moment when the number of cancer cells reaches $C_d = 10^{11}$.

The solution of this problem is as well based on consideration of auxiliary version of the model with constant concentration of active antibodies in blood. Using bisection method, we find their constant concentration, providing greater value of overall survival, for simplicity estimated under assumption that the nuclides are eliminated from the system at the moment when the total amount of toxic decays reaches lethal level.

The testing of multi-dose schedules is performed similarly to the method described in step 2. The schedules are obtained by discretization of the continuous infusion term in the optimal solution of the auxiliary model, starting with the interval between injections $T = 1$ day. The interval is increased as long as the resulting schedule provides greater overall survival. Each schedule is normalized by applying a common multiplier for all doses so that the amount of toxic decays is close to $A_{bl}^{cr}$, since this action yields maximal overall survival without exceeding this threshold. If during these operations overall survival exceeds 365 days, which is chosen as the maximum observation time, this step terminates.

If the resulting schedule is curative, it is normalized by the process, described in step 2, so that so that the minimal number of viable cancer cells $N_m$ is kept equal to $N_{cur}$, and the algorithm terminates.

Step 5. The resulting multi-dose schedule is compared to the schedule with *maximal non-lethally toxic single dose*, which, injected once, eventually yields $A_{bl}^{cr}$ of toxic decays. This dose is also found by bisection method. If it provides greater overall survival, such single-dose schedule replaces the one found at the previous step. This concludes the work of the optimization algorithm.

#### S.5.3 Multi-dose optimization under parameter sweep

##### S.5.3.1 Comparison of results of single-dose and multi-dose personalized numerical optimization

Using the above discussed algorithm, we performed the personalized multi-dose optimization within the training set of virtual mice. As Fig. S.29 shows, the use of optimized multi-dose schedules allows increasing one-year survival rate by almost 10%, compared to personalized single-dose optimization.

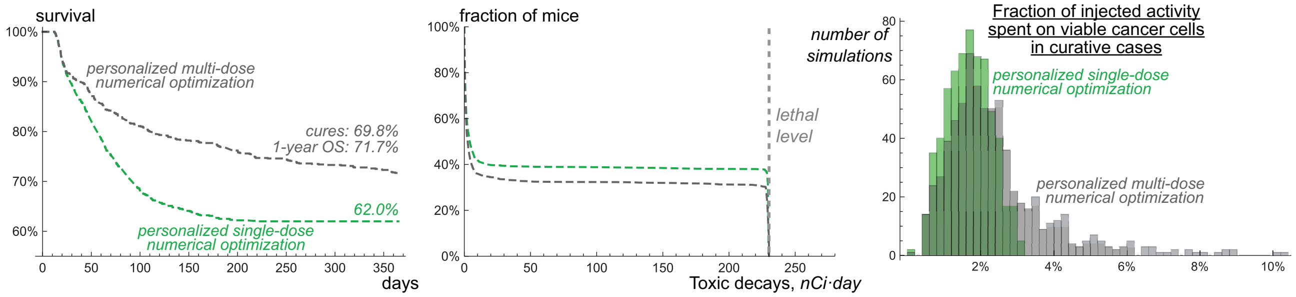


Figure S.29: *Left*, survival curves and *middle*, toxicity curves for personalized numerically optimized single-dose and multi-dose schedules for a single test set of one thousand virtual mice, treated by radioconjugates with coefficient of drug impurity $\eta = 1780$. *Right*, histograms of the fraction of injected activity spent on viable cancer cells, for the corresponding curative cases.

The window in the parameter space for which the use of multi-dose schedules can be beneficial is defined by the ratio of the maximal possible rate of cancer cell damage due to irradiation from nuclides, anchored to them, and cell proliferation rate, $\frac{\alpha\lambda\gamma}{\nu[\eta+1]}/\rho$. This measure lies within the range [1.2-11.9] for the cases where optimized multi-dose schedule yields an increase in overall survival of at least one day compared to the optimal single dose. For virtual mice, corresponding to lower values of this measure, the treatment effectively cannot be optimized, while the mice having its greater values are cured by optimal single doses.

The criteria, derived in section S.5.2.1, correctly identify the majority of virtual mice, that can be cured in the multi-dose setting. Out of all the mice having $C_{SD} > 0$ and $C_{LT} > 0$, only for ≈7.1% and ≈14.3%, respectively, the curative schedules not exceeding the lethal level of toxic decays are not found during numerical optimization. Only seven of the cured mice correspond to both $C_{SD} < 0$ and $C_{LT} < 0$. They represent the above discussed cases, that can be cured by a single dose, but remain effectively incurable in the long term in result of occupation of a significant part of cancer cell receptors by inert antibodies after the first dose.

The middle plot in Fig. S.29 shows that the curative cases are less toxic in multi-dose setting. The key to treatment optimization in curative cases is redistribution of injected radioactivity towards still viable cancer cells. This is evidenced by the right plot in Fig. S.29, which compares the fraction of injected activity spent on viable cancer cells in curative cases for considered single-dose and multi-dose schedules. Notably, for some multi-dose treatments this measure exceeds the corresponding theoretical threshold for a single-dose setting, estimated to be <5% in Section S.2.2.2.

As discussed in Section S.5.1.2, there exist three major factors, that allow redistribution of injected activity towards viable cells. The first factor is the inability of minimal single curative dose to fit onto cancer cells, which is relevant only if cancer by itself can be cured by a single dose without exceeding lethal toxicity. For other cases the remaining two factors come into play, which relevance depends on the relative significance of self-damage, $k_s$. For its low values, when cross-fire plays decisive role, redistribution of activity towards viable cells can be achieved by keeping a considerable fraction of cancer receptors free, but initiating shrinkage of cancer mass with the first dose in order to allow redistribution of further injected radioconjugates over fewer remaining cells. Under high values of $k_s$ multi-dose schedules benefit from maintaining near-saturation levels of cancer binding capacity, so that radioconjugates and deposited dose are redirected to still viable cells, which continue expressing specific receptors. This suggests that optimized treatments under low $k_s$ should aim at undersaturation of cancer binding capacity, while under high $k_s$ they should pursue the proximity of its saturation. This is confirmed by the simulations with optimized multi-dose schedules for virtual mice, which cannot be cured by a single dose without exceeding lethal toxicity. The average fraction of occupied receptors for these cases up to the moments of cure is ≈86% at $k_s < 1/3$; ≈94% at $1/3 < k_s < 2/3$ and ≈95% at $k_s > 2/3$.

##### S.5.3.2 Cancer binding capacity-based multi-dose optimization

The above-described results of numerical optimization are obtained under exact knowledge of all model parameters for each virtual mouse and with knowledge of exact moments of cancer cure. Therefore, they are close to unachievable theoretical limit. From the practical point of view, the crucial question is whether the mentioned optimization techniques can be utilized in a robust way given the uncertainty of physiological parameters in real-life conditions. For the study of this question, we follow our previously introduced assumptions. Namely, we assume that cancer binding capacity $\gamma N_0$ at the moment of drug injection is pre-estimated and is the only known parameter. However, the known physiologic ranges of parameters allow estimating maximal safe doses, that definitely will not result in unacceptable toxicity for any specific parameter set. Also, we assume that the range of used particles and approximate knowledge of cancer cell density allow for rough estimations of relative significance of self-damage and cross-fire. Thus, the virtual mice can be stratified into three groups, corresponding to low, intermediate, and high values of $k_s$.

For these groups the efficacy of the protocols of the following general form were tested. The first dose contained $X \cdot \gamma N_0 + 3$ of antibodies (and $\eta + 1$ times less radioconjugates), or was replaced with maximum safe dose $A_{sf}$, defined by Eq. (S.13), if exceeded it. As estimated previously, the level of toxic decays due to the first dose cannot exceed $A_{bl}^{M}$, defined by Eq. (S.12). The following doses with the amount of antibodies $k \cdot \gamma N_0$ were injected at constant intervals $T$ until their sum was lower than $A_{bl}^{cr} - A_{bl}^{M} \cdot [\lambda + \kappa_c^{min}]/\lambda$. The last dose could be decreased to not exceed this limitation. Thus, it was guaranteed that the total level of toxic decays during treatment could not exceed $A_{bl}^{cr}$ even if the first dose resulted in cure and all the further injected radionuclides that decayed in the bloodstream were attached to intact antibodies rather than to their fragments. The maximum number of doses could not exceed ten, and the treatment duration could not exceed 365 days.

The universal schedules of this form thus depend on three parameters. Initially, they were varied separately in the ranges $X \in [0.7, 3]$, $T \in [3, 40]$ and $k \in [0.01, 1.5]$ for three mice groups, corresponding to $k_s < 1/3$, $1/3 < k_s < 2/3$ and $k_s > 2/3$. It was noted that the approximate ratio of the first dose to the initial cancer binding capacity, $X$, is the major parameter defining the efficacy of the universal schedules, as Fig. S.30 shows. Namely, each group did benefit from initial saturation of cancer cell receptors. For groups with intermediate and high values of $k_s$ this enables redistribution of further injected activity and deposited dose towards viable cells, suggesting that this method can be used in a robust manner.

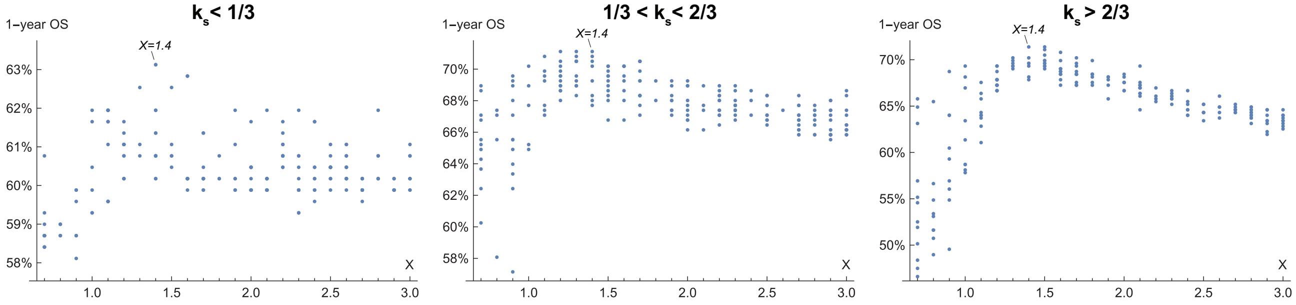


Figure S.30: Dependence of one-year overall survival on $X$, the ratio of the first dose to initial cancer binding capacity, for three groups of virtual mice from the training set, stratified by relative significance of self-damage, $k_s$.

As discussed above, under significant cross-fire irradiation the efficacy of treatment can benefit from initial undersaturation of cancer binding capacity. The left plot in Fig. S.30 suggests that this technique is not robust, since the universal schedules for the group with low values of $k_s$ also benefit from its initial saturation. This is due to the fact that in a large group of different virtual mice initial undersaturation of cancer binding capacity compromises the treatment efficacy for some mice, in particular for those that can be cured by a properly chosen single dose but are incurable in the long term.

In order to reduce the number of free parameters, we further fixed $X$ to be equal to 1.4, since in each of the three groups of mice one of the schedules having the highest one-year overall survival in the performed parameter sweep corresponds to this value of $X$. Using it, we performed the search for optimal universal schedules in the parameter plane $(T, K)$, which results are shown in Fig. S.31. Each of the three groups benefit from relatively small second and subsequent doses, with the optimal inter-dose intervals increasing with the decrease of $k_s$. This is due to the fact that for lower values of $k_s$ the treatment efficacy increasingly depends on cross-fire irradiation. Its efficiency increases along with the shrinkage of cancer mass due do the death of viable cells, which is generally a relatively slow process.

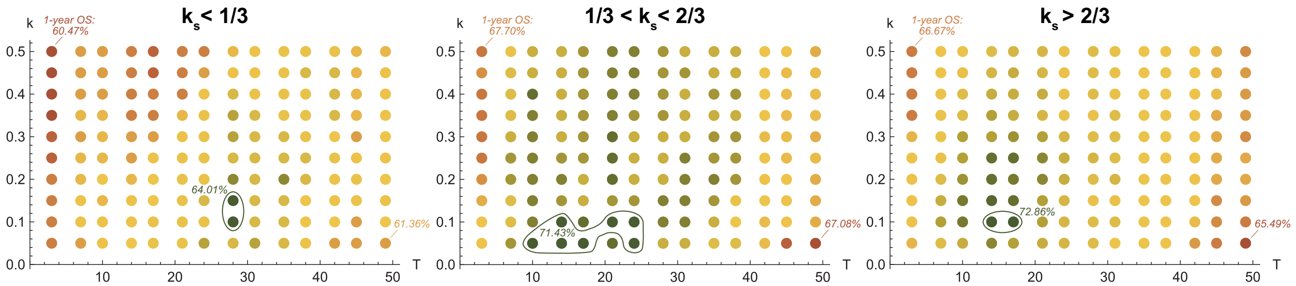


Figure S.31: Dependence of one-year overall survival on inter-dose interval, $T$, and the ratio of second and subsequent doses to initial cancer binding capacity, $k$, under fixed ratio of first dose to cancer binding capacity, $X = 1.4$, for three groups of virtual mice from the training set, stratified by relative significance of self-damage, $k_s$.

The protocols corresponding to $k = 0.1$ and $T = 14$ yield the greatest one-year survival rate for groups with intermediate and high $k_s$. The protocol corresponding to $k = 0.1$ and $T = 28$ is among the two optimal ones for group with low $k_s$. The survival curves for the mentioned protocols are shown in Fig. S.32, where they are compared to the survival curves for previously discussed personalized optimized schedules, obtained under assumption of precise knowledge of all individual parameters.

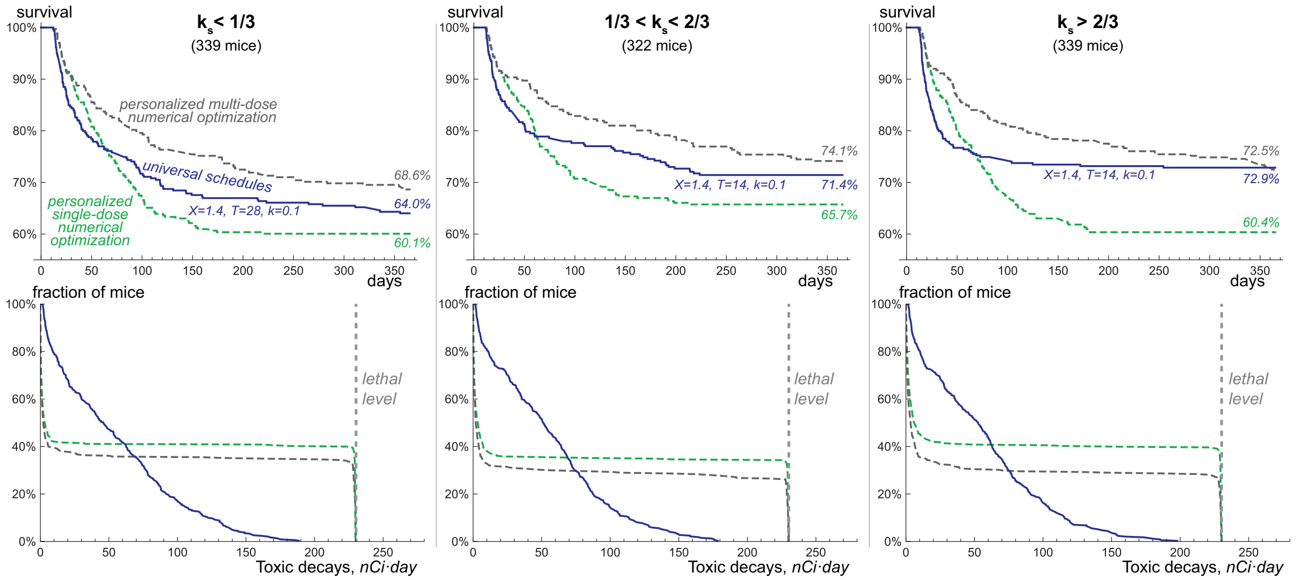


Figure S.32: Survival curves and toxicity curves for personalized numerically optimized single-dose and multi-dose schedules and optimized universal schedules for three groups of virtual mice from the training set, stratified by relative significance of self-damage $k_s$.

Overall, these results confirm that under significant self-damage the redistribution of radioconjugates towards viable cells represents a robust method of increasing the efficacy of targeted radionuclide therapy. Notably, for the group with high $k_s$ the optimized universal protocol exceeds the performance of personalized numerically optimized protocols in terms of one-year survival rate. However, this comes at expense of lower overall survival of non-cured virtual mice. The lower plots of Fig. S.32 show the corresponding toxicity curves, which confirm that the total level of toxic decays during treatment by universal optimized protocols never exceeds the lethal threshold. The verification of universality of chosen protocols is presented in Fig. 5 of main paper, where they are applied to the test set of virtual mice, yielding comparable increase of treatment efficacy.

### References


[1] Megan Minnix, Vikram Adhikarla, Enrico Caserta, Erasmus Poku, Russell Rockne, John E Shively, and Flavia Pichiorri. Comparison of CD38-targeted $\alpha$-versus $\beta$-radionuclide therapy of disseminated multiple myeloma in an animal model. *Journal of Nuclear Medicine*, 62(6):795–801, 2021.

[2] Ryan L Kelly, Tingwan Sun, Tushar Jain, Isabelle Caffry, Yao Yu, Yuan Cao, et al. High throughput cross-interaction measures for human IgG1 antibodies correlate with clearance rates in mice. In *MAbs*, volume 7, pages 770–777. Taylor & Francis, 2015.

[3] Ian D Ferguson, Bonell Patiño-Escobar, Sami T Tuomivaara, Yu-Hsiu T Lin, Matthew A Nix, Kevin K Leung, et al. The surfaceome of multiple myeloma cells suggests potential immunotherapeutic strategies and protein markers of drug resistance. *Nature communications*, 13(1):4121, 2022.

[4] Estefanía García-Guerrero, Ralph Götz, Sören Doose, Markus Sauer, Alfonso Rodríguez-Gil, Thomas Nerreter, et al. Upregulation of CD38 expression on multiple myeloma cells by novel HDAC6 inhibitors is a class effect and augments the efficacy of daratumumab. *Leukemia*, 35(1):201–214, 2021.

[5] Jarosław Wajs and Waldemar Sawicki. The morphology of myeloma cells changes with the progression of the disease. *Contemporary Oncology/Współczesna Onkologia*, 17(3):272–275, 2013.

[6] Xili Liu, Maria Moscvin, Seungeun Oh, Tianzeng Chen, Wonshik Choi, Benjamin Evans, et al. Characterizing dry mass and volume changes in human multiple myeloma cells upon treatment with proteotoxic and genotoxic drugs. *Clinical and Experimental Medicine*, 23(7):3821–3832, 2023.

[7] David A Scheinberg and Michael R McDevitt. Actinium-225 in targeted alpha-particle therapeutic applications. *Current radiopharmaceuticals*, 4(4):306–320, 2011.

[8] Natascia Di Iorgi, Ashley O Mo, Kate Grimm, Tishya AL Wren, Frederick Dorey, and Vicente Gilsanz. Bone acquisition in healthy young females is reciprocally related to marrow adiposity. *The Journal of Clinical Endocrinology & Metabolism*, 95(6):2977–2982, 2010.

[9] Michael C Joiner and Albert J van der Kogel. *Basic clinical radiobiology.* CRC press, 2018.

[10] Sophie Poty, Lynn C Francesconi, Michael R McDevitt, Michael J Morris, and Jason S Lewis. $\alpha$-emitters for radiotherapy: from basic radiochemistry to clinical studies – Part 1. *Journal of Nuclear Medicine*, 59(6):878–884, 2018.

[11] Jayeeta Ghose, Domenico Viola, Cesar Terrazas, Enrico Caserta, Estelle Troadec, Jihane Khalife, et al. Daratumumab induces CD38 internalization and impairs myeloma cell adhesion. *Oncoimmunology*, 7(10):e1486948, 2018.

[12] HE Daldrup-Link, TM Link, EJ Rummeny, C August, S Könemann, H Jürgens, and W Heindel. Assessing permeability alterations of the blood–bone marrow barrier due to total body irradiation: in vivo quantification with contrast enhanced magnetic resonance imaging. *Bone marrow transplantation*, 25(1):71–78, 2000.

[13] Maxim Kuznetsov and Andrey Kolobov. Optimization of antitumor radiotherapy fractionation via mathematical modeling with account of 4 R's of radiobiology. *Journal of Theoretical Biology*, 558:111371, 2023.
